# Chromosomal instability shapes the tumor microenvironment of esophageal adenocarcinoma via a cGAS–chemokine–myeloid axis

**DOI:** 10.1101/2025.05.06.652454

**Authors:** Bruno Beernaert, Rose L Jady-Clark, Parin Shah, Erik Ramon-Gil, Nora M Lawson, Zack D Brodtman, Somnath Tagore, Frederik Stihler, Alfie S Carter, Shannique Clarke, Tong Liu, Winston Zhu, Erkin Erdal, Alistair Easton, Leticia Campo, Molly Browne, Stephen Ash, Nicola Waddell, Thomas Crosby, Simon R Lord, Derek A Mann, Ignacio Melero, Carlos E de Andrea, Andréa E Tijhuis, Floris Foijer, Ester M Hammond, Kadir C Akdemir, Jack Leslie, Benjamin Izar, Eileen E Parkes

**Affiliations:** Department of Oncology, University of Oxford, Oxford, UK; Centre for Immuno-Oncology, Nuffield Department of Medicine, University of Oxford, Oxford, UK; Department of Medicine, Division of Hematology and Medical Oncology, Columbia University Irving Medical Center, Herbert Irving Comprehensive Cancer Center, New York, New York, USA; Faculty of Medical Sciences, Newcastle University, Newcastle, UK; Department of Neurosurgery, Division of Surgery, The University of Texas MD Anderson Cancer Center, Houston, Texas, USA; Translational Histopathology Laboratory, University of Oxford, Oxford, UK; Ludwig Institute for Cancer Research, University of Oxford, Oxford, UK; QIMR Berghofer Medical Research Institute, Brisbane, Australia; Velindre Cancer Centre, Velindre Hospital, Cardiff, UK; Departments of Pathology, Oncology and Immunology, Clínica Universidad de Navarra (CCUN), Pamplona, Spain; CIBERONC, Madrid, Spain; European Research Institute for the Biology of Ageing, University of Groningen, Groningen, The Netherlands

**Keywords:** Chromosomal instability, cGAS, esophagogastric cancer, tumor microenvironment, chemokine, myeloid

## Abstract

Chromosomal instability (CIN), a characteristic feature of esophageal adenocarcinoma (EAC), drives tumor aggressiveness and therapy resistance, presenting an intractable problem in cancer treatment. CIN leads to constitutive stimulation of the innate immune cGAS–STING pathway, which has been typically linked to anti-tumor immunity. However, despite the high CIN burden in EAC, the cGAS– STING pathway remains largely intact.

To address this paradox, we developed novel esophageal cancer models, including a CIN-isogenic model, discovering myeloid-attracting chemokines – with the chemokine *CXCL8* (IL-8) as a prominent hit – as conserved CIN-driven targets in EAC.

Using high-resolution multiplexed immunofluorescence microscopy, we quantified the extent of ongoing cGAS-activating CIN in human EAC tumors by measuring cGAS-positive micronuclei in tumor cells, validated by orthogonal whole-genome sequencing-based CIN metrics. By coupling *in situ* CIN assessment with single-nucleus RNA sequencing and multiplex immunophenotypic profiling, we found tumor cell-intrinsic innate immune activation and intratumoral myeloid cell inflammation as phenotypic consequences of CIN in EAC. Additionally, we identified increased tumor cell-intrinsic *CXCL8* expression in CIN^high^ EAC, accounting for the inflammatory tumor microenvironment.

Using a novel signature of CIN, termed CIN^MN^, which captures ongoing CIN-associated gene expression, we confirm poor patient outcomes in CIN^high^ tumors with signs of aberrantly rewired cGAS–STING pathway signaling.

Together, our findings help explain the counterintuitive maintenance and expression of cGAS–STING pathway components in aggressive, CIN^high^ tumors and emphasize the need to understand the contribution of CIN to the shaping of a pro-tumor immune landscape. Therapeutic strategies aimed at disrupting the cGAS-driven inflammation axis may be instrumental in improving patient outcomes in this aggressive cancer.

## Introduction

Esophageal adenocarcinoma (EAC) is an aggressive and frequently lethal cancer, with 5-year survival rates of less than 20%^1^. Chromosomal instability (CIN) is a defining characteristic of EAC, with 97% of cases reported as belonging to the CIN subtype in the largest study to date of genomic characterization of gastro-esophageal cancers^2^. Manifestations of CIN, including chromosomal alterations, aneuploidies and extrachromosomal DNA, occur early in the development of EAC, arising at non-malignant stages of the precursor lesion Barrett’s Esophagus, and are associated with increased risk of progression to carcinoma^3–6^. Genomic dysregulation from the earliest stages of development of EAC results in an almost ubiquitous mutational inactivation of *TP53* commonly associated with loss of *CDKN2A*^7,8^. Loss of these cell-cycle checkpoints permits accumulation of oncogenic copy number alterations and genomic catastrophes, such as whole-genome doubling and chromothripsis^9,10^, resulting in the chromosomal aberrations and complex genomic rearrangements observed in established EAC.

The chronic inflammatory background on which Barrett’s Esophagus develops likely influences the pattern of immune infiltration subsequently observed in the tumor microenvironment of EAC. Certain chemokines and cytokines, including ligands to the myeloid receptors C-X-C motif Chemokine Receptor 1 and 2 (CXCR1/2; CXCL1, CXCL2, CXCL4, CXCL6 and CXCL8), IL-6 and IL-1β, have been reported to demonstrate a stepwise increase in their expression paralleled with disease progression^11,12^. Additionally, microenvironmental shifts with declining CD8^+^ T cells and enrichment of myeloid and T regulatory cells are associated with progression from Barrett’s Esophagus to EAC^13,14^.

In CIN settings, we and others have reported the chronic stimulation of the innate immune cyclic GMP-AMP synthase (cGAS)-Stimulator of Interferon Genes (STING) pathway^15–18^. Chromosome segregation errors frequently lead to the formation of cytoplasmic structures known as micronuclei (MN) containing whole chromosomes or chromosome fragments that lag behind during anaphase. These mis-segregated chromosomes recruit their own structurally unstable nuclear envelope, which frequently ruptures with resultant exposure of nucleic acids to the cytosol^19^. The double-stranded DNA sensor cGAS detects cytoplasmic DNA, producing the second messenger 2’3’-cGAMP for subsequent activation of STING in an autocrine and paracrine manner. The subsequent chemokine cascade, when acutely activated, results in antitumor immune responses^20^. However, in the chronic setting a paradoxical pro-tumorigenic impact can be observed, with increased IL-6 secretion promoting tumor cell survival^21^, upregulation of the cGAMP hydrolase ecto-nucleotide pyrophosphatase/phosphodiesterase (ENPP1), which results in augmented extracellular adenosine and causes immunosuppression^16^. Furthermore, chronic stimulation results in rewiring of the STING pathway, fostering survival and metastasis via non-canonical NF-κB signaling and ER stress responses^15,17^.

However, the impact of chromosomal instability on the immune landscape in EAC is unknown. We hypothesized that the interaction between cancer and stromal cells, mediated by chronic cGAMP production or cGAS–STING-driven chemokine signaling, may profoundly influence the tumor microenvironment in CIN^high^ cancers and promote tumor aggressiveness while blunting therapeutic efficacy. Here, we provide evidence for CIN causing chronic cGAS–STING activation that enhances the expression of chemokines attracting pro-tumor inflammatory infiltrates. Therefore, understanding the consequences of CIN-driven cGAS stimulation provides novel insights into mechanisms whereby this axis could be disrupted for patient benefit.

## Results

### EAC maintains cGAS–STING despite high burdens of CIN

Given the postulated central role of cGAS–STING in promoting anti-tumor immune responses, some cancers have been reported to suppress cGAS–STING via epigenetic silencing^22,23^. However, analysis of genomic, methylation and transcriptomic data from primary esophageal cancers from The Cancer Genome Atlas (TCGA) demonstrated a maintenance of *cGAS* and *STING* expression across esophageal tumors versus matched normal tissue (**Extended Data Fig. 1a–c**). Broad expression maintenance of *cGAS* and *STING* in esophageal lines was reflected *in vitro*, both within the Cancer Cell Line Encyclopedia (CCLE; **Extended Data Fig. 1d**) as well as across a panel of 10 esophageal cell lines (**Extended Data Fig. 1e**), comprising cell lines derived from non-dysplastic and high-grade dysplastic Barrett’s Esophagus (BE) tissue (CP-A and CP-C, respectively), as well as EAC. Multiplex immunofluorescence (mIF)-based profiling of 24 treatment-naïve EAC biopsy samples confirmed cGAS and STING protein expression in both malignant and stromal cell compartments in EAC, with cGAS expression predominantly in the malignant compartment and STING in the stroma (**Fig. 1a,b**).

**Figure 1.**
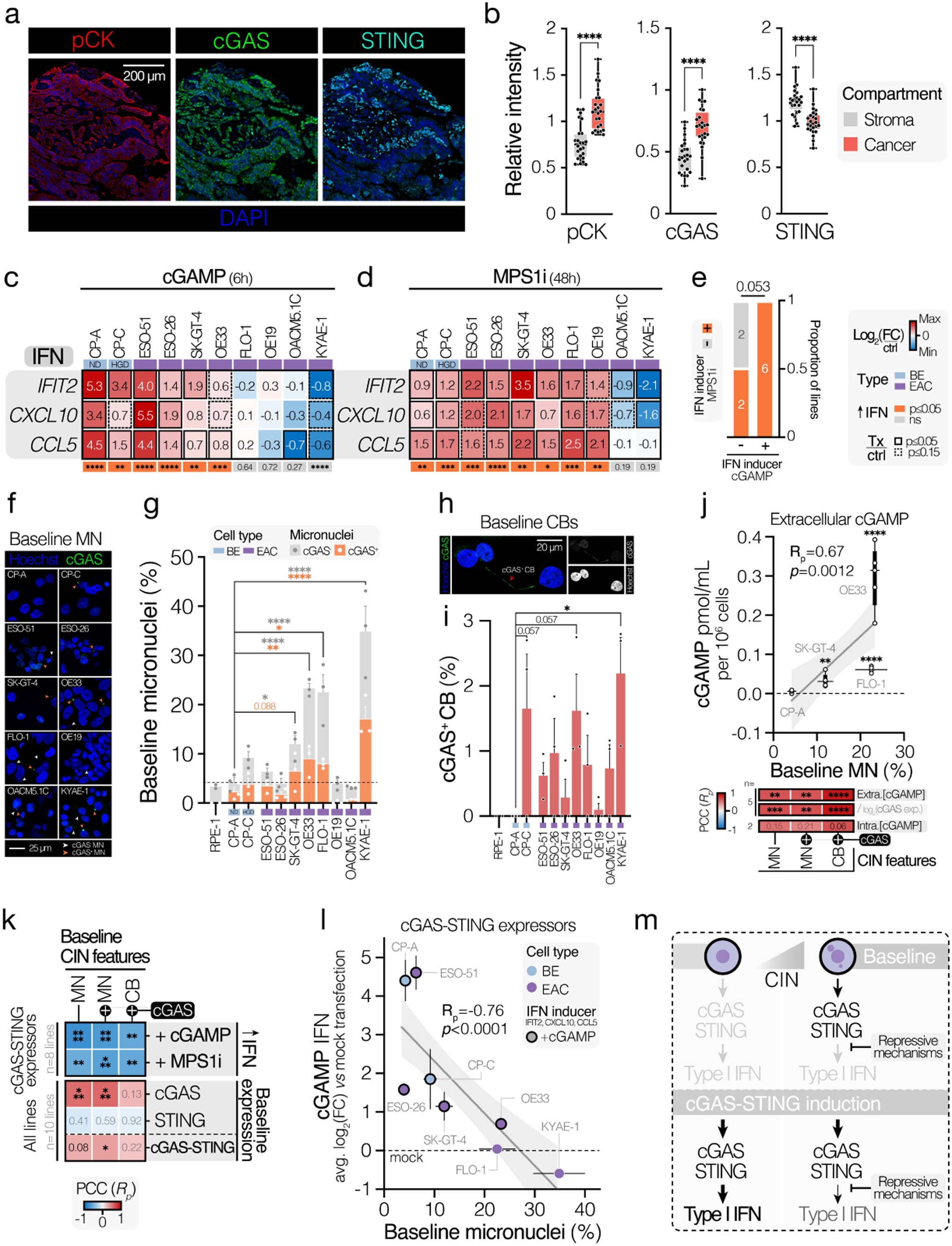
cGAS–STING functionality and CIN are prevalent features of EAC. (**a**) Representative high-resolution image of a human EAC tumor biopsy specimen stained with DAPI (DNA), as well as anti-cGAS, anti-pan-cytokeratin (pCK) and anti-STING antibodies. Scale bar corresponds to 200 μM. (**b**) Quantification of (**a**). *B*ox plots showing the average cancer compartment versus stromal compartment intensities for markers of interest across EAC biopsy specimens. Boxes represent the median of marker intensities ± interquartile range split by cell compartments of interest. Box plot whiskers range from minimum to maximum. Data were analyzed by Wilcoxon matched-pairs signed-rank test, pairing malignant and stromal detections from the same tumors. (**c**,**d**) Heatmap of log_2_-normalized fold-changes (FC) in relative mRNA abundance of the interferon-inducible genes *IFIT2*, *CXCL10* and *CCL5* upon (**c**) 2’3’-cGAMP transfection (2 μg/mL 2’3’-cGAMP + 2 μL/mL lipofectamine-2000, 6h) or (**d**) MPS1i treatment (0.5 μM reversine, 48h). Expression values were obtained through RT-qPCR. Fold-changes (FC) were derived using the 2^-ΔΔCt^ method, normalizing expression values to *18S* expression and respective control treatments. Control treatments: 2 μL/mL lipofectamine-2000 (6h; i.e. ‘mock transfection’) for 2’3’-cGAMP treatments and DMSO (48h) for MPS1i treatments. Color-mapped values represent the average log_2_(FC) of n=3 independent experiments. Data were analyzed by two-sided paired t-test. Significance of overall treatment-induced IFN gene induction was determined via two-sided paired t-test, comparing the pooled condition-level averages of Z-normalized IFN gene expression values of treated versus control samples. Cell lines with significant (p ≤0.05, increased induction) overall treatment-induced IFN-inducible gene induction were considered ‘IFN inducers’. (**e**) Bar graph depicting the relationship between IFN gene induction capacity in response to 2’3’-cGAMP treatment and IFN induction following MPS1 inhibitor treatment in BE and EAC cell lines. Significant inducers of IFN-stimulated genes following treatment are denoted as (+), non-inducers as (-). Significance was tested by two-sided Chi-square (χ^2^) test. (**f**) Representative confocal microscopy images of BE and EAC cell lines under baseline conditions (DMSO-treated, 48h), showing examples of cGAS^-^ MN and cGAS^+^ MN. Cells were stained with anti-cGAS and Hoechst (DNA). Images represent maximum-intensity projections of Z-stacks, taken at 1 μm intervals. Scale bars correspond to 25 μm. (**g**) Quantification of cGAS^-^ and cGAS^+^ micronuclei in BE and EAC lines under baseline conditions. Bars represent the mean ± SEM of n=3, with ∼≥100 cells counted per experiment. Data were analyzed by ANOVA with FDR-based correction. (**h**) Representative confocal microscopy image of baseline (DMSO-treated, 48h) SK-GT-4 cells, showing an example of a cGAS^+^ chromatin bridge. Cells were stained with anti-cGAS and Hoechst (DNA). The image represents a maximum-intensity projection of a Z-stack taken at 1 μm intervals. Scale bar corresponds to 20 μm. (**i**) Quantification of baseline cGAS^+^ chromatin bridge frequencies in EAC and BE cell lines. Bars represent the mean ± SEM of n=3, with ∼≥100 cells counted per experiment. Data were analyzed by ANOVA with FDR-based correction. (**j**) Upper panel: Scatter plot of baseline extracellular 2’3’-cGAMP abundance (quantified by ELISA, normalized to cell number) versus baseline micronucleus frequency (quantified by IF) in select BE and EAC lines. Box plot lines represent the median and error bars represent the range (minimum to maximum) of n=5 independent cGAMP measurements. The bounds of box plots represent the interquartile range (Q1 to Q3). The position of box plots along the x-axis corresponds to the mean MN frequency for each cell line. Horizontal error bars represent ± SEM of n=3 independent IF-based MN estimates. Shown is the estimated simple linear regression line, 95% confidence intervals, Pearson correlation coefficient (R_p_) and the p-value from a Pearson correlation analysis. The significance of differences in extracellular cGAMP concentrations between CP-A and EAC lines (OE33, SK-GT-4, FLO-1) was determined by unpaired two-tailed t-test. Lower panel: Heatmap of Pearson correlation coefficients of correlations between IF-derived CIN feature frequencies and extracellular (Extra.) or intracellular (Intra.) cGAMP abundance across profiled lines. This includes *cGAS* expression-normalized (normalized to log_2_-transformed relative mRNA abundance) extracellular cGAMP abundance, showing an association with CIN feature frequencies irrespective of differences in *cGAS* mRNA expression levels between lines. (**k**) Heatmap of Pearson correlation coefficients between CIN features (cGAS^+^ micronuclei/chromatin bridges and overall micronuclei burden) and baseline mRNA expression of *cGAS* and *STING* or treatment-induced (48h 0.5 μM MPS1i/reversine or 6h 2 μg/mL 2’3’-cGAMP transfection) IFN gene induction. Significance corresponds to Pearson correlation analysis p-values. (**l**) Scatter plot of baseline MN burden versus average log_2_-transformed IFN-stimulated gene fold-change (of *IFIT2*, *CXCL10* and *CCL5*) upon 2’3’-cGAMP stimulation (6h, 2 μg/mL; versus mock transfection) across EAC and BE lines with meaningful cGAS and STING expression levels (i.e. cGAS–STING expressors). Shown are the estimated simple linear regression line, 95% confidence intervals, Pearson correlation coefficient (R_p_) and the p-value from a Pearson correlation analysis. Horizontal and vertical error bars represent the ± SEM from n=3 independent IF and qPCR experiments, respectively. (**m**) Model of the relationship between cGAS–STING pathway function and inherent cellular CIN in esophageal cells. CIN^low^ cells exhibit normal cGAS–STING pathway functionality and can drive potent type I IFN responses upon treatment-induced activation of the cGAS–STING pathway. CIN^high^ cells suffer from upstream cGAS–STING activation under baseline conditions owing to their high burden of cytosolic DNA. Repressive mechanisms are, thus, put in place to avoid engagement of the downstream antiproliferative type I IFN cascade. As a result, CIN^high^ cells are less poised towards IFN I engagement upon further exacerbation of cGAS–STING activation, owing to the pre-existing presence of downstream repressive mechanisms. **** p ≤0.0001; *** p ≤0.001; ** p ≤0.01; * p ≤0.05; ns, not significant.

To assess whether the maintenance of cGAS–STING expression in BE and EAC lines corresponded to functional type I IFN signaling upon STING pathway induction, we treated cells with the STING agonist 2’3’-cGAMP or induced CIN using the MPS1 inhibitor (MPS1i) reversine (**Extended Data Fig. 2a,b**) and monitored the expression of the interferon-stimulated genes (ISGs) *CCL5*, *CXCL10*, and *IFIT2 –* key targets of STING-driven type I IFN signaling (**Fig. 1c,d**). Non-transformed, genomically stable BE lines (CP-A and CP-C) exhibited strong induction of IFN-responsive genes (referred to as ‘IFN’ genes) in response to STING-activating treatments, whereas EAC lines exhibited more variable responses, including an expected absence of 2’3’-cGAMP-driven IFN gene induction in cell lines with suppression of core cGAS–STING machinery expression (OE19 and OACM5.1C; **Extended Data Fig. 1e**). Some EAC lines also demonstrated failure of IFN gene induction in response to 2’3’-cGAMP in lines with competent cGAS–STING expression levels (FLO-1, KYAE-1). Notably, some cell lines that did not demonstrate IFN gene expression in response to direct STING agonism through 2’3’-cGAMP treatment maintained competent IFN signaling in response to MPS1 inhibition (e.g. OE19 and FLO-1), suggestive of STING-independent modes of CIN-driven IFN gene activation. Nevertheless, IFN induction capacity in response to cGAMP exposure coincided with IFN upregulation following MPS1i, suggesting a relationship between STING activation-driven and CIN-driven IFN induction capacity (**Fig. 1e**).

Constitutive CIN has recently been proposed to drive the adaptive rewiring of signaling downstream of STING, enabling tumor cells to evade the deleterious effects of type I IFN activation^17^. To assess the prevalence and extent of CIN in EAC cells and evaluate its relationship with IFN gene induction capacity, we next measured the frequency of proposed cGAS-activating CIN features, including micronuclei (MN), cGAS^+^ MN and cGAS^+^ chromatin bridges (CBs), under baseline conditions by immunofluorescence (**Fig. 1f–i**).

cGAS^+^ MN and chromatin bridges were observed to be similarly low in both the non-dysplastic BE CP-A line and the genomically stable, non-cancerous hTERT RPE-1 cell line. However, CIN feature frequencies differed significantly across high-grade dysplastic BE as well as EAC lines, with 4 out of 8 EAC lines (OE33, SK-GT-4, FLO-1 and KYAE-1) exhibiting significantly elevated MN or chromatin bridge burdens over the CP-A cell line. Baseline incidences of putatively cGAS-activating CIN features were found to be significantly intercorrelated (**Extended Data Fig. 2c**). CIN features also correlated with tonic cGAS activity, as inferred through 2’3’-cGAMP ELISA, irrespective of *cGAS* expression, across profiled cell lines (**Fig. 1j**). In contrast, baseline CIN feature burdens were strongly anti-correlated with IFN gene induction capacity following either 2’3’-cGAMP or MPS1i stimulation, indicating diminished interferon pathway engagement in response to stimulation of cGAS–STING in CIN^high^ cells (**Fig. 1k,l**).

Taken together, these results suggest that EAC cells broadly maintain cGAS and STING expression and retain at least some degree of cGAS–STING–interferon pathway functionality. However, the magnitude of effective interferon pathway response upon cGAS–STING stimulation is inversely proportional to the inherent CIN burden to which tumor cells are subjected *in vitro* (**Fig. 1m**).

### CIN-driven cGAS–STING controls a distinct subset of immune target genes in EAC

To interrogate the nature of CIN-induced cGAS–STING innate immune responses in EAC cells, we abrogated cGAS using CRISPR-Cas9-mediated knockout (cGAS^KO^) in a subset of EAC lines, selected specifically for their high inherent micronucleation rates and differential IFN I induction capacities (OE33, FLO-1 and SK-GT-4). cGAS^KO^ clones showed a loss of cGAS at the protein level as well as a significantly diminished cGAS mRNA abundance (**Extended Data Fig. 3a–c**), and failed to produce 2’3’-cGAMP in response to both CIN induction through MPS1 inhibition and direct cGAS agonism through G_3_-YSD (a Y-form DNA selective cGAS agonist^24^) transfection, compared to respective empty-vector (Cas9) clones (**Extended Data Fig. 3d**).

To identify cGAS-dependent CIN-induced targets, we profiled the transcriptomes of Cas9 control and cGAS^KO^ clones of OE33 and SK-GT-4 cells, with or without MPS1i treatment (**Fig. 2a**). A substantial proportion of CIN-upregulated genes in both cell lines were found to be dependent on cGAS function, with 144 (23.3%) and 255 (41.8%) genes in OE33 and SK-GT-4 cells, respectively, exhibiting a significantly diminished upregulation in cGAS^KO^ backgrounds (**Fig. 2b–d**; **Extended Data Fig. 3e–h**). Of these, approximately 1 in 5 had known immune or inflammatory functions. Eighteen consensus hits were identified, including the CXCR1/2-acting chemokines *CXCL8* (encoding IL-8) and *CXCL2* (also known as MIP2-alpha), the interferon pathway genes *IRF1* and *IFIT2,* as well as components of the unfolded protein response (UPR) endoplasmic reticulum (ER) stress and autophagy pathways (e.g. *CHAC1, ZFAND2A, TRIM16* and *MAP1LC3B* [known as LC3]), potentially reflecting reported non-canonical roles for cGAS in these pathways^25,26^ (**Fig. 2e**). Overrepresentation analysis indicated that CIN-induced cGAS-dependent genes in both OE33 and SK-GT-4 were enriched for immune-related functions, including interferon and chemokine/cytokine signaling pathways (**Fig. 2f**). Gene set enrichment analysis (GSEA) of MPS1i-induced transcriptional changes revealed that the engagement of inflammatory response pathways (including NF-κB, chemokine, cytokine and inflammatory signaling gene sets), rather than interferon pathways (including interferon, ISG and JAK-STAT gene sets) was most pronounced upon CIN induction and most markedly diminished by cGAS deletion in both cell lines (**Fig. 2g**).

**Figure 2.**
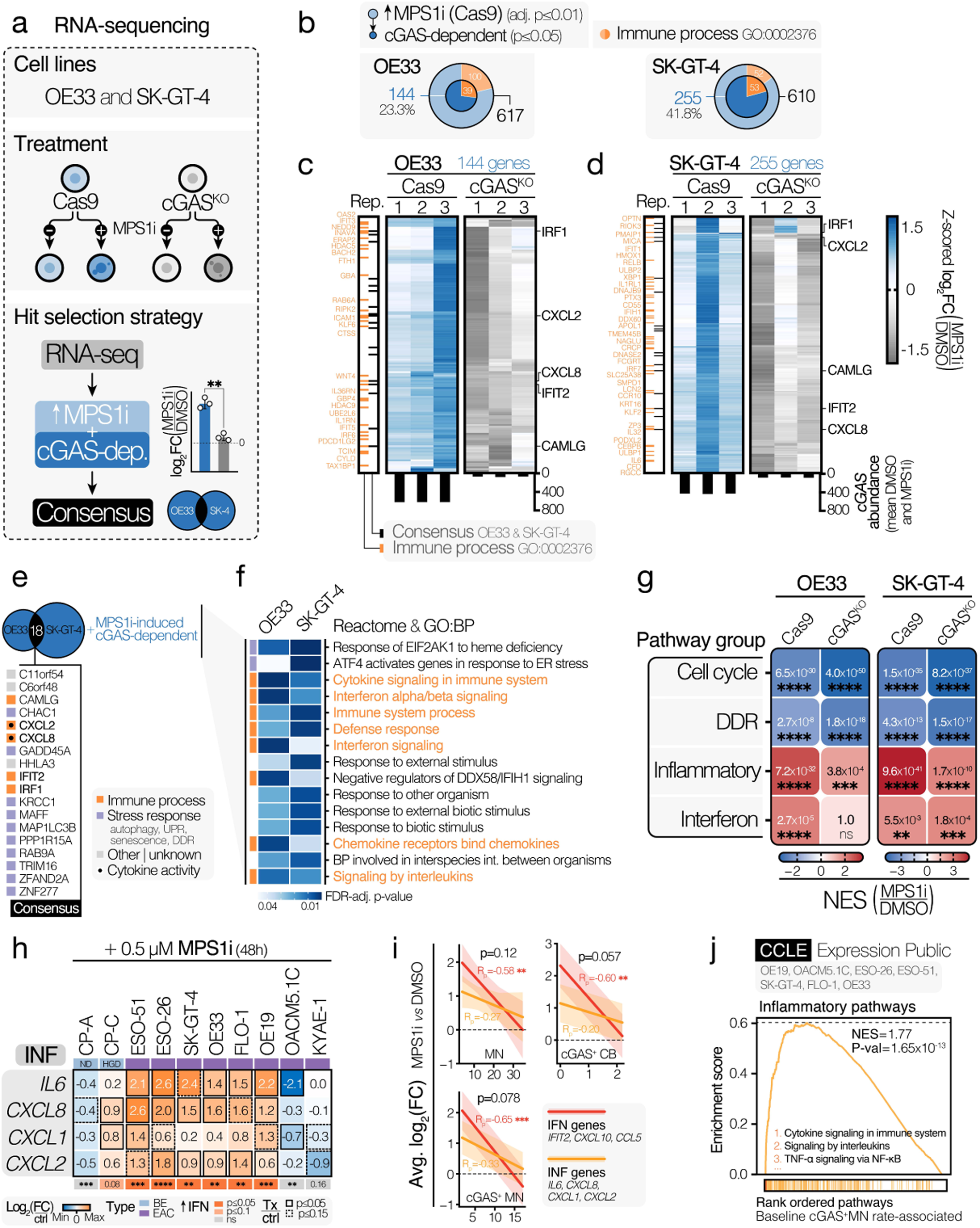
CIN-driven cGAS–STING activation drives inflammatory gene expression in EAC cells. (**a**) RNA-sequencing strategy to identify cGAS-dependent CIN (i.e. MPS1i)-induced genes in OE33 and SK-GT-4 cells. CGAS-proficient Cas9 cells and cGAS^KO^ cells were exposed to either DMSO or the MPS1 inhibitor reversine (0.5 μM) for 48h before sample collection. Genes that were significantly upregulated (adj. p ≤ 0.01) upon MPS1-inhibition in cGAS-proficient cells were considered MPS1i-induced. Genes that were significantly more strongly induced in Cas9 proficient over cGAS^KO^ cells were considered CIN-driven cGAS-dependent hits. Performing this analysis in two independent lines allowed for the identification of consensus hits between the two lines. (**b**) Venn diagram of identified CIN/MPS1i-upregulated (outer circle) and CIN-induced cGAS-dependent (inner circle) hits for OE33 and SK-GT-4 cells. The proportion of each subset of hits annotated with the ‘Immune System Process’ gene ontology (GO) term (GO:Biological Process) has been indicated. (**c**, **d**) Heatmaps of log_2_-transformed fold-changes (log_2_[FC]; MPS1i vs DMSO, RNA-seq) of cGAS-dependent CIN-driven genes for (**c**) OE33 and (**d**) SK-GT-4 Cas9 and cGAS^KO^ cells across n=3 biological repeats. Genes belonging to the ‘Immune System Process’ GO term have been highlighted in orange. Consensus cGAS-dependent CIN-driven hits annotated with ‘Immune System Process’ are shown in black. Average cGAS expression levels (from RNA-seq) across DMSO- and MPS1i-treated samples are plotted at the bottom of heatmaps. Color maps to row-wise Z-scored log_2_(FC). (**e**) Venn diagram showing the overlap between all detected MPS1i-induced cGAS-dependent genes in OE33 cells and SK-GT-4 cells. All 18 consensus hits are shown and are annotated with associated biological functions. (**f**) Heatmap of overrepresentation analyses (using GO: Biological Process and Reactome gene set collections) of MPS1i-induced cGAS-dependent genes in OE33 cells and SK-GT-4 cells. The top 15 shared overrepresented pathways are shown and ranked from top to bottom by cumulative significance across the two lines. Pathways highlighted in orange relate to immune-promoting functions. (**g**) Heatmap of gene set enrichment analysis (GSEA) outputs showing MPS1i (reversine)-induced pathway changes in OE33 and SK-GT-4 Cas9 and cGAS^KO^ clones, with a focus on overall enrichment in pathways relating to the cell cycle, DNA damage response (DDR), inflammatory response or interferon signaling. Values within cells represent the significance of positive or negative enrichment (FDRq) for a given pathway group upon MPS1i treatment. Color maps to normalized enrichment score (NES). (**h**) Heatmap of log_2_-normalized fold-changes (FC) in relative mRNA abundance of the inflammatory (INF) targets *IL6*, *CXCL8*, *CXCL1* and *CXCL2* upon MPS1i treatment (0.5 μM reversine, 48h) versus DMSO treatment across BE and EAC cell lines. Expression values were determined by RT-qPCR. Fold-changes were derived using the 2^-ΔΔCt^ method, normalizing expression values to *18S* expression and DMSO control treatment gene expression. Color-mapped values represent the average log_2_(FC) of n=3 independent experiments. Data were analyzed by two-sided paired t-test. Significance of overall treatment-induced INF gene induction was determined via two-sided paired t-test, comparing the pooled condition-level averages of Z-normalized INF gene expression values of treated versus control samples. Cell lines with significant (p ≤0.05, increased induction) overall reversine-induced INF gene induction were considered ‘INF gene inducers’. (**i**) Scatter plots showing the relationship between baseline CIN feature (cGAS^+^ MN and chromatin bridges and all micronuclei) frequencies and IFN-inducible gene (‘IFN’; IFIT2, CXCL10, CCL5) or inflammatory target (‘INF’; *IL6*, *CXCL8*, *CXCL1* and *CXCL2*) induction upon MPS1i treatment (0.5 μM reversine, 48h) across cGAS–STING expressing esophageal cell lines. The simple linear regression line, 95% confidence intervals, Pearson correlation coefficients (R_p_), Pearson p-values and linear model interaction term p-values are shown. (**j**) GSEA enrichment plot showing enrichment of inflammatory pathway gene expression with increasing baseline cGAS^+^ MN rates across n=7 indicated EAC cell lines. **** p ≤0.0001; *** p ≤0.001; ** p ≤0.01; * p ≤0.05; ns, not significant.

Several cGAS-dependent immune hits (*IL6, CXCL8, CXCL2* and *IFIT2*) were confirmed using RT-qPCR, demonstrating cGAS-dependent CIN-induced upregulation across multiple independent clones (**Extended Data Fig. 3i**). Similarly, MPSi-induced secretion of IL-8 was measured by ELISA in cell supernatants and confirmed to be significantly reduced in cGAS^KO^ cell lines (**Extended Data Fig. 3j**).

CIN-induced expression upregulation of inflammation-associated cytokines (*IL6*, *CXCL8*, *CXCL1* and *CXCL2;* ‘INF’ genes) appeared broadly conserved across BE and EAC lines (in 7 of 10 lines; **Fig. 2h**) and did not exhibit the same degree of baseline CIN-associated suppression seen with IFN-responsive genes (**Fig. 2i**). Indeed, baseline inflammatory pathway activity (inferred from CCLE RNA-sequencing data) was commensurate with the inherent cGAS^+^ MN burdens of EAC lines, as determined via IF-based profiling (**Fig. 2j**), suggestive of tonic CIN-driven inflammatory activity in cell lines with high inherent micronucleation rates.

Nevertheless, CIN-induced inflammatory gene induction was still noted in the cGAS non-expressing line OE19 (**Extended Data Fig. 1d**), and in the STING pathway-impaired FLO-1 cell line (**Fig. 1c**) upon cGAS abrogation (**Extended Data Fig. 3k**), indicative of cGAS–STING-independent modes of CIN-driven inflammatory target induction.

Collectively, our data support a model wherein CIN results in the constitutive expression of cGAS-dependent inflammatory mediators, although some inflammatory signaling is retained in certain CIN^high^ settings independent of cGAS status.

### CIN scales with inflammatory activity in an isogenic Barrett’s Esophagus model

To ensure our findings were a direct result of CIN and not due to the impact of drug-induced acute induction of CIN on cell viability, and to more closely model the chronicity of CIN in EAC, we developed novel isogenic esophageal cell line models with varying levels of steady-state CIN (**Extended Data Fig. 4a–e**). Using the non-dysplastic, h-TERT-immortalized Barrett’s Esophagus-derived CP-A cell line^27^ as a founder model, we validated its reported *TP53* wildtype status and confirmed possession of a functional cGAS–STING pathway (**Extended Data Fig. 5a–e**). To mimic a genetic background permissive to high levels of CIN in this cell line, we first serially inactivated *TP53* and *CDKN2A* (p53^KO^ and p16^KO^, respectively) using CRISPR-Cas9 mediated gene editing (**Extended Data Fig. 5f**). Disruption of these pathways is among the most prevalent genetic events in EAC tumors and are typically early events in disease progression^7^. Subsequently, we introduced a dominant-negative mutant form of the mitotic regulator MCAK (known as KIF2C; dnMCAK) on the p53^KO^ background, to induce the high rate of chromosome mis-segregations encountered in EAC (**Fig. 3a**). Single cell clones were selected using fluorescence-activated cell sorting (FACS), puromycin selection and IF-based screening of baseline cGAS^+^ MN frequencies across clones (**Extended Data Fig. 4a–e**). Selected clones were validated through amplicon sequencing of gRNA target sites, challenge with Nutlin-3 and puromycin, and immunoblotting for p53 and MCAK (**Extended Data Fig. 5f–n**).

**Figure 3.**
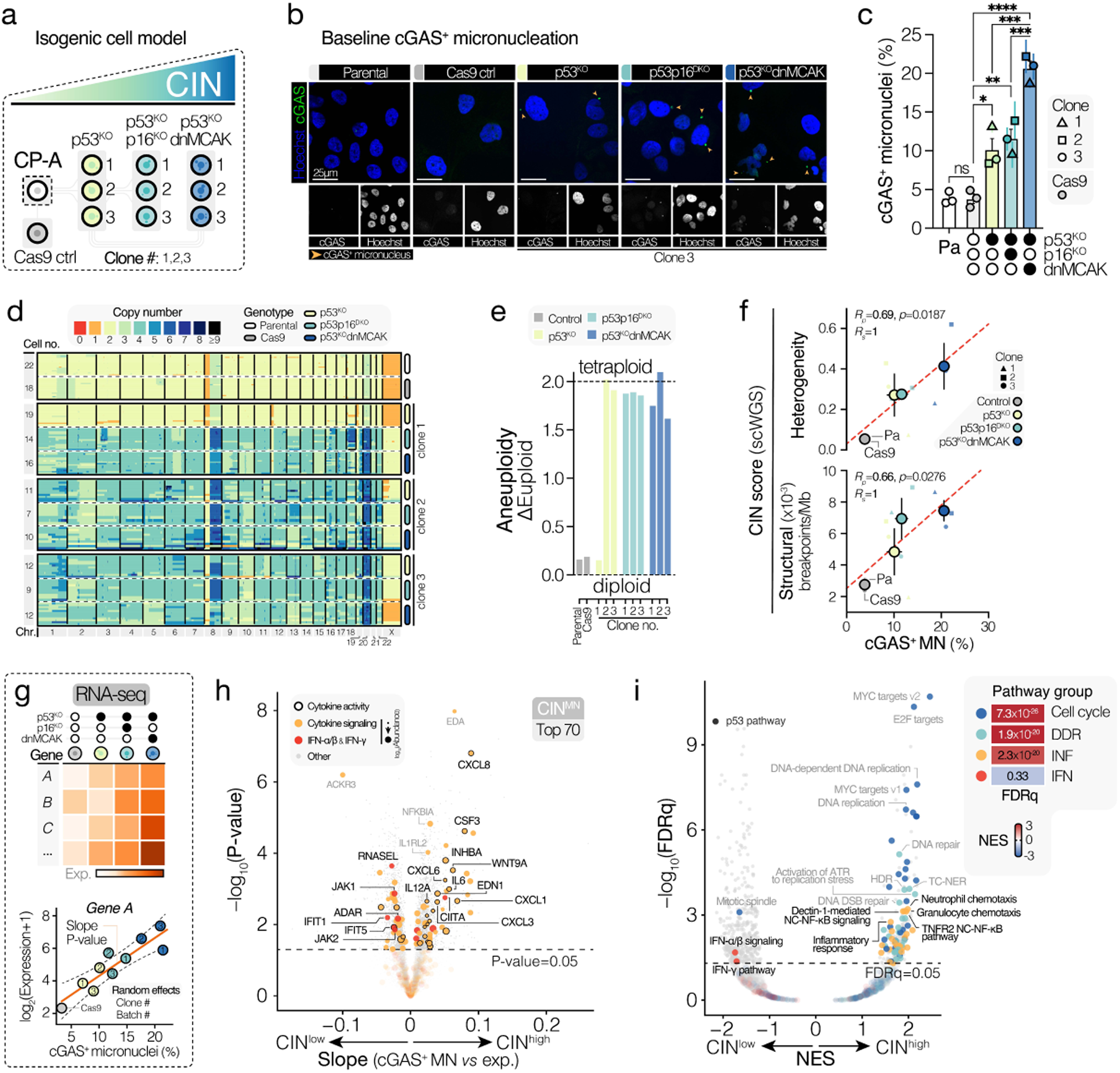
Inflammatory pathway activity scales with inherent level of CIN in an isogenic Barrett’s esophagus cell line model. (**a**) Schematic representation of the isogenic Barrett’s esophagus (BE) CIN line model, showing the genotypes of altered cell lines derived from the parental (dashed box) non-dysplastic BE founder cell line, CP-A. (**b**) Representative confocal microscopy images showing the baseline incidence of cGAS^+^ micronuclei across the cell line models. Images of p53^KO^, p53p16^DKO^, p53^KO^dnMCAK cells all correspond to cells derived from the same progenitor (clone 3). Cells were stained with anti-cGAS and Hoechst (DNA). The image represents a maximum-intensity projection of a Z-stack taken at 1 μm intervals. The scale bar corresponds to 25 μm. (**c**) Quantification of cGAS^+^ micronuclei in parental CP-A cells and derived altered (Cas9 control, p53^KO^, p53p16^DKO^, p53^KO^dnMCAK cell lines under baseline conditions. Bars for p53^KO^, p53p16^DKO^, p53^KO^dnMCAK genotypes represent the mean of n=3 independent clones ± SEM, where each point represents the mean ± SEM of n=3 independent experiments. Bars for parental and Cas9 cells represent the mean ± SEM of n=3 independent experiments. Datapoint shapes correspond to clone numbers for p53^KO^, p53p16^DKO^, p53^KO^dnMCAK lines. Circa ≥ 100 cells were counted per independent experiment. Data were analyzed by ANOVA with FDR-based correction. (**d**) Heatmap of single-cell whole-genome sequencing (scWGS) results. Each row represents a single cell of the indicated genotype. Cell numbers sequenced per genotype are displayed in the left-hand column. Columns represent autosomes 1–22 (left to right) and the X chromosome. Color maps to inferred ploidy at the indicated locus. (**e**) Bar plots of inferred aneuploidy scores from scWGS (relative to a euploid control), showing parental and Cas9 CP-A cells are near-euploid and altered clones (p53^KO^, p53p16^DKO^, p53^KO^dnMCAK) are predominantly near-tetraploid. (**f**) Scatter plots of genotype-level average cGAS^+^ MN burdens (at baseline) versus average scWGS-derived CIN metrics, including the inferred number of structural breakpoints per megabase (Mb; upper panel) and between-cell karyotype heterogeneity (lower panel). The estimated simple linear regression line, Pearson correlation coefficient (R_p_), Spearman correlation coefficient (R_s_) and the Pearson correlation analysis p-value are shown, showing a significant positive association between genotype-level cGAS^+^ MN burdens and scWGS inferred CIN metrics. Datapoint shapes for p53^KO^, p53p16^DKO^, p53^KO^dnMCAK correspond to clone number. Error bars along the x-axis correspond to ± SEM of genotype-pooled average cGAS^+^ MN burdens. Error bars along the y-axis correspond to ± SEM of genotype-pooled average scWGS-derived CIN metrics. (**g**) Experimental strategy for the detection of CIN-associated genes from RNA-sequencing data. A linear mixed-effects model was used iteratively to assess the relationship between observed baseline cGAS^+^ MN frequencies (by immunofluorescence) and the expression level of every gene detected by RNA-sequencing, whilst accounting for confounding variables (random effects), such as the clone number and batch in which samples were collected. This allowed an evaluation of which genes and biological pathways are most strongly upregulated in cells with high baseline micronucleus frequencies. (**h**) Volcano plot of linear mixed-effects model outputs assessing the linear relationship between inherent cGAS^+^ MN burden and expression level of a gene in the CP-A cell line model. ANOVA p-value (y-axis) indicates the statistical significance of the association between expression and cGAS^+^ MN burden. Slope (x-axis) indicates the magnitude change in gene expression per unit increase in cGAS^+^ MN burden. Genes annotated with the ‘Cytokine signaling’ gene ontology term (GO:0019221) are highlighted in orange. Genes with known IFN-α/β or IFN-γ pathway (R-HSA-909733 or R-HSA-877300, respectively) involvement are highlighted in red. Genes with reported cytokine activity (GO:0005125) are marked by a border. Top ranking genes with immune pathway involvement are labelled. Dot size in immune pathway genes maps to the (log_10_-transformed) mRNA abundance in the RNA-seq experiment across all samples. The top 70 most significantly positively associated genes were used to derive an esophageal cell-specific transcriptional signature of CIN, termed CIN^MN^, used in downstream analyses. (**i**) Volcano plot of gene set enrichment analysis (GSEA) outputs showing pathways most strongly associated with cGAS^+^ MN burden in CP-A cells. Select pathways have been categorized based on their broad biological function into one of four major pathway groups: cell cycle-related pathways, DNA damage response (DDR)-related pathway, inflammatory response pathways (INF) and interferon response pathways (IFN). The p53 pathway (from the MsigDB Hallmarks collection) has been highlighted given its strong negative association with inherent cGAS^+^ MN burden. The heatmap in the top right corner shows the degree of enrichment of broad pathway groups across all enriched pathways. **** p ≤0.0001; *** p ≤0.001; ** p ≤0.01; * p ≤0.05; ns, not significant.

To evaluate the extent of cGAS-activating CIN in our isogenic cell line model, we quantified the baseline frequency of cGAS^+^ MN using IF. Deletion of *TP53* led to a significant increase in cGAS^+^ MN, which was further amplified by dnMCAK overexpression, with strong concordance between independent clones (**Fig. 3b,c**). Additional deletion of *CDKN2A* (p53p16^DKO^) did not augment the observed CIN phenotype compared to p53^KO^ CP-A cells. Using shallow single-cell whole genome sequencing (scWGS)^28^, CP-A parental and Cas9 cells were confirmed to be largely karyotypically homogenous and near-diploid (**Fig. 3d**). Whole-genome doubling was observed in 2 of 3 p53^KO^ clones and all successively derived clones (p53p16^DKO^ and p53^KO^dnMCAK clones; **Fig. 3e**). Elevated rates of CIN were reflected at the genomic level, with altered lines being increasingly characterized by genomic aberrations associated with chromosome mis-segregations, including an elevated frequency of inferred chromosomal breakpoints and increased karyotype heterogeneity across sequenced cells, scaling in a similar manner to observed cGAS^+^ MN rates (**Fig. 3f**).

We performed RNA-sequencing (RNA-seq) profiling of all cell lines of our CIN-isogenic model (Cas9, p53^KO^, p53p16^DKO^, p53^KO^dnMCAK) to assess the impact of stably elevated CIN on transcriptomic profiles, using a linear mixed-effects model to assess the relationship between gene expression and the observed cGAS^+^ MN frequency of each cell line (**Fig. 3g**). Similarly to MPS1i-treatment induced changes, we again observed a significant scaling of expression of multiple inflammatory response-related cytokine and chemokine genes with baseline cGAS^+^ MN burden, including CXCR1/2 ligands (*CXCL8, CXCL1, CXCL3, CXCL6, CXCL7*) as well as *IL6, CSF1* and *CSF3* (**Fig. 3h**). Expression profiling of several CIN-associated chemokines (*CXCL8, IL6* and *CXCL1*) through RT-qPCR confirmed a proportional increase with CIN burden, with a similar increase noted in IL-8 secretion as measured by ELISA in cell supernatants (**Extended Data Fig. 6a–c**). At a pathway level, gene set enrichment analysis (GSEA) showed broad enrichment of multiple inflammation-related pathways with increasing CIN, including neutrophil and granulocyte chemotaxis, cytokine-cytokine receptor interaction and non-canonical NF-κB (**Fig. 3i**). Activation of non-canonical NF-κB signaling was also observed in CP-A p53^KO^dnMCAK cells, with an increased p52 protein abundance and p52/p100 ratio compared to parental CP-A cells (**Extended Data Fig. 6d,e**).

To assess the degree to which *in vitro* transcriptional outcomes of CIN in CP-A cells reflected CIN as encountered in other cells and in EAC tumors, we selected the top 70 genes most strongly positively associated with cGAS^+^ MN burden (termed the CIN^MN^ signature; **Supplementary Table 1**) and evaluated the association between our *in vitro* model-derived CIN^MN^ signature scores and previously published orthogonal measures of CIN, including the transcriptional CIN70 signature score^29^ (referred to as CIN^70^) and the WGS-derived Aneuploidy score^30^. Using RNA-seq data from the CCLE, we identified a positive correlation between transcriptome-derived CIN^MN^ scores and baseline cGAS^+^ MN burdens across 7 EAC cell lines (**Extended Data Fig. 7a**). Additionally, across cell lines derived from multiple tumor sites in CCLE as well as in the EAC and the closely related chromosomally unstable stomach adenocarcinoma (STAD-CIN) TCGA cohorts, CIN^MN^ scores significantly positively associated with CIN^70^ and Aneuploidy scores (**Extended Data Fig. 7b–d**).

To assess whether observed correlations between CIN scores also reflected a similar underlying biology in CIN^high^ EAC tumors, we used GSEA to determine which pathways scaled with CIN^MN^, CIN^70^ and Aneuploidy scores. All three CIN scores were associated with similar biological processes, including expression of E2F targets, MYC targets and G_2_/M checkpoints (**Extended Data Fig. 7e**). In keeping with our *in vitro* findings, CIN measures were consistently associated with increased expression of CXCR1/2 ligands. The inclusion of cGAS–STING expression scores as a covariate in our model enabled evaluation of genes that scaled with CIN in a manner dependent on cGAS–STING expression levels. Multiple chemokines and cytokines were found to correlate with CIN^MN^ in a cGAS– STING-dependent manner, including *CXCL8*, *IL6* and *CSF3* (**Extended Data Fig. 7f**), accompanied by a broad enrichment in cytokine-related pathways, consistent with our *in vitro* modelling (**Extended Data Fig. 7g**). Together, these data are consistent with CIN as a driver of a distinct inflammatory response within the tumor microenvironment.

### Micronucleus burden in EAC tumors correlates with innate immune activity

To define the role of elevated CIN in tumor cell-intrinsic innate immune activation and microenvironmental inflammation, we performed single-nucleus transcriptomic analysis of human EAC tumors across the gradient of CIN. Here, we first refined a mIF-based approach to detect cGAS^+^ MN as a measure of ongoing cGAS-activating CIN, based on our previous methodology^16^.

Formalin-fixed paraffin-embedded (FFPE) tumor sections were stained using a mIF panel consisting of antibodies detecting cGAS and pan-cytokeratin (pCK) alongside the nuclear counterstain DAPI, enabling the identification of cGAS^+^ MN in malignant EAC cells (**Extended Data Fig. 8a**). We quantified cGAS^+^ micronuclei within the cancer cell compartment, normalizing to tumor cell number per section using a semi-automated method that closely matched manual (‘ground-truth’) counts (**Extended Data Fig. 8b**). Consistent with CIN as a hallmark of malignant cells, cGAS^+^ MN were largely confined to tumor cell compartments versus stromal areas (**Extended Data Fig. 8c**). In addition, cGAS^+^ MN scores showed high reproducibility across patient-matched biopsy and resection samples, with lower intra-patient variance compared to inter-patient differences **(Extended Data Fig. 8d)**.

Next, we conducted whole-genome sequencing (WGS) on 12 matched tumor samples to quantify the extent of CIN inferred from genomic features and compare it to cGAS^+^ MN scores. We observed striking correlations between cGAS^+^ MN frequencies and WGS-derived measures of CIN, including the cumulative load of all structural variants (SVs), as well as specific SVs, including inversions, break-ends, duplications and deletions (**Extended Data Fig. 8e,f**).

Having validated our *in situ* CIN scoring approach, we shortlisted 9 tumors spanning the CIN spectrum observed across 45 primary EAC tumors for single-nucleus RNA-sequencing (snRNA-seq; **Extended Data Fig. 9a,b**). SnRNA-seq was performed on FFPE sample-matched fresh frozen (FF) tumor samples using the 10X Genomics platform, sampling 4 CIN^low^ and 5 CIN^high^ tumors, with CIN^high^ tumors harboring, on average, 2.7 times the cGAS^+^ MN burden of CIN^low^ tumors (**Fig. 4a,b**).

**Figure 4.**
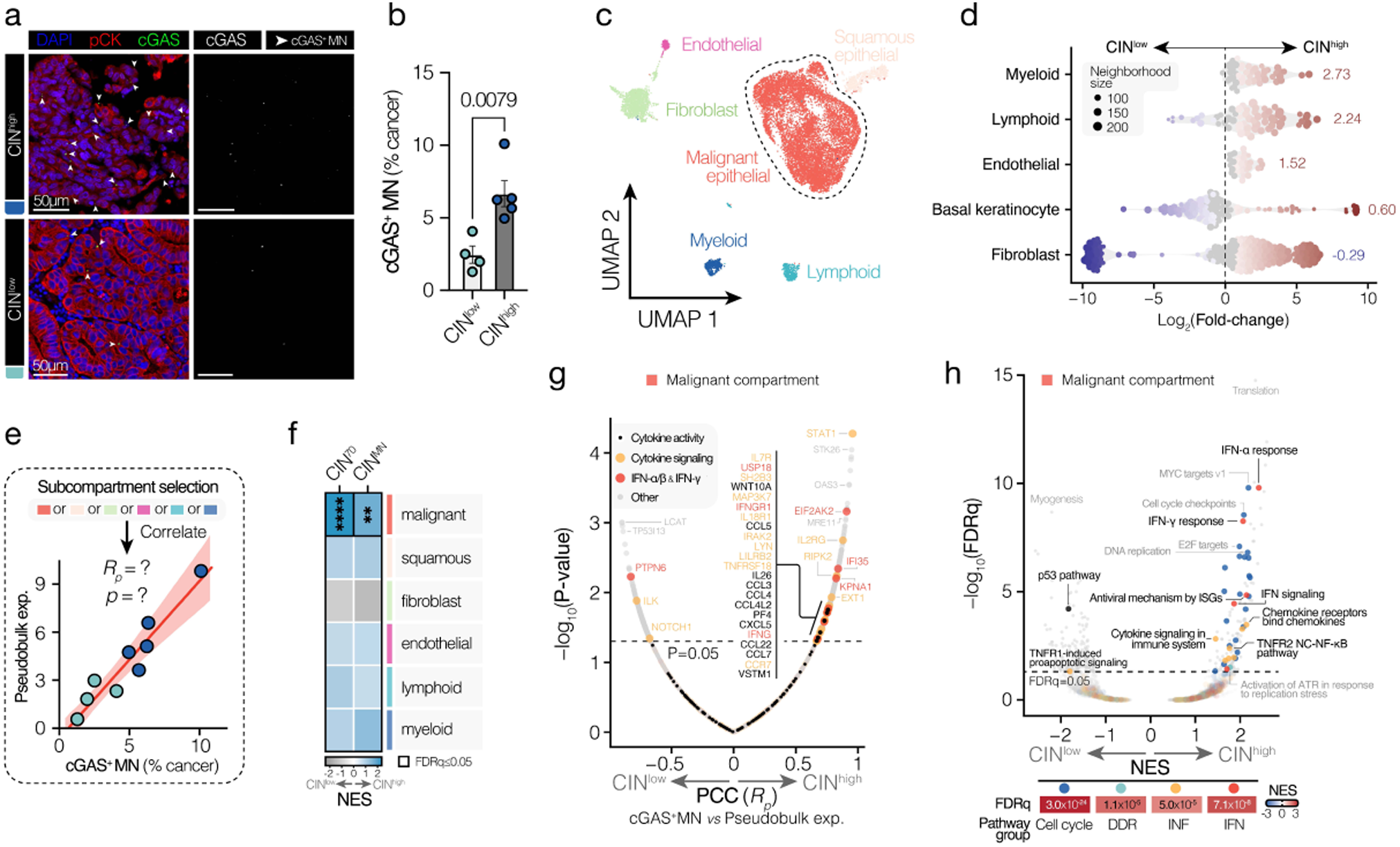
Micronucleus burden scales with innate immune activity in human EAC tumors. (**a**) Representative high-resolution images of human EAC tumor specimens stained with DAPI (DNA), anti-cGAS Ab and anti-pCK Ab showing a high frequency of cGAS^+^ MN in a snRNA-seq-shortlisted tumor from the upper tertile of the cGAS^+^ MN range of n=45 scored primary EAC tumors and a low frequency in a shortlisted tumor from the bottom tertile. Scale bar corresponds to 50 μM. (**b**) Tumoral cGAS^+^ MN (cancer cell compartment-specific) frequencies for n=9 tumors shortlisted for snRNA-seq, showing a clear separation between CIN^high^ tumors and CIN^low^ tumors. Bars represent the mean ± SEM. Data were analyzed by two-tailed unpaired t-test. (**c**) Uniform manifold approximation and projection (UMAP) dimensionality reduction (using Harmony embeddings to remove batch effects) of all cells remaining after quality control (QC) and filtering, colored by high-level manually curated cell identity. (**d**) Beeswarm plot of Milo analysis outputs for non-malignant cell clusters. Dot size corresponds to neighborhood size. Directionality and weighted mean indicate the CIN category (CIN^high^ or CIN^low^) in which there is a greater relative abundance of the indicated cell type. Weighted means for each cell cluster are shown. The x-axis represents log_2_(fold-change) in relative cell abundance. (**e**) Hit selection strategy for identifying genes most strongly associated with tumoral cGAS^+^ MN frequencies in each cell compartment. Single-cell expression matrices were pseudobulked for each cell type on a patient sample level, yielding a single gene expression value for each gene in every sample’s cell compartment. Resulting pseudobulked expression values for each gene were then iteratively correlated with cGAS^+^ MN frequencies across patient samples, yielding Pearson correlation coefficients (R_p_) and p-values for each gene in each cell compartment. (**f**) Heatmap of gene set enrichment analyses (GSEA) querying the degree of CIN signature (CIN^MN^ and CIN^70^ signatures) enrichment in ranked lists of cGAS^+^ MN-correlated genes in each cell compartment showing selective enrichment of CIN signatures among cGAS^+^ MN-associated genes in the malignant compartment. Color maps to normalized enrichment score (NES). Cell border indicates a significance of enrichment (FDRq) ≤ 0.05. (**g**) Volcano plot showing Pearson correlation coefficients (x-axis) and p-values (y-axis) of the association between a malignant compartment pseudobulked expression and tumoral cGAS^+^ MN frequency, showing a strong association between immune target expression in the malignant compartment and CIN. Genes annotated with the ‘Cytokine signaling’ gene ontology term (GO:0019221) are highlighted in orange. Genes with known IFN-α/β or IFN-γ (R-HSA-909733 or R-HSA-877300) pathway involvement are highlighted in red. Genes with reported cytokine activity (GO:0005125) are marked by a black central dot. (**h**) Volcano plot of gene set enrichment analysis (GSEA) outputs, showing the relationship between pathway enrichment and tumoral cGAS^+^ MN burden (CIN). Select pathways have been categorized based on their broad biological function into one of four major pathway groups: cell cycle-related pathways, DNA damage response (DDR)-related pathways, inflammatory response pathways (INF) or interferon response pathways (IFN). The p53 pathway (from the MSigDB Hallmarks collection) has been highlighted given its strong negative association with tumoral cGAS^+^ MN burden. The heatmap at the bottom shows the degree of enrichment of broad pathway groups with CIN within the overall distribution of all pathways. **** p ≤0.0001; ** p ≤0.01.

In total, we obtained the transcriptomes of 16,127 cells. Graph-based clustering of transcriptomes, followed by manual curation of sub-clusters, generated 6 major cellular compartments, comprising epithelial cells (malignant EAC cells and squamous epithelial keratinocytes), immune cells (lymphoid and myeloid cells), fibroblasts and endothelial cells, which exhibited gene markers and pathway enrichment in line with assigned cluster identities (**Fig. 4c, Extended Data Fig. 9c–f**). Malignant EAC cells (12,201 cells) comprised the majority of detected cells. Both lymphoid and myeloid cells were enriched in tumors with higher preponderances of cGAS^+^ MN, suggesting that CIN^high^ tumors are more heavily immune infiltrated (**Fig. 4d, Extended Data Fig. 9g**).

Whilst abundant lymphocytic infiltrate is typically indicative of effective anti-tumor immune responses, our observation that CIN^high^ tumors are also strongly infiltrated with myeloid cells and tumor-associated macrophages (TAMs) suggests an elevated degree of myeloid-mediated immunosuppression.

To identify tumor cell-intrinsic expression patterns associated with tumoral CIN levels, we performed linear correlation analysis between microscopy-assessed cGAS^+^ MN frequencies and compartment-specific pseudobulked gene expression, focusing on the malignant cell compartment (**Fig. 4e**). We found that CIN^70^ and CIN^MN^ signatures were significantly and selectively enriched across CIN-correlated genes in the malignant compartment (**Fig. 4f**). This expected specificity to the malignant compartment supported that this approach could identify cell type-specific CIN-associated expression patterns. Multiple cytokine and IFN-signaling related genes were strongly positively correlated with tumoral cGAS^+^ MN frequencies (**Fig. 4g**). These included *STAT1* as well as *IFNG* and ISGs including *OAS3* and *EIF2AK2 (*encoding the double-stranded RNA sensor PKR). Immune targets that scaled with cGAS^+^ MN included *CCL3*, *CCL4*, *PF4*, the CXCR2 ligand *CXCL5* and CCR2 ligand *CCL7*.

Upregulation of immune targets was reflected at a pathway level, with significant enrichment of multiple immune pathways with increasing CIN, including interferon-α and interferon-γ response pathways as well as cytokine and non-canonical NF-κB signaling (**Fig. 4h**).

Whilst a significant linear association between *CXCL8* and cGAS^+^ MN burden was not observed, the overall abundance of *CXCL8* mRNA and other *in vitro* cytokine hits (*CXCL1, CXCL2, CXCL3, CSF3*) was broadly increased in the malignant compartments of CIN^high^ tumors (**Extended Data Fig. 9h**). Enhanced immune target expression in CIN^high^ tumors was further accompanied by the expression of immune checkpoints in the malignant compartment including *CD274* (PD-L1), *IDO1, CD47* and the cGAS–STING checkpoint *ENPP1* (**Extended Data Fig. 9h**). Together with increased expression of T cell exhaustion markers in the lymphoid compartment, (**Extended Data Fig. 9i**), this is potentially indicative of ongoing evasion of both myeloid-mediated (e.g. CD47-SIRP1α phagocytosis checkpoint) and T cell-mediated tumor cell killing in CIN^high^ EAC.

### Mapping the tumor microenvironment of CIN^high^ tumors

As our data supported a strong link between CIN-driven innate immune signaling with a distinct chemokine signaling pattern and increased myeloid infiltrate, we sought to further characterize the tumor microenvironmental landscape of EAC tumors with ongoing CIN (**Fig. 5a**). Two tissue microarrays (TMAs), comprising 50 primary EAC tumor cores (24 treatment-naïve biopsy samples, 26 resections, containing 21 matched pre-post neoadjuvant chemotherapy samples) were profiled.

**Figure 5.**
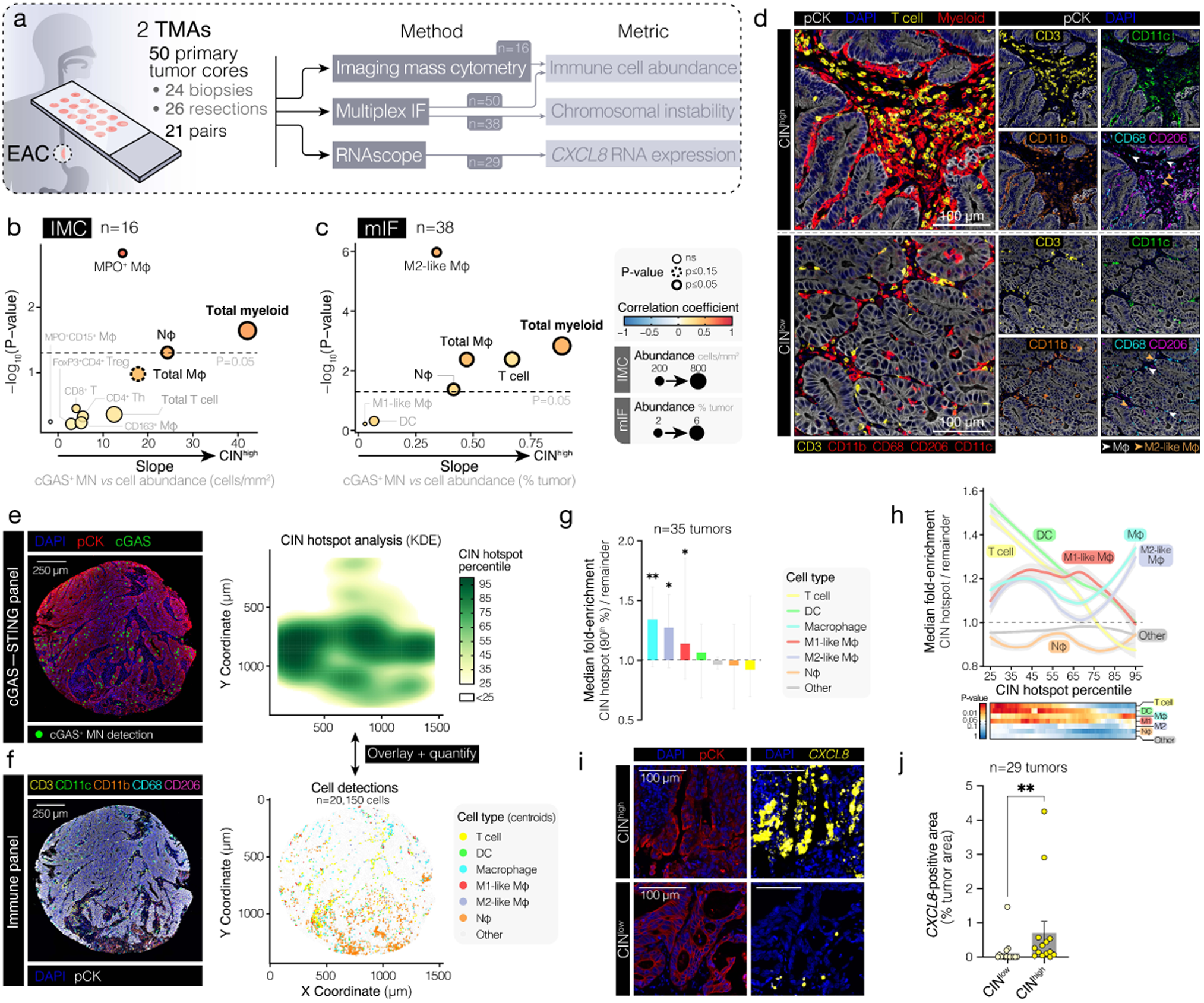
CIN is associated with intratumor myeloid cell inflammation in human EAC tumors. (**a**) EAC tumor microenvironment profiling workflow. Intratumoral immune composition and abundance were queried through multiplex immunofluorescence (mIF) and imaging mass cytometry (IMC). The extent of CIN was evaluated through mIF-based cGAS^+^ MN detection. *In situ* tumor cell-intrinsic *CXCL8* mRNA expression was measured using RNAscope. (**b, c**) Volcano plot showing immune cell subtypes whose intratumoral density is associated with the extent of tumoral CIN (cGAS^+^ MN frequency) (**b**) across n=16 human EAC biopsies queried by multiplexed imaging mass cytometry (IMC) or (**c**) n=38 pre- and pos-treatment EAC tumors stained via multiplex immunofluorescence (mIF). Immune cell subtypes whose density correlates significantly (Pearson correlation) with cGAS^+^ MN frequency are highlighted with a thick border, as indicated. The x-axis represents the –log_10_-transformed Pearson correlation p-value. The y-axis maps to the slope of the association between CIN and intratumoral cell abundance. Dot size maps to the mean cell density or abundance of a given immune cell type across all samples. Dot color maps to Pearson correlation coefficient for (**b**) and Spearman correlation coefficient for (**c**). (**d**) Representative high-resolution images of CIN^high^ and CIN^low^ (based on observed cGAS^+^ MN frequencies) human EAC tumor biopsy specimens stained with DAPI (DNA), as well as anti-pan-cytokeratin (pCK), anti-CD11b, anti-CD68, anti-CD206, anti-CD3 and anti-CD11c antibodies, showing broadly increased myeloid cell marker (CD68, CD206, CD11c, CD11b) predominance in the CIN^high^ tumor. Scale bars correspond to 100 μm. (**e**) *Left panel:* Representative high-resolution image of a human EAC tumor core stained with DAPI (DNA), as well as anti-pan-cytokeratin (pCK) and anti-cGAS antibodies. The spatial positions of cGAS^+^ MN detections are highlighted. Scale bar corresponds to 250 μm. *Right panel:* Heatmap of quantilized cGAS^+^ MN Kernel densities across the tumor area. (**f**) *Left panel:* Representative high-resolution image of a human EAC tumor core sample, directly adjacent (4 μm sections) to section in (**d**), stained with a multiplex immune antibody panel. Scale bar corresponds to 250 μm. *Right panel:* Spatial positions of classified cell centroids. (**g**) Fold-enrichment of the relative abundance of immune cell types in cGAS^+^ MN-dense areas (≥90^th^ percentile: ‘CIN hotspot’) of the tumor versus the remainder of the tumor (<90^th^ percentile) across n=35 EAC tumors. Bars are shown as the median with 95% confidence interval. A one-sample t-test was used to compare immune cell fold-enrichment in CIN hotspots across tumors to 1. (**h**) Line plot of median fold-enrichment of immune cell types in CIN hotspots across increasing hotspot percentiles, showing a progressively higher relative abundance of Mφs and M2-like Mφs and lower T cell and DC abundance in increasingly cGAS^+^ MN dense regions. Heatmap indicates significance of enrichment by one-sample t-test at each CIN density threshold. (**i**) Representative high-resolution immunofluorescence images of *CXCL8* RNAscope in CIN^high^ and CIN^low^ EAC tumors. (**j**) Tumor-cell intrinsic *CXCL8* mRNA expression levels in CIN^high^ versus CIN^low^ EAC tumors. Bars represent the mean ± SEM from n=29 EAC tumors. Significance was determined by Mann-Whitney U test. ** p ≤0.01, * p ≤0.05.

Treatment-naïve samples were first profiled using imaging mass cytometry (IMC), employing a 32-color hyperplexed panel, with 16 samples suitable for detailed immunophenotyping (**Extended Data Fig. 10a–d, Supplementary Table 2**). Correlating mIF-derived cGAS^+^ MN scores with immune cell abundances across tumors revealed an overall enrichment of intratumoral myeloid cells with increasing tumor CIN. Specific myeloid subsets exhibiting particularly strong associations with CIN included CD68^+^CD163^+^ myeloperoxidase (MPO)^+^ macrophages (MPO^+^Mac), a pro-tumorigenic tumor-associated macrophage (TAM) population)^31^, as well as tumor-associated neutrophils (CD15^+^HIF-1α^+^; **Fig. 5b**).

To further characterize CIN-associated immune infiltrates in a broader cohort (comprising 38 tumors with assessable CIN scores), we stained our TMAs using a mIF panel that included antibodies against myeloid cell markers (CD11c, CD11b, CD68, CD206), as well as CD3, pan-cytokeratin (pCK), and DAPI (**Extended Data Fig. 11a–c**). Observed patterns of immune marker positivity overlap, reflecting the phenotypically plastic and heterogeneous myeloid compartment, were used to classify stromal cells as macrophages (CD68^+^; Mφ), M1-like (CD68^+^CD11c^+^CD206^-^) macrophages, M2-like (CD68^+^CD206^+^) macrophages, dendritic cells (DCs; CD11c^+^CD68^-^) and tumor-associated neutrophils (Nφ; CD11b^+^CD68^-^; **Extended Data Fig. 11d–f**). CD3^+^ cells were classified as T cells. Interestingly, peripheral blood neutrophil counts correlated with intratumoral neutrophil infiltrate, but not other immune cell infiltrate (**Extended Data Fig. 11g**), suggesting that intratumoral neutrophil inflammation tracks peripheral neutrophil inflammation. Multiplex IF-derived immune cell abundances correlated closely with IMC-derived measurements (**Extended Data Fig. 11h**). Consistent with IMC results, in the broader mIF-stained TMA cohort, overall myeloid infiltration, as well as M2-like macrophage and tumor-associated neutrophil abundance, again showed significant associations with cGAS^+^ MN frequencies **(Fig. 5c,d**).

To map immune cell infiltrates within CIN^high^ niches (‘CIN hotspots’) for each tumor, we spatially aligned cGAS^+^ MN detections onto adjacent mIF immune panel-stained images and used a probability density function estimation approach to define CIN hotspots (**Fig. 5e**). We then quantified the relative proportion of infiltrated immune cells within CIN hotspots versus the remainder of the tumor (**Fig. 5f**). Tumor regions with high cGAS^+^ MN densities were proportionally enriched for macrophages and M2-like macrophages. In contrast, T cells and DCs appeared to be largely restricted from CIN^high^ regions (**Fig. 5g,h**).

As these data suggested that cGAS^+^ MN-rich tumor regions were preferentially macrophage- and myeloid cell-infiltrated, we sought to confirm whether elevated cGAS^+^ MN frequencies coincided with enhanced CIN-associated myeloid-attracting chemokine expression *in situ*. To do so, we performed *CXCL8* RNA *in situ* hybridization^32^, measuring the extent of *CXCL8* expression in malignant and stromal regions of tumor cores (**Extended Data Fig. 12a,b**). Across tumor cores, malignant cell expression of *CXCL8* was associated predominantly with cGAS^+^ MN frequencies (**Fig. 5h,i**), as well as with intratumoral myeloid cell population (macrophages, neutrophils) infiltration (**Extended Data Fig. 12c**). In contrast, *CXCL8* expression in the stromal compartment was most significantly correlated with infiltrated neutrophil abundance, in keeping with its high reported expression in this cell population ^33,34^ (**Extended Data Fig. 12d,e**). Together, these data indicate that IL-8 derived from CIN^high^ tumor cells results in an inflamed, myeloid-skewed TME typically associated with pro-tumorigenic activity.

To evaluate the impact of IL-8 on myeloid cell recruitment in CIN settings, we leveraged our *in vitro* engineered chronic CIN (isogenic CP-A cells) and acute CIN (MPS1i)-driven cell line models using transwell migration assays to measure the migration of peripheral blood-derived human CD14^+^ monocytes – important precursors of tumor-infiltrating myeloid cells^35^ – towards cell line-conditioned media (CM; **Extended Data Fig. 13a**). All three independent CIN^high^ CP-A p53^KO^dnMCAK clones exerted a significantly increased monocyte recruitment capacity over the matched genomically stable CP-A Cas9 clone (**Extended Data Fig. 13b**). Pre-incubation of monocytes with the CXCR1/2 inhibitor reparixin almost completely abrogated monocyte recruitment, demonstrating that CIN-dependent recruitment capacity was largely dependent on the production of CXCR1/2 ligands, many of which we identified as major CIN-dependent chemokines. Similar results were observed using MPS1i-driven induction of CIN in OE33 and SK-GT-4 cells, where MPS1i resulted in monocyte recruitment in CM from cGAS-proficient, but not in cGAS^KO^, cells (**Extended Data Fig. 13c,d**). The effect of cGAS^KO^ was partially recapitulated by CXCR1/2 inhibition, which did not abolish recruitment altogether, suggesting that additional cGAS–STING dependent chemotactic factors beyond CXCR1/2 ligands are secreted by CIN^high^ cells.

### CIN reshapes the tumor microenvironment and drives therapeutic resistance in EAC

As our results indicated that elevations in CIN were associated with myeloid-enriched, immunosuppressive and potentially therapy-resistant microenvironments, we investigated the relationship between response to neoadjuvant chemotherapy (NA Tx) and immune features, including intratumoral immune infiltration, the extent of CIN and cGAS–STING protein expression across pre-treatment tumors (**Extended Data Fig. 14a,b**).

Expectedly, non-response to neoadjuvant chemotherapy (defined as Tumor Regression Grades 4–5; TRG, also known as Mandard score) was associated with poor overall and recurrence-free survival (**Extended Data Fig. 14c**). In keeping with previous reports^36^, increased T cell and DC infiltrate was associated with improved response to neoadjuvant chemotherapy (defined as TRG 1–2; **Extended Data Fig. 14d**), whereas an increased intratumoral macrophage to T cell skew was associated with resistance (**Extended Data Fig. 14e**). However, neither cGAS^+^ MN burden, nor cGAS or STING protein expression alone were significantly associated with response (**Extended Data Fig. 14f,g**).

We employed a linear modelling approach to assess whether the extent of CIN exhibited response-specific associations with tumor immune features (**Extended Data Fig. 14h**). In contrast to NA Tx responders, CIN^high^ non-responder tumors were marked by a decrease in STING protein levels in malignant cells, consistent with previously reported STING tachyphylaxis in CIN^high^ settings associated with a rewired cGAS–STING pathway driving a tumor-supportive myeloid infiltrate^17^ (**Extended Data Fig. 14h,i**). Consistent with this, increased CIN was associated with an increased macrophage:T cell skew, specifically in non-responder versus responder tumors (**Extended Data Fig. 14h,j**). Pre-treatment tumors from non-responder patients were predominantly composed of CIN^high^ myeloid-dominated, and CIN^high^STING^low^ tumors, features that were largely absent among responder patients (**Extended Data Fig. 14k,l**).

To identify the role of CIN in clinical outcomes in EAC, we calculated CIN^MN^ scores derived from bulk RNA-seq data to stratify EAC tumors across 3 independent datasets (TCGA [n=87]^37,38^, DOCTOR^39^ [n=79] and SCOPE1^40^ [n=40] trials). Based on our results supporting myeloid:T cell enrichment as a significantly associated parameter with poor response to neoadjuvant chemotherapy, we derived a myeloid enrichment score measuring myeloid:lymphoid skew using gene signatures derived from our snRNA-seq data (**Supplementary Table 1**). Univariate Cox analyses across datasets supported the association of CIN^MN^ scores with worse pathological outcomes (**Fig. 6a**). Consistent with our findings, an increased myeloid:lymphoid skew and cGAS–STING mRNA abundance were also associated with clinical outcomes.

**Figure 6.**
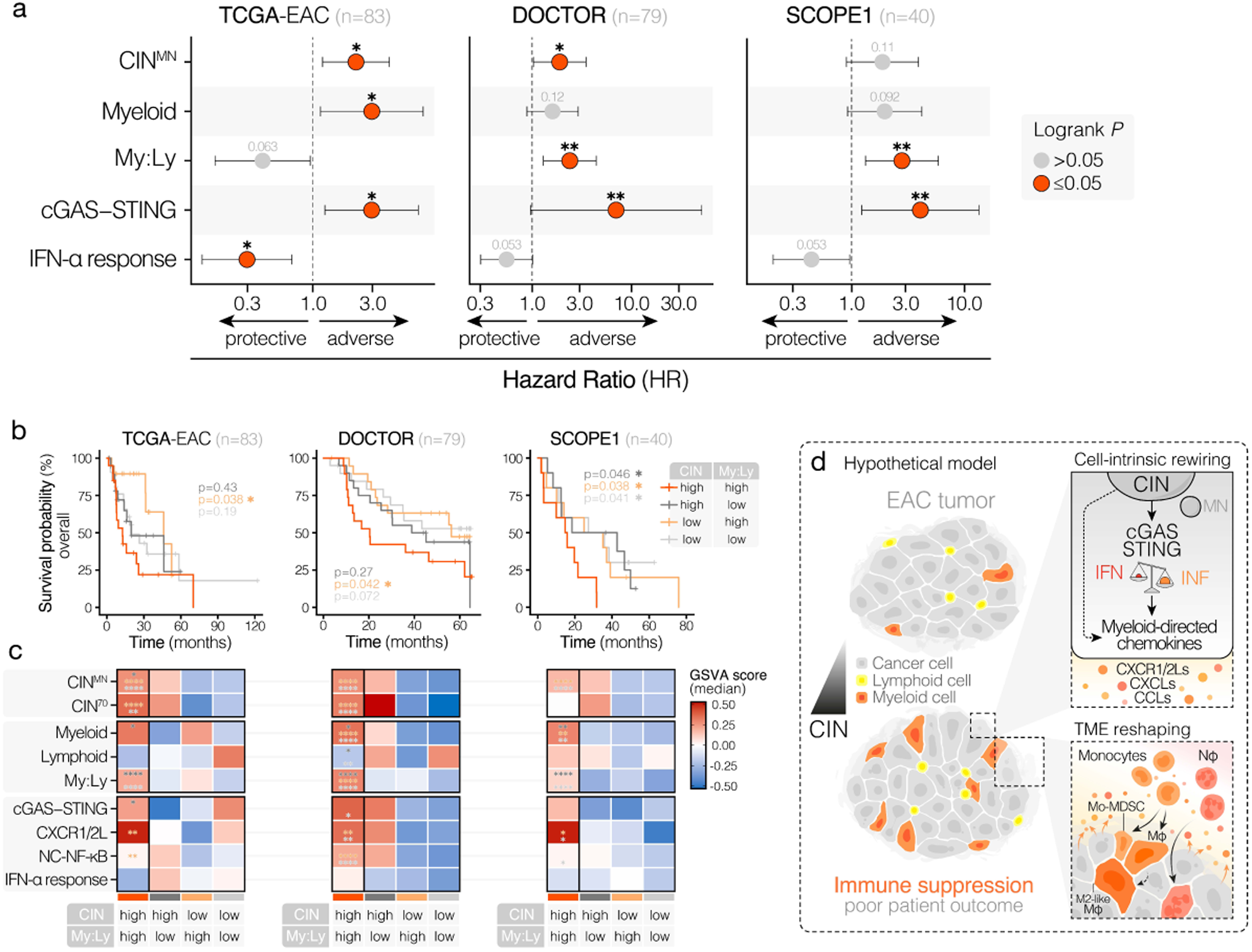
Chromosomally unstable, myeloid-dominated EAC tumors are associated with poor patient prognosis. (**a**) Forest plots of Cox proportional hazards models, showing the hazard ratio of high expression (determined using the maximal statistic approach) for indicated signatures in three independent EAC RNA-seq cohorts. Significance was determined using the log-rank test. (**b**) Kaplan–Meier curves showing the overall survival probability of human EAC patients partitioned according to CIN^MN^ score (median stratification) and subsequent partitioning (median) of CIN^high^ and CIN^low^ groups based on the degree of myeloid cell enrichment. Myeloid cell enrichment was determined by computing the extent of myeloid to lymphoid cell skew (My:Ly) for each patient using snRNA-seq-derived immune cell type signatures. Significance was determined through pairwise comparisons between the CIN^high^, myeloid-dominated group and other groups using the log-rank test. (**c**) Heatmaps showing the median expression for indicated signatures among patient clusters of interest. Asterisks denote the significance, determined through ANOVA with Tukey’s HSD, of the difference between the CIN^high^ myeloid-dominated cluster and other clusters (denoted by asterisk color). **** p ≤0.0001, *** p ≤0.001, ** p ≤0.01, * p ≤0.05. (**d**) Hypothetical model through which CIN drives cGAS–STING-dependent and -independent intratumor myeloid cell inflammation.

CIN^MN^ high, myeloid-dominated tumors were consistently characterized by elevated CXCR1/2 ligands, non-canonical NF-κB signaling and cGAS–STING expression, and exhibited significantly worse prognoses across the TCGA cohort and trial cohorts of EAC patients treated with chemotherapy and chemoradiotherapy^39,40^ (**Fig. 6b,c**). Collectively, our results indicate that cGAS-dependent chemokine signaling in CIN^high^ EAC contributes to a profoundly immunosuppressive myeloid-rich TME, resulting in non-response to neoadjuvant therapy and poor clinical outcomes for patients with this aggressive cancer subtype (**Fig. 6d**).

## Discussion

Chromosomal instability and chronic inflammation are prevalent features of EAC, emerging early in disease progression and promoting the aggressive characteristics of this tumor. Here, we demonstrate that EAC cells largely retain proficient cGAS–STING pathway machinery, whose activation by CIN-associated cytoplasmic dsDNA converges on a prominent engagement of inflammatory signaling and the production of cytokines and chemokines acting on myeloid leukocytes. By coupling the scoring of cGAS^+^ MN, which we validated as correlating to CIN by complementary WGS of cells and tumors, with snRNA-seq and spatial mapping of human EAC tumors, we demonstrate that CIN^high^ tumors are associated with heightened inflammatory activity and a myeloid-enriched, immunosuppressive microenvironment.

Our data revealed enrichment of negative regulators of type I IFN signaling, including upregulation of the ubiquitin-specific peptidase 18 (USP18), as well as interferon-induced protein 35 (IFI35), associated with CIN^high^ tumors. In contrast, we also observed that IFN-responsive genes, including STAT1, scaled with tumoral cGAS^+^ MN burden – in line with recent findings in genomically unstable breast cancer^41^ – suggesting that enhanced STAT1 activation may impose IFN I-centered selection pressures on CIN^high^ EAC. Of note, our *ex vivo* findings of STAT1, IFN-α and IFN-γ response pathway enrichment with increasing CIN differed from *in vitro* results which indicated downregulation of IFN response pathways with CIN. This can be partly accounted for by the complexity of the tumor microenvironment compared to a 2D cell line model, with amplification of CIN-induced immune signaling resulting in a rewired tumor microenvironment. Our data is in keeping with recent reports that highlight potential tumor promotion in chronically-IFN active TMEs, whereby targeting of IFN or JAK/STAT pathways can sensitize tumors to immune checkpoint blockade^42–45^. Whether this is of similar therapeutic potential in patients with CIN^high^ EAC remains to be elucidated.

Our findings contrast with reports of aneuploid tumors as “immune cold”^46,47^. This has been attributed in part to loss of chromosome 9p and the associated IFN gene cluster, and thus loss of interferon and inflammatory signaling, as well as loss of antigen presentation machinery^48–50^. However, we find sustained and enhanced inflammatory signaling with increased CIN in EAC, associated with an inflamed tumor microenvironment. Whilst this could be a site-specific phenomenon, it may reflect that CIN^high^ and aneuploid tumors are distinct entities, with associated patterns of immune infiltrate and inflammatory signaling. Making this differentiation is crucial, as aneuploid tumors and tumors with ongoing CIN may not share overlapping therapeutic vulnerabilities. A challenge in measuring CIN has been the dynamic nature of this state, as a genomic snapshot of a CIN^high^ tumor resembles that of an aneuploid tumor. Therefore, measurement of cGAS^+^ MN alongside myeloid infiltrates may enable distinction of actively CIN tumors compared to those in a steady state of aneuploidy, with inflammatory or immune restricted tumor microenvironments, respectively.

Here, cGAS^+^ micronuclei were scored as a proxy for the extent of cGAS–STING-activating chromosome segregation errors. However, recent reports have suggested that micronuclei may not be the sole or potentially even the primary source of cGAS activation in response to CIN^51,52^. CIN^high^ tumor cells may contain diverse nucleic acid species in their cytosol beyond micronucleated genomic DNA, including dsRNA^53^ and RNA–DNA hybrids^54^, that may activate cGAS or engage other innate immune cytosolic sensors such as MDA5 or RIG-I. The exact biochemical nature of the CIN-associated cGAS agonist remains to be established, although it is most likely closely linked to micronuclei formation.

The chemokine IL-8 (*CXCL8*) is heavily implicated in promoting tumor progression, via direct impact on cancer cell proliferation as well as through the recruitment of myeloid-derived suppressor cells, tumor-associated neutrophils and macrophages^55^. Elevated circulating IL-8 in patients receiving immune checkpoint blockade has been reportedly associated with poorer outcomes^56,57^. However, the source of IL-8 has been unclear. Here, we identify CIN-driven cGAS activation as resulting in increased *CXCL8* expression in CIN^high^ cancers, thereby accounting for the intratumoral myeloid infiltrate observed. This is consistent with *in vivo* studies which have reported that the murine homolog *Cxcl1* is enriched in CIN^high^ tumor models^17,21^. Targeting the receptors for IL-8, particularly CXCR2, has been explored in early phase clinical trials in prostate cancer^58^ and hepatocellular cancer^59^. Initial responses are promising, although peripheral neutropenia and risk of bacterial sepsis suggests that tumor-specific targeting may be required. Importantly, mice do not express *CXCL8*, and although homologs *Cxcl1* and *Cxcl2* recapitulate some functions, these do not perfectly mirror human *CXCL8* with non-redundant roles reported^60^. Species-restricted biology of the IL-8–CXCR1/2 axis therefore requires near-human models to accurately understand its role in tumor growth and therapeutic response^61^.

Locally and systemically administered STING agonists are under clinical development in combination with immune checkpoint inhibitors^62^. Early results of intratumorally administered STING agonists have not demonstrated broad efficacy. A potential explanation for the discrepancies observed between murine models and patient responses could be that STING expressed in tumor cells is poised to induce a type of inflammation that is immunosuppressive and pro-tumor. CIN^high^ tumors with chronic cGAS–STING activation may, therefore, be less likely to benefit from STING agonist strategies.

The poor clinical outcomes experienced by patients with CIN^high^ tumors are unlikely to be rescued by the advent of immune checkpoint inhibitors in EAC^63,64^ as CIN is associated with resistance to immunotherapy^65^. Therefore, innovative therapeutic interventions are urgently required to transform care for patients with CIN^high^ cancers. Our findings explain the counterintuitive maintenance and heightened expression of cGAS and STING in these aggressive tumors, and suggest that therapeutic approaches that disrupt the cGAS-driven pro-tumor inflammatory axis in CIN^high^ tumors may be effective strategies.

## Methods

### Cell culture

ESO-26, ESO-51, OACM5.1C, OE19, OE33 and SK-GT-4 cell lines were cultured in RPMI-1640, FLO-1 EAC and HEK293T cells were cultured in DMEM, KYAE-1 cells were cultured in RPMI-1640:Hams F12 (1:1 ratio), and hTERT-immortalized RPE-1 cells were cultured in DMEM:Ham’s F-12. Media was supplemented with 10% FBS and 100 U/mL penicillin and streptomycin P/S. Barrett’s esophagus-derived hTERT-immortalized CP-A (ATCC, KR-42421) and CP-C (ATCC, CRL-4029) cells^27,66^ were cultured in MCDB-153 supplemented with 0.4 μg/mL hydrocortisone, 20 ng/mL recombinant human epidermal growth factor, 8.4 μg/mL cholera toxin, 20 mg/mL adenine, 140 μg/mL bovine pituitary extract, 1X insulin-transferrin-selenium, 4 mM L-glutamine, 200 U/mL P/S and 5% FBS. All cell lines were maintained in a 37°C humidified incubator at 5% CO_2_ and routinely tested negative for mycoplasma contamination.

### Vector engineering and stable cell line generation

To generate cell lines with *TP53, CDKN2A,* and *CGAS* KO, target sequences were cloned into pL-CRISPR.SFFV.eGFP (Addgene plasmid #57827), pL-CRISPR.SFFV.tRFP (Addgene plasmid #57826) or lentiCRISPR v2 (Addgene plasmid #52961) transfer vector backbones according to the lentiCRISPRv2-Puro target guide sequence cloning protocol^67^. Single-guide RNAs (sgRNAs) are summarized in **Supplementary Table 3**. Plasmids were transformed into One Shot Stbl3 Chemically Competent *E. coli* (Invitrogen) and successfully transformed clones were selected from LB-agar ampicillin (100 mg/ml) plates. To confirm successful sgRNA integration, plasmids were sequenced by Sanger sequencing (TubeSeq, Eurofins Scientific).

To obtain recombinant lentivirus for transduction, ∼3.0x10^6^ HEK293T cells were transfected with lipofectamine-2000 (Invitrogen, 11668027) and opti-MEM, supplemented with 5 μg of the appropriate transfer vector, 2.5 μg pCMV-VSV-G (Addgene, #8454) envelope vector, and 2.5 μg pCMV-delta R8.2 (Addgene, #12263) packaging vector. Successful transfection of HEK293T cells with CRISPR.SFFV.eGFP (Addgene plasmid #57827) and CRISPR.SFFV.tRFP (Addgene plasmid #57826) vectors was confirmed by epifluorescence microscopy. Lentivirus was used to transduce CP-A or EAC cells in the presence of 5 μg/mL polybrene (Sigma-Aldrich). Successfully transduced cells were selected via 1.5 μg /mL puromycin treatment (for lentiCRISPR v2 [Addgene plasmid #52961] and pLV-CMV-dnMCAK [Addgene plasmid #205993] transduction) or flow cytometric single-cell sorting of GFP^+^ or GFP^+^RFP^+^ cells using a BD FACSAria Fusion Flow Cytometer (BD Biosciences; for pL-CRISPR.SFFV.eGFP and pL-CRISPR.SFFV.tRFP transductions). Single-cell clones from puromycin-selected mixed populations were obtained through limiting dilution in 96-well plates.

CP-A-derived single-cell clones (Cas9 control, p53^KO^, p53p16^DKO^ and p53^KO^dnMCAK) were screened by amplicon sequencing to confirm successful disruption of sgRNA target sites. PCR primers are detailed in **Supplementary Table 4**.

### Flow cytometry

To estimate the degree of GFP positivity in pL.CRISPR.SFFV.eGFP-(Cas9 control) and pL.CRISPR.SFFV.eGFP.sgTP53 (p53^KO^)-transduced CP-A cells, cells were trypsinized, washed twice by centrifugation (5 min at 300 x g) and resuspended in PBS. Data acquisition was performed using an Attune NxT Flow Cytometer (Thermo Fisher Scientific). SSC-A/FSC-A and FSC-A/FSC-H gates were set to exclude cell debris and cell doublets, respectively. RL1 and BL1 gates were set to select eGFP-positive cells. Data analysis was performed using the FlowJo (v.10.4) software.

### Immunoblotting

#### Cell lysate preparation

Cells were lysed in RIPA lysis buffer, (Thermo Fisher Scientific) supplemented with protease inhibitor cocktail (Roche, CO-RO) and phosphatase inhibitor (Roche, PHOSS-RO). Cell lysates were sonicated for 20s at 10% amplitude, and resulting homogenates were cleared by centrifugation at 16,000 x g for 15 min at 4°C. Protein concentrations were derived using the BCA Protein Assay (Thermo Fisher Scientific).

#### Western blotting

Lysates were boiled for 5 min at 90°C in 4X Laemmli Sample Buffer (Bio-Rad) supplemented with 355 mM 2-mercaptoethanol. Equal amounts of protein were resolved by SDS-PAGE on 4-20% polyacrylamide gels and transferred onto 0.2 μm nitrocellulose membranes. Membranes were blocked in 3% Bovine Serum Albumin (BSA) in Tris-buffered saline-Tween (TBS-T; 50 mM Tris-HCl [pH 7.5], 50 mM NaCl, 0.1% Tween-20) for 1h at room temperature (RT) and probed by overnight incubation in primary antibody-blocking solution at 4°C. For bound primary antibody (Ab) detection, membranes were incubated for 1h at RT in blocking buffer solution supplemented with either secondary anti-mouse (Cell Signaling Technology, #7076) or anti-rabbit (Cell Signaling Technology, #7074) streptavidin-conjugated horseradish peroxidase (HRP) Ab (1:1000). Membranes were incubated in Immobilon Western HRP substrate (Sigma-Aldrich) for enhanced chemiluminescence (ECL)-based protein detection and imaged using the ChemiDoc MP (Bio-Rad) or iBright FL1500 (Thermo Fisher Scientific) systems. Resulting images were analyzed using the Image Lab Software 5.0 (Bio-Rad). Primary antibodies are detailed in **Supplementary Table 5**.

### Immunofluorescence staining

#### Sample seeding

Adherent cells were seeded on glass coverslips. Suspension cells underwent treatments in suspension and were seeded onto poly-L/D-lysine-coated glass coverslips 6h before treatment endpoints. All cells were allowed to reach 70-90% confluence prior to sample collection.

#### Immunostaining

Cells were fixed with paraformaldehyde (PFA) for 12 min at RT and permeabilized for 5 min in 0.5% Triton X-100, followed by blocking in 2.5% BSA in 0.05% PBS-Tween for 1h at RT. For binding of primary Ab, coverslips were inverted onto 75 μL drops of primary anti-cGAS Ab solution (1:200; Cell Signaling Technology, #15102) in 2.5% BSA in PBS-Tween and incubated overnight at 4°C. Primary Ab were labelled by incubation in Alexa Fluor 488-conjugated goat anti-rabbit secondary antibody (Thermo Fisher Scientific, #R37116) in blocking solution for 1h at RT. Cells were counterstained for DNA by incubation in 5 μg/mL Hoechst for 15 min and mounted with Prolong Gold Antifade Mountant (Thermo Fisher Scientific).

#### Image acquisition, processing and analysis

Images were acquired with a 63x oil immersion objective on an Andor Dragonfly spinning disc confocal microscope (Oxford Instruments) or with a 40x objective on an Olympus SpinSR SoRa spinning disc confocal microscope (Olympus Life Science). Images were acquired as Z-stacks (3-11 slices) taken at 1 μm intervals. Maximum intensity-projected images were processed and analyzed using the Fiji image processing package of the ImageJ software^68^.Custom Fiji macros were used to count nuclei and guide manual counts for micronuclei and cGAS^+^ micronuclei (cGAS^+^ MN). CGAS^+^ chromatin bridges were scored manually.

### RT-qPCR

RNA isolation was performed using the RNeasy Plus Mini Kit (Qiagen) and complementary DNA (cDNA) was derived using the SuperScript III First-Strand Synthesis SuperMix (Thermo Fisher Scientific, 11752050) or HiScript IV SuperMix (Vazyme, R423-01). Quantitative RT-PCR was performed using Fast SYBR Green Mastermix (Applied Biosystems, 4385618) or Taq Pro Universal SYBR qPCR Master Mix (Vazyme, Q712-03) on a StepOnePlus Real-Time PCR System (Applied Biosystems). Relative abundance of mRNA was determined through the 2^-ΔΔCT^ method^69^ using *18S* as the reference gene. Primer pair sequences are listed in **Supplementary Table 6**.

### 2’3’-cGAMP stimulation

EAC cells were seeded at a density of 5.0x10⁵ cells per well in 6-well plates and allowed to adhere for 24h. Cells were then transfected with 2 μg/mL 2’3’-cGAMP (InvivoGen) or cGAMP diluent (sterile DEPC-treated water) with 2 μL/mL lipofectamine-2000 (Invitrogen) in opti-MEM media for 6h. Prior to transfection, 2’3’-cGAMP/diluent:lipofectamine transfection mixtures were incubated at RT for 20 min to facilitate transfection complex formation.

### 2’3’-cGAMP ELISA

Intracellular and extracellular 2’3’-cGAMP quantification was largely carried out as previously described^16^. Cells were seeded in 100 mm culture dishes and allowed to grow to a confluency of 80-90% before changing the media to reduced serum and phenol red-free opti-MEM media (Thermo Fisher Scientific). Cells were treated with 1 μM MPS1 inhibitor reversine or DMSO 48h prior to media exchange. For quantification of cGAMP levels following dsDNA-driven cGAS stimulation, cells were transfected with 2.5 μg/mL G_3_-YSD Y-form DNA (InvivoGen, tlrl-ydna) using 2 μL/mL lipofectamine-2000 (Invitrogen) in opti-MEM media for 24h.

For extracellular cGAMP, conditioned media was collected 24h following media change and centrifuged at 600 x g at 4°C for 15 mins. For intracellular cGAMP, adherent cells were collected and counted. Cell suspensions were pelleted for 15 min at 600 x g at 4°C, lysed in RIPA lysis buffer, and sonicated for 10s at 10% amplitude. Resulting homogenates were cleared by centrifugation and protein concentrations were derived by BCA assay. For intracellular cGAMP quantification, protein concentrations were adjusted to 0.5-1 mg/mL (kept consistent within experiments). Conditioned media for extracellular cGAMP quantification was assayed directly. cGAMP ELISA was performed using the 2’3’-Cyclic GAMP Competitive ELISA Kit (Invitrogen). 2’3’-cGAMP concentrations of cell supernatants were normalized to absolute cell counts for each sample.

### IL-8 ELISA

For quantification of IL-8 concentrations of cell culture supernatants, OE33, SK-GT-4 and CP-A cells were seeded at a density of 3.0x10^5^ or 5.0x10^5^ cells, respectively, in 100 mm cell culture dishes. After 24h, cell media was either replaced with fresh media or, where indicated, replaced with 1 μM reversine- or DMSO-treated media for 24h, followed by drug wash-out and replacement with fresh media. Cell culture media was conditioned for 48h, collected, and cleared by centrifugation at 1000 x g for 10 minutes at 4°C. IL-8 ELISA was performed using the human IL-8 ELISA kit (Thermo Fisher Scientific).

### Viability assay

Cells were plated in 96-well plates at a density of 1,500 or 2,000 cells per well (EAC cells and CP-A lines, respectively) to ensure mock-treated cells reached ∼90% confluency at the assay endpoint. After 24h, cells were treated with puromycin, Nutlin-3a (Selleck Chemicals) or reversine. Cell viability was assessed using the Cell Counting Kit 8 (CCK8; Abcam). WST-8 solution was added to wells 4h before treatment endpoint. Cell viability was determined by measuring absorbance at 460 nm and normalizing to the average absorbance of mock-treated wells.

### RNA-sequencing

#### RNA-sequencing and data pre-processing

EAC and CP-A cells were seeded at a density of 5.0x10^5^ cells in 100 mm cell culture dishes. After 24h, cell culture media was replaced with either fresh media, or 0.5 μM reversine- or DMSO-treated media, for CP-A and EAC cells, respectively, and incubated for an additional 48h. RNA was isolated using the RNeasy Plus Mini Kit (Qiagen) and sample quality was assessed using the 5400 Fragment Analyzer System (Agilent). Paired-end 150 base-pair (bp) sequencing of cDNA libraries was performed on the Illumina NovaSeq6000 platform. Reads were mapped to the human reference genome GRCh38 using HISAT2 (v.2.0.5)^70^. The resulting alignments were summarized and quantified at the gene level using featureCounts (v.1.5.0) to obtain read counts for downstream differential expression analysis^71^.

#### Identification of cGAS-dependent CIN-induced genes

To assess MPS1 inhibitor (MPS1i) treatment-induced gene expression changes in cGAS-proficient and cGAS^KO^ EAC cells, batch effects were corrected using the ‘ComBat’ function from the sva R package (v.3.36.0)^72^. Genes were filtered to retain those with a total read count of ≥ 5 counts across all samples and non-zero expression in ≥ 4 samples. Filtered datasets were analyzed for differential expression using DESeq2 (v.1.42.0)^73^ and significance was tested using the default Wald test, corrected for multiple testing via the Benjamini-Hochberg method, between n=3 biological replicates of MPS1i- and DMSO-treated samples. Significantly differentially expressed genes were identified using an adjusted p-value ≤0.01 cut-off.

Genes whose MPS1i-induced differential expression was cGAS-dependent, were identified by paired t-tests, comparing the log_2_(fold-change) of significantly differentially expressed genes between MPS1i and DMSO-treated samples in cGAS-proficient (Cas9) versus cGAS^KO^ cells. Genes that were significantly upregulated (adj. p ≤ 0.01, log_2_(FC) > 0) in MPS1i-treated cGAS-proficient cells with a significantly reduced MPS1i-induced upregulation (paired t-test, p ≤0.05; paired by batch number) in cGAS^KO^ cells were classified as cGAS-dependent MPS1i-induced genes. Conversely, genes that were significantly downregulated (adj. p ≤0.01, log_2_(FC) < 0) in MPS1i-treated cGAS-proficient cells, with a significantly less pronounced downregulation (p ≤0.05) in cGAS^KO^ cells, were considered cGAS-dependent MPS1i-suppressed genes.

#### Identification of cGAS^+^ MN-correlated genes

To identify genes whose expression correlated with the baseline cGAS^+^ MN burden of cell lines in our isogenic CP-A cell line model (Cas9 control, p53^KO^, p53p16^DKO^ and p53^KO^dnMCAK lines), gene-level counts were filtered to retain genes with a cumulative read count of ≥ 50 counts across all (n=30) samples and non-zero expression in ≥ 4 samples. Filtered counts were then normalized using the variance-stabilizing transformation (VST) from the DESeq2 package.

For each gene, a linear mixed-effects model was fitted using the ‘lm’ function of the nlme R package (v.3.1-165)^74^ to assess the relationship between gene expression and cGAS^+^ MN burden (the number of baseline cGAS^+^MN per 100 cells; i.e. cGAS^+^MN %). The model included the average cGAS^+^MN burden (n=3, as determined by IF) for each cell line as a fixed effect and incorporated random effects attributable to different clones (n=3 independent cell clones) and experimental batches (n=3 biological replicates), enabling identification of genes whose expression significantly scaled with cGAS^+^ MN burden while accounting for confounding effects stemming from clone and batch variability.

Specifically, the model was defined as follows:

> Expression_(gene X)_ ∼ average cGAS^+^MN + (1 | Clone / Batch)

For each gene, the p-value (ANOVA; indicating statistical significance of the association between expression and cGAS^+^ MN burden) and slope (magnitude change in gene expression per unit increase in cGAS^+^ MN burden) of the cGAS^+^ MN effect were extracted from the model output and used to identify hits whose expression was most strongly associated with micronucleus burden.

### Functional enrichment analyses

#### Gene set enrichment analysis

GSEA was performed using the fgsea (v.1.28.0) package^75^ in R, using the Hallmark and Canonical Pathways (CP; comprised of Reactome, Kyoto Encyclopedia of Genes and Genomes [KEGG], Pathway Interaction Database [PID], WikiPathways and BioCarta databases) collections from the Molecular Signatures Database (MSigDB)^76^, as well as Gene Ontology Biological Processes (GO:BP) database gene sets^77^. Gene set collections were retrieved using the msigdbr (v.7.5.1) package^78^.

To evaluate the broader distribution of enrichment for gene sets related to cell cycle, DNA damage repair (DDR), interferon (IFN) response, and inflammatory response (INF) pathways in GSEA outputs, gene sets were classified based on character strings in their assigned pathway names. Gene sets identified as belonging to Cell Cycle, DDR, IFN, or INF pathway groups were compiled into custom lists (**Supplementary Table 7**) and analyzed for enrichment through GSEA using rank metrics for gene sets computed from the original GSEA outputs.

#### Overrepresentation analysis

Overrepresentation analysis (ORA) was performed using the ‘gost’ function of the gprofiler2 R package (v.0.2.3)^79^, focusing on GO:BP and Reactome database pathways. Enrichment probabilities were computed using the default one-sided Fisher’s exact test corrected for multiple testing via the Benjamini-Hochberg method. Database terms mapping to >2,000 genes were excluded from the final analysis outputs, as they were considered too broad to provide meaningful insights.

### Single-cell whole-genome sequencing

To evaluate the karyotypes of our isogenic CP-A cell line model, ∼1x10^6^ cells of each genotype and clone (Pa, Cas9 control, p53^KO^, p53p16^DKO^, p53^KO^dnMCAK; clones 1, 2 and 3) were profiled using shallow single-cell whole-genome sequencing (scWGS) (Research Sequencing and iPSC/CRISPR Facility, ERIBA, University Medical Centre Groningen, University of Groningen).

Isolation and staining of nuclei, as well as nuclei sorting and library preparation were performed as previously described^80^, with small variations. Briefly, 24 single nuclei per cell line were dry sorted into 96-well plates. Overnight re-hydration in TPLK was performed and proteolysis started. MNase Digestion, AMPure Bead Cleanup and End Repair and A-Tailing steps were replaced in favor of the Fragmentase enzyme (NEBNext Ultra II FS DNA Library Prep Kit, New England Biolabs). Subsequent library preparation steps were unchanged.

Libraries were sequenced on a NextSeq2000 (Illumina) P1-100c (1x10^6^ reads per single cell) sequencer, using a 77bp read-length and 11bp dual index read configuration. Resulting FASTQ files were mapped to the human reference genome GRCh38 using the Burrows–Wheeler aligner^81^. The aligned read data (BAM files) were analyzed using the AneuFinder algorithm^28^. Following GC correction and blacklisting of artifact-prone regions (i.e. extreme low or high coverage in control samples), libraries were analyzed using the Dnacopy and Edivisive copy number calling algorithms with variable width bins (average bin size = 1 Mb; step size = 500 kb). Breakpoints were refined using refine.breakpoints = TRUE. Analyses were carried out using a euploid reference as described previously^80^.

Samples corresponding to p53^KO^dnMCAK clone 1 and p53^KO^ clone 2 were re-analyzed with the developer version of AneuFinder (Version 1.7.4; available at: https://github.com/ataudt/aneufinder^28^) using a minimum ground ploidy of 3 or 3.5 (min.ground.ploidy=3.0 or 3.5) and a maximum ground ploidy of 4 or 4.5 (max.ground.ploidy=4.0 or 4.5), respectively.

As some of the libraries of these two samples gave differing results (copy numbers too low compared to other libraries; less optimal fits), results were curated by requiring a minimum concordance of 90% between the results of the two algorithms.

Libraries with on average < 10 reads per bin and per chromosome copy were discarded. The aneuploidy score of each bin was calculated as the absolute difference between the observed copy number and the expected copy number when euploid. The score for each library was calculated as the weighted average of all the bins (size of the bin as weight) and the sample scores were calculated as the average of the scores of all libraries. The heterogeneity score of each bin was calculated as the proportion of pairwise comparisons (cell 1 vs cell 2, cell 1 vs cell 3, etc.) that showed a difference in copy number (e.g. cell 1: 2-somy and cell 2: 3-somy). The heterogeneity score of each sample was calculated as the weighted average of all the bin scores, using size of the bin as weight.

### Whole-genome sequencing

DNA was extracted from patient-matched tumor and normal (buffy coat) samples using the DNeasy Blood and Tissue Kit (Qiagen) according to the manufacturer’s protocol. DNA samples were quantified using the Qubit 2.0 Fluorometer (Life Technologies) and DNA integrity was checked using the TapeStation 4200 (Agilent). The NEBNext Ultra II DNA Library Prep kit (New England Biolabs) was used for DNA library preparation following the manufacturer’s recommendations. Briefly, genomic DNA was fragmented by acoustic shearing using a Covaris instrument. Fragmented DNA was end-repaired and adenylated. Adapters were ligated after adenylation of the 3’ ends, followed by enrichment by limited cycle PCR. Resulting adapter-ligated DNA libraries were cleaned up and validated using the Agilent TapeStation and quantified using a Qubit 2.0 Fluorometer.

Short-read WGS of multiplexed libraries was performed on the Illumina NovaSeq platform with a paired-end 150bp read configuration at a target coverage of 60x and 30x for tumor and normal samples, respectively. Resulting raw sequencing data (.bcl) files were converted to FASTQ format and de-multiplexed using the bcl2fastq software (Illumina).

Reference-based mapping on raw sequencing data was conducted by aligning FASTQ files to the GRChg38 reference genome using BWA-mem (v.0.717)^81^ generating BAM files for downstream analysis. Resulting BAM files were subsequently sorted, indexed, and marked for duplicates using Picard (v.3.1.0)^82^ and Samtools (v.1.15)^83^.

Structural variants (SVs), tumor purity and ploidy were called using the LINX pipeline (HMF tools, v.3.9)^84,85^. SVs were categorized by PURPLE as deletions, duplications, inversions or break-ends and were filtered to remove failed calls.

Copy number analysis was performed using ASCAT (v.3.1.2), JaBbA (v.1.0), and PURPLE. JaBbA^86^ was used to visualize copy number alterations (CNAs) and SVs for each tumor. The percentage of the genome altered was calculated using major copy number per segment data from ASCAT by determining the number of megabases that were amplified, deleted, or unaffected (copy number = 2). Amplifications were defined by a copy number > 2, and deletions by a copy number < 2. The percentage of the genome amplified was calculated using PURPLE when the copy number exceeded 2.

### Single-nucleus RNA sequencing

#### Tissue processing

To enable tissue-sparing extraction of nuclei for snRNA-seq from clinical-grade frozen tissue specimens, we optimized a previously described adaptation of the salt-tris (ST)-based extraction method^87, 88^ for EAC tissue samples. Clinical samples were collected under the Oxford Radcliffe Biobank (ORB)-approved study 21/A093.

Frozen tissue specimens were embedded in optimal cutting temperature (OCT) compound on dry ice and mounted on the sample holder of a LeicaCM1950 cryostat (Leica Biosystems) set to -20°C. Between 40-60 20 μm tissue scrolls were cut per tissue and stored at -80°C until further sample processing.

All subsequent sample processing steps were performed on wet ice using ice-cold buffers and centrifuges set to 4°C unless stated otherwise. For extraction of nuclei, sample tubes were moved from dry to wet ice and allowed to equilibrate for 30s. Next, 5 mL of sterile PBS was added to tubes and samples were inverted 3 times to dissolve OCT, followed by centrifugation at 300 x g for 2 min. Resulting tissue pellets were resuspended in 2 mL ST buffer (146 mM NaCl, 10 mM Tris-HCl, pH 7.5, 1 mM CaCl_2_ and 21 mM MgCl_2_ in DEPC-treated water) supplemented with 0.03% Tween-20 and 0.1% BSA. Cell suspensions were pipetted vigorously to mechanically disrupt the tissue and incubated on ice for 7.5 min, repeating the pipetting step every 2.5 min. Following the incubation period, homogenates were quenched by adding 1 mL of ST buffer with 40 U/mL of RNaseOUT ribonuclease inhibitor (Thermo Fisher Scientific) and filtered twice through 70 μm filters. Remnants of dissociated tissue in sample tubes were rinsed with 3 mL ST buffer with RNAse inhibitor and filtered to enhance nucleus yields. Nucleus suspensions were then centrifuged at 500 x g for 5 min and resuspended in 1 mL ST buffer. To determine the concentration and degree of dissociation of nuclei in the samples, nuclei were stained with Hoechst and counted using an EVOS M5000 benchtop fluorescence microscope (Thermo Fisher Scientific).

#### snRNA library preparation

14,000 nuclei (with the exception of sample #5, for which ∼7,600 nuclei were loaded) were loaded in ST buffer on a Chromium X using the Chromium Next GEM Single Cell 5’ Kit v2 (10x Genomics, PN-1000263) following the Chromium Next GEM Single Cell 5’ v2 (Dual Index) user guide (10x Genomics, CG000331 Rev E). GEMs underwent reverse transcription, cleanup using Dynabeads MyOne Silane beads (Thermo Fisher Scientific) and amplification. To account for the lower abundance of RNA in nuclei versus whole cells, an additional PCR amplification cycle was added (14 total cycles). Construction of final gene expression (GEX) libraries was performed using the Library Construction Kit (10x Genomics, PN-1000190) and Dual Index Kit TT set A (10x Genomics, PN-1000215) according to the user guide. The fragment size distribution of cDNA and final sequencing-ready GEX libraries was assessed using a TapeStation 2200 system (Agilent) with TapeStation D5000 reagents. The concentrations of final GEX libraries were determined using a Qubit fluorometer (Life Technologies).

#### snRNA library sequencing

GEX libraries were sequenced on the Illumina NovaSeq X Plus platform with a paired-end 150bp read configuration, covering ≥ 20,000 read pairs per cell. The sequencing run was set up with a 26-10-10-90 cycle configuration, consisting of 26 cycles for read 1, 10 cycles for the i7 index, 10 cycles on the i5 index, and 90 cycles for read 2. A 2% PhiX spike-in was included to monitor sequencing quality and provide an internal control.

#### Single-cell gene expression matrices

Generation of single-cell expression matrices, background noise removal, quality control and filtering were performed according to a 10x Genomics Cell Ranger workflow-based analysis pipeline as described previously^88^. De-multiplexed FASTQ files from snRNA-sequencing reads were aligned to the human GRCh38 reference genome and gene counts were quantified using the Cell Ranger (v.6.0.0, 10x Genomics) ‘count’ function. As nuclear mRNA is highly intronic, introns were included in the analysis.

#### Background noise removal

Technical ambient RNA and empty droplets were removed from gene expression matrices, using the ‘remove-background’ function of CellBender (v.0.2.0)^89^ on Cell Ranger-generated ‘raw_feature_bc_matrix.h5’ files. The ‘Estimated Number of Cells’ parameter inferred by Cell Ranger was used to define the ‘expected-cells’ for extraction. The ‘total-droplets-included’ parameter was adjusted to the midpoint of plateaus in the barcode-rank plot, based on visual inspection.

#### Quality control and filtering

Expression matrices were processed individually in R using the Seurat package (v.4.2.74)^90,91^. Retained cells had detected genes ranging between 200-9,000, UMI counts between 800-40,000 and mitochondrial reads < 15%. Doublets were removed using Scrublet v.0.2.1^92^.

Filtered gene-barcode matrices were then normalized using the Seurat ‘NormalizeData’ function and ‘LogNormalize’ method. Variable genes were identified by applying the ‘vst’ method in the ‘FindVariableFeatures’ function, selecting the top 2,000 variable genes. Next, gene expression matrices were scaled and centered using the ‘ScaleData’ function. Dimensionality reduction was performed using principal component analysis (PCA) and uniform manifold approximation and projection (UMAP) using the top 30 principal components.

#### Single-cell cluster curation

Pre-processed data were normalized using ‘SCTransform’, followed by PCA dimensionality reduction. The resulting top 30 SCT principal components were used to integrate data across multiple samples using Harmony (v.1.2.0)^93^, setting sample labels as grouping variables to enable correction for inherent batch effects. Integrated Harmony embeddings were then used to recompute the nearest neighbors and construct a shared nearest neighbors (SNN) graph. Clustering of the SNN was performed using the ‘FindClusters’ Seurat function (with res=0.35) to identify cell clusters reflecting distinct cell types. Visualization of clusters was performed through UMAP, using the top 30 principal components from Harmony-integrated PCA dimensionality reduction. Cluster identities were assigned by identifying differentially expressed genes between clusters using a variation of the negative binomial exact test, based on the sSeq method^94^ implemented in the Cell Ranger pipeline. Resulting cluster markers were cross-referenced with published literature to determine cell type identities and clusters with similar functional marker genes were manually curated to assign high-level cluster identities. Differential gene expression (DGE) analysis of curated clusters yielded high-level cluster markers and was used to perform GSEA using – *log_10_(adj.p-value) x sign(log_2_[FC])* from DGE outputs as gene ranking metrics, enabling assessment of cluster-level pathway enrichment.

#### Identification of cell type-specific cGAS^+^ MN-correlated genes

To enable assessment of cell-type-specific gene expression with tumoral cGAS^+^ MN burdens across samples, a pseudobulking approach was used to infer the relative cell-type-specific expression level of genes across samples. Unnormalized RNA counts for each sample were aggregated on a per-cluster basis using the ‘AggregateExpression’ function of the Seurat package. Aggregated expression matrices for each cell type were then normalized across samples using DESeq2. Expression values were then iteratively correlated with mIF-derived tumoral cGAS^+^ MN scores, yielding a Pearson correlation coefficient (R_p_) and p-value. To infer the degree of pathway-level scaling with cGAS^+^ MN, GSEA was performed using the R_p_ for each gene as the rank metric.

#### Differential neighborhood abundance testing

To quantify differential cell state abundances between CIN^high^ and CIN^low^ tumors, we used the R package Milo^95^, which models local variations in cellular composition using graph-based neighborhoods, using default parameters. Each neighborhood comprised an index cell and its surrounding cells in a k-nearest neighbor graph, representing discrete cell states within the dataset. For each predicted neighborhood, hypothesis testing between the compared groups was performed to identify differentially abundant cell states while controlling for false discovery rate (FDR). The number of cells in each neighborhood were counted on a per-sample basis to perform differential abundance testing. To correct for multiple hypothesis testing, a weighted FDR was used, accounting for the spatial overlap between neighborhoods.

#### Myeloid and lymphoid cell signature development

To define EAC-associated intratumoral myeloid and lymphoid cell gene signatures, pairwise DGE analyses were performed between all curated subcompartment clusters using Cell Ranger’s implementation of the sSeq method. The top 50 (ranked by significance) genes exclusively significantly upregulated in either myeloid or lymphoid subcompartments were defined as myeloid or lymphoid gene signatures, respectively, and are detailed in **Supplementary Table 1**.

### Multiplex immunofluorescence

#### TMA construction

The tissue microarray (TMA) samples were accessed from the ORB (study number 20/A144), cut at the Oxford Centre for Histopathology Research (OCHRe), annotated and reviewed by a gastrointestinal histopathologist (AE), and assembled by the Translational Histopathology Laboratory (THL, Oxford).

#### Multiplex immunostaining

Multiplex immunofluorescence (mIF) of human esophageal adenocarcinoma biopsies was carried out on 4 μm formalin-fixed paraffin-embedded (FFPE) sections on a Leica Bond RXm Autostainer (Leica Biosystems) using the Opal multiplex immunohistochemistry system (Akoya Biosciences).

Slides were baked and dewaxed using BOND dewax solution (Leica Biosystems, AR9222). Epitope retrieval was performed at 100°C for 20 min using the BOND Epitope Retrieval ER2 Solution (Leica Biosystems, AR9640), followed by endogenous peroxidase blocking for 5 min in 3-4% hydrogen peroxide. Multiplex immunostaining was performed in up to six successive cycles using the primary Ab-Opal fluorophore pairings, Ab dilutions and incubation periods are outlined in **Supplementary Table 8**.

Sections were counterstained with spectral DAPI (Akoya Biosciences, FP1490) and mounted with VECTASHIELD Vibrance Antifade Mounting Medium (Vector Laboratories, H-1700-10). Whole slide scans and multispectral images (MSIs) were acquired at 40x magnification on a Vectra Polaris Automated Quantitative Pathology Imaging System (Akoya Biosciences). Batch analysis of MSIs was performed with the inForm Tissue Analysis software (v.2.4.8). Batch-analyzed multispectral images were stitched on the HALO image analysis platform (Indica Labs) to generate spectrally unmixed reconstructed images.

#### Analysis of mIF images

Reconstructed mIF images were analyzed using QuPath (v.0.5.0)^96^. Cell segmentation was achieved using the built-in QuPath cell detection tool or StarDist (using the pretrained DSB 2018 model)^97^ through the DAPI counterstain. Detected cells were classified as epithelial (i.e. malignant) or stromal, or identified as artefactual (e.g. necrotic tissue), using supervised machine learning classifiers bespoke to each tumor specimen.

Random Trees object classifiers were trained on ≥ 10 manual annotations per compartment per tumor, guided by the DAPI and pan-cytokeratin (pCK) channels, with training overseen by a consultant gastrointestinal histopathologist (A.E.). Stromal cells were further classified by immune marker positivity and normalized to the total number of cell detections in tissue sections.

To enable between-tumor comparisons of marker protein levels (e.g. pCK, cGAS and STING), average marker intensity measurements for each cell were normalized to (divided by) its respective mean inferred autofluorescence (AF) value. Cancer and stromal compartment protein intensities were computed as the average of all the mean AF-normalized cellular intensities belonging to said compartment class.

#### cGAS^+^ micronucleus detection

To identify cGAS^+^ micronuclei (identified as small intense intracellular cGAS foci), tumor and stromal cell detections were merged into large stromal or tumoral cell regions. cGAS^+^ MN detection was carried out within these regions using the cell detection tool, focusing on the cGAS channel and applying area constraints of 0.25-7.5 μm^2^. The optimal cGAS intensity threshold for cGAS^+^ MN detection was determined for each tumor. cGAS^+^ MN detections in tumor cells were normalized to the total number of tumor cell detections, yielding a cGAS^+^ MN score (indicative of the degree of ongoing CIN) for each tumor.

#### CIN hotspot analysis

To map intratumoral regions with high cGAS^+^ MN densities (i.e. ‘CIN hotspots’), cGAS^+^ MN detections from cGAS–STING mIF panel-stained sections (manually annotated detections where available [n=11 tumors] or SEMi-automated detections where not [n=24 tumors]) were first spatially aligned and projected onto adjacent (4 μm) immune panel-stained section images. Tumors with ≤10 cGAS^+^ MN detections were omitted from the analysis.

Kernel Density Estimation (KDE), using the ‘kde2d’ function of the MASS package (v.7.3-60), was used to estimate the probability density function of cGAS^+^ MN across tumor images and resulting kernel densities were quantized into 1% bins (ranging from 0–100%) to yield relative density distributions for each tumor.

To assess enrichment of cell types within CIN hotspots, increasing density thresholds (in 1% increments, ranging from the 25–95^th^ percentile) were used to define hotspot regions. Fold-enrichment of cell types in CIN hotspots at each density threshold was calculated by comparing the relative proportion of a given cell type located inside versus outside the hotspot area. Tumors were excluded from analysis at a given threshold if no cells of a given class were present within either the hotspot or the remainder region.

#### Response-specific CIN associations

Linear interaction models were used to examine patient response-specific associations between intratumoral immune cell infiltration metrics or compartment-specific cGAS–STING pathway component (cGAS and STING) protein expression and tumoral cGAS^+^ MN frequencies. Patient response status to neoadjuvant treatment was inferred from histopathological response scores (tumor regression grade, TRG; also known as Mandard score). Patients with TRG 1-2 were categorized as neoadjuvant treatment (NA Tx) responders (R), whereas patients with TRG 4-5 were considered non-responders (NR). The linear model was defined as follows:

> Feature_(X)_∼cGAS^+^ MN x Response status

Where response variables (Feature_(x)_) consisted of inferred intratumoral immune cell abundances or compartment-specific cGAS–STING pathway component protein expression, and the cGAS^+^ MN predictor variable consisted of the tumor-level cancer-compartment specific cGAS^+^ MN frequency.

Interaction term p-values (cGAS^+^MN x Response status) and the differences between slopes of cGAS^+^ MN with queried features between response classes response class (slope NR – slope R) were extracted to identify features showing preferential scaling with cGAS^+^ MN in one response class over another.

### Multiplex imaging mass cytometry

FFPE slides were dewaxed for 2h at 58°C, followed by incubations in xylene and successive concentrations of 100%, 90%, 70% and 50% ethanol. Samples were subjected to heat-mediated epitope retrieval in Tris-EDTA (pH 9) at 98°C for 20 min. Tissues were then blocked with 3% BSA (A7906, Sigma) for 30 min at RT and stained overnight with the IMC antibody cocktail detailed in **Supplementary Table 2** at 4°C. The following day, excess antibody was washed off with TBS-T, nuclei were stained with DNA intercalator (201192A, Standard BioTools) at RT for 30 min, and slides were washed and left to dry, ready for laser ablation.

Image acquisition was performed using the Hyperion Fluidigm (Standard BioTools). Cell segmentation on resulting images was performed using the OPTIMAL pipeline^98^. Clustering of cell types was performed using the FCS Express software.

### *CXCL8* RNAscope

*In situ* detection of *CXCL8* mRNA transcripts in 4 µm FFPE human EAC tumor sections was carried out using the RNAscope assay (Advanced Cell Diagnostics) coupled to quantitative immunofluorescence on a Leica BOND Rx Autostainer, using a 20bp probe to the target region of *CXCL8* (2-1082, Advanced Cell Diagnostics). Tissue sections were pre-treated with heat and protease before hybridization with oligonucleotide probes. Detection and amplification were performed with the RNAscope Leica Multiplex Fluorescent Assay (Advanced Cell Diagnostics). Image analysis and *CXCL8* mRNA quantification was performed using QuPath software (v.0.5.0).

Briefly, tissue compartment (malignant and stromal) masks were designed using supervised machine learning pixel classifiers (Random Trees) bespoke to each tumor specimen, trained on mIF-stained sections (pCK and DAPI) adjacent (4 µm) to RNAscope sections. Compartment masks were aligned onto *CXCL8* mRNA images and images were thresholded to designate *CXCL8-*positive areas within tissue compartments. Resulting compartment-restricted *CXCL8*-positive areas were normalized to total tissue compartment areas as a percentage for each tumor.

### Transwell migration

#### Conditioned media generation

To generate cell-conditioned media (CM), EAC (OE33, SK-GT-4) and CP-A cells were seeded at a density of 3.0x10^5^ or 5.0x10^5^ cells, respectively, in 100 mm cell culture dishes. After 24h, cell media was either replaced with fresh media or, where indicated, replaced with 1 μM of the MPS1 inhibitor reversine (Cell Guidance Systems, SM85)- or DMSO-treated media for 24h, followed by drug wash-out and replacement with fresh serum-reduced (5% FBS) media. Cell culture media was conditioned for 48h before collection and cleared by centrifugation at 1,000 x g for 10 minutes at 4°C. For use in migration assays, conditioned media was further diluted with serum-free media (1:1 ratio) to a final concentration of 2.5% FBS. Migration controls included serum-free and unconditioned media (UM) as negative controls. UM was diluted with serum-free media, as described above. For positive controls, human recombinant IL-8 (Bio-Techne, 208-IL) or CCL2 (Proteintech, HZ-1334) were added to diluted UM.

#### CD14^+^ cell isolation

Human peripheral blood mononuclear cells (PBMCs) were isolated from leukapheresis cones from healthy donors (NHS Blood and Transplant, UK) obtained under ethical approval reference REC 19LO1848, by Ficoll-Paque density centrifugation. CD14^+^ monocytic cells were isolated using human CD14 MicroBeads (Miltenyi Biotec, 130-050-201) and LS autoMACS Columns (Miltenyi Biotec, 130-042-401) as per the manufacturer’s protocol.

#### Transwell migration assay

CD14^+^ monocytes (5.0x10^5^ cells in 100 μL UM) were added to the transwell inserts (6.5 mm transwells with 5.0 μm pores; Corning) and allowed to migrate for 24h towards CM or control media (600 μL) in the lower chambers. Where specified, CD14^+^ cells were incubated with 50 μg/mL CXCR1/2 inhibitor Reparixin (Selleck Chemicals) for 30 minutes before seeding in transwell inserts in the presence of Reparixin.

Migrated CD14^+^ cells in the lower chamber were imaged using a Celigo Image Cytometer (Nexcelom). At least two whole-well images were captured of each well and a custom ImageJ script was used to determine cell counts for each image. Migration indexes were calculated by normalizing cell counts to the corresponding control well counts.

### TCGA analyses

#### Transcriptomic data access

Transcripts Per Kilobase Million (TPM)-normalized and raw read RNA-sequencing counts for EAC tumors comprised in The Cancer Genome Atlas (TCGA)^99^ were accessed using the TCGAbiolinks R package (v.2.30.0)^100^. Raw read counts were filtered, retaining genes with a total read count of ≥ 75 counts across all samples and non-zero expression in > 25% of samples. Filtered genes were then DESeq2-normalized.

Raw read RNA-sequencing counts from TCGA stomach adenocarcinoma (STAD) were accessed as described above and filtered to comprise only chromosomally unstable STAD tumors (STAD-CIN), as defined by previous molecular subtyping by the TCGA Research Network^101^. Downstream filtering and normalization steps were performed as for EAC tumors.

#### Promoter methylation

Mean CpG-aggregated promoter methylation data for *CGAS* and *STING* from EAC tumor and adjacent esophageal tissue samples comprised in the TCGA were obtained through the SMART App web browser (available at http://www.bioinfo-zs.com/smartapp)^102^.

#### Survival analyses

Gene signature scoring of patient tumors was performed on DESeq2-normalized RNA-seq counts with the GSVA R package (v.1.5.0) using the ‘single-sample GSEA’ (ssGSEA) method^103^. Gene sets used for GSVA scoring are listed in **Supplementary Table 1.**

Survival data were accessed using the TCGAbiolinks R package. Patient cohorts were dichotomized using the maximally selected rank statistic (’Max-Stat’ method) with the maxstat package (v.0.7.25)^104^. Univariate survival analyses were performed using the Kaplan–Meier (log-rank) test with the survival (v.3.5.7) R package ^105^.

### Linear models

#### CIN score agreement

Linear associations between orthogonal measures of CIN (Aneuploidy, CIN^MN,^ and CIN^70^ CIN signature scores) in EAC and STAD-CIN tumors were computed using linear regression models, accounting for tumor purity and leukocyte fraction as covariates. Tumor purity estimates (obtained from the ESTIMATE algorithm^106^; Aneuploidy scores^30^ and leukocyte fraction calls were obtained from source publications^107^. CIN^70^ and CIN^MN^ signature scores were computed as described in the *‘Survival analyses’* subsection. The models were defined as follows:

> CIN score_(CIN70|Aneuploidy score)_∼CIN^MN^ + Purity + Leukocyte fraction

Resulting p-values between orthogonal CIN scores were extracted to infer the significance of CIN score interrelation.

#### CIN-associated expression

Linear regression models to evaluate which genes are most strongly associated with CIN scores across EAC tumors were computed using DESeq2-normalized gene expression data. CIN scores were computed as described above. Linear models accounting for tumor sample leukocyte fraction and tumor purity were iterated over all detected genes in filtered gene expression matrices:

> Expression_Gene X_∼CIN score + Purity + Leukocyte fraction

Where expression represents DESeq2-normalized expression for a given gene and CIN score represent either CIN^70^ score, CIN^MN^ score or Aneuploidy score. CIN-associated pathway enrichment was computed through GSEA, using p-values indicative of the significance of association between CIN scores and gene expression, as well as the associated slope estimates of associations to compute gene rankings. Gene ranks for GSEA were computed as follows: – *log_10_(p-value) x sign(slope)*.

#### cGAS–STING-dependent CIN-associated expression

A linear interaction model was used to identify genes whose expression scaled with CIN^MN^ score in EAC tumors, in a manner dependent on cGAS–STING expression. CIN^MN^ signature and cGAS– STING expression scores were computed as described in the *‘Survival analyses’* subsection. Tumor leukocyte fraction and tumor purity calls were included as covariates. Interaction models were defined as follows, and iterated across all detected genes within the filtered DESeq2-normalized gene expression matrix:

> Expression_Gene X_∼CIN^MN^ x cGAS–STING + Purity + Leukocyte fraction

The slopes and p-values of associations between CIN^MN^ (the predictor variable) and gene expression (the response variable) were extracted from model outputs to determine which genes demonstrate significant (positive or negative) scaling with CIN^MN^ score across tumors. Slopes and p-values of the interaction term were used to determine genes that exhibited a significantly different magnitude in association with CIN^MN^ score across different levels of cGAS–STING expression. Genes exhibiting significant positive associations (CIN^MN^ term p-value ≤0.05, positive slope) with CIN^MN^ and exhibited a significantly (interaction p-value ≤0.05) steeper slope across higher levels of cGAS–STING expression score were considered CIN-enriched in a cGAS–STING-dependent manner. Conversely, genes exhibiting significantly negative associations with CIN^MN^ (CIN^MN^ term p-value ≤0.05, negative slope) and exhibited a significantly (interaction p-value ≤0.05) more negative slope across higher levels of cGAS–STING expression score were considered CIN-depleted in a cGAS–STING-dependent manner.

Functional enrichment of genes exhibiting a significant positive relationship with CIN^MN^ in a cGAS– STING-dependent manner was performed using overrepresentation analysis.

### CCLE analyses

#### Data access

Cell line Aneuploidy scores and RPKM (reads per kilobase of transcript per million reads mapped)-normalized mRNA expression data were accessed through the DepMap portal (https://depmap.org/portal/)^108^. RPKM expression data were log_2_(+1)-normalized for visual comparison of relative expression levels between cell line types.

#### CIN score agreement

Linear associations between orthogonal CIN metrics (Aneuploidy score, CIN^MN^ and CIN^70^ signature scores) among cell lines of the Cancer Cell Line Encyclopedia (CCLE) were computed using linear regression models, accounting for cell lineage as a covariate. CIN^70^ and CIN^MN^ signature scores were computed with the GSVA package (ssGSEA method) using RPKM-normalized expression data. The models were defined as follows:

> CIN score^(CIN70 | Aneuploidy score)^∼CIN^MN^ + Lineage

Resulting p-values, indicative of the significance of association between CIN scores were extracted.

## Data availability statement

The patient-derived data that support the findings of this study will be available via the Translational Data Platform, Oxford Cancer Centre. Cell line transcriptomic data will be available via Gene Expression Omnibus (GEO).

## Acknowledgements

The authors gratefully acknowledge the patients and families of patients who contributed to this study. This work was funded by the Wellcome Trust (Clinical Career Development Fellowship to EEP, 224623/Z/21/Z); Cancer Research UK (CRUK) and the CRUK Oxford Centre (CRUK supported studentships to BB and SC; C2195/A31281; CTRQQR-2021\100002); the National Institute for Health and Care Research (NIHR) Oxford Biomedical Research Centre (BRC); the Medical Research Council (grant number MR/W006731/1. studentship to ASC); the Academy of Medical Sciences (Clinical Lecturer Starter Grant to EEP); the Oxford Experimental Cancer Medicine Centre (ECMC). DAM and JL are funded by a CRUK programme (DRCRPG-Nov22/1000007), the CRUK HUNTER Accelerator Award (grant number 175 A26813), and MRC project grant MR/Y003365/1. JL is also supported by an Academy of Medical Sciences Springboard award (SBF009\1103) and Royal Society Research Grant (RG\R2\232323). ER-G is funded by a Newcastle University Faculty of Medical Sciences PhD Studentship. This work has been supported by a Kleberg Innovative Investigator Award to KCA. This work was supported by the National Institute of Health (NIH), National Cancer Institute (NCI) grants, R37CA258829, R01CA266446, R01CA280414 and U54CA274506 to BI, a Pershing Square Sohn Cancer Research Alliance Award to BI, and a CRI Lloyd J. Old STAR Award to BI (CRI5579). This work was additionally supported by the Herbert Irving Comprehensive Cancer Center (HICCC) Human Tissue Immunology and Immunotherapy Initiative (P30CA013696). We acknowledge the contribution to this study made by the Oxford Centre for Histopathology Research and the Oxford Radcliffe Biobank, which are funded by the University of Oxford, the Oxford CRUK Cancer Centre and the NIHR CRN Thames Valley network. The views expressed are those of the authors and not necessarily those of the NIHR or the Department of Health and Social Care.

---

**Extended Data Figure 1.**
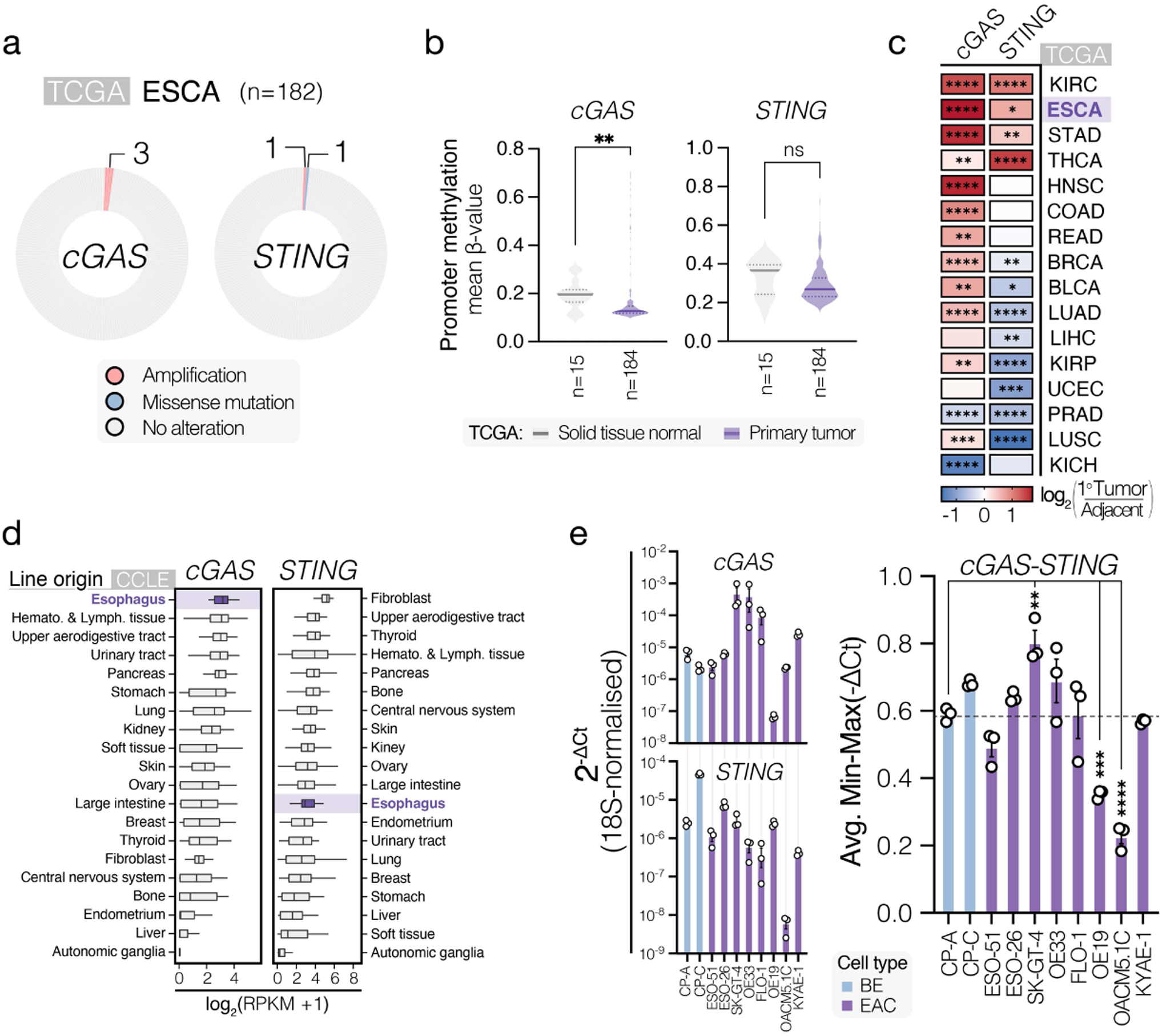
cGAS–STING expression across esophageal cancers and cell lines. (**a**) Pie chart showing the mutations and copy-number alterations in the genes encoding cGAS and STING across 182 surveyed esophageal tumors comprised in the TCGA database. (**b**) Violin plots of mean CpG-aggregated promoter methylation β-values levels in esophageal primary tumor samples (ESCA) and putatively normal adjacent tissue (NAT) samples (i.e. ‘solid tissue normal’) from the TCGA for genes encoding cGAS and STING. Significance was determined by Mann-Whitney U test. (**c**) Heatmap of log_2_(fold-change) differences between primary tumor and putatively healthy adjacent tissue sample mRNA abundance (TPM-normalized) of *cGAS* and *STING* for select solid tumor types comprised within the TCGA. Significance was determined by Mann-Whitney U test. (**d**) Tukey box plots of log_2_-normalized *cGAS* (left panel) and *STING* (right panel) mRNA expression (RPKM+1) levels for Cancer Cell Line Encyclopedia (CCLE) cell lines grouped by cell line origin. Box plots are plotted in descending order based on median expression values. Box plots for esophagus-derived cancer cell lines are highlighted in purple. (**e**) Quantitative reverse transcription (RT-qPCR) analysis of steady-state *cGAS* and *STING* mRNA expression levels across BE and EAC cell lines, normalized to the *18S* housekeeping gene. Average cGAS–STING expression represents the average of min-max-normalized -ΔCt values. Data from Barrett’s esophagus (BE) cell lines are shown in light blue, data from EAC lines are shown in purple. Bars are shown as the mean ± SEM from n=3 independent experiments and were analyzed by one-way ANOVA with FDR-based correction. **** p ≤0.0001, *** p ≤0.001, ** p ≤0.01.

**Extended Data Figure 2.**
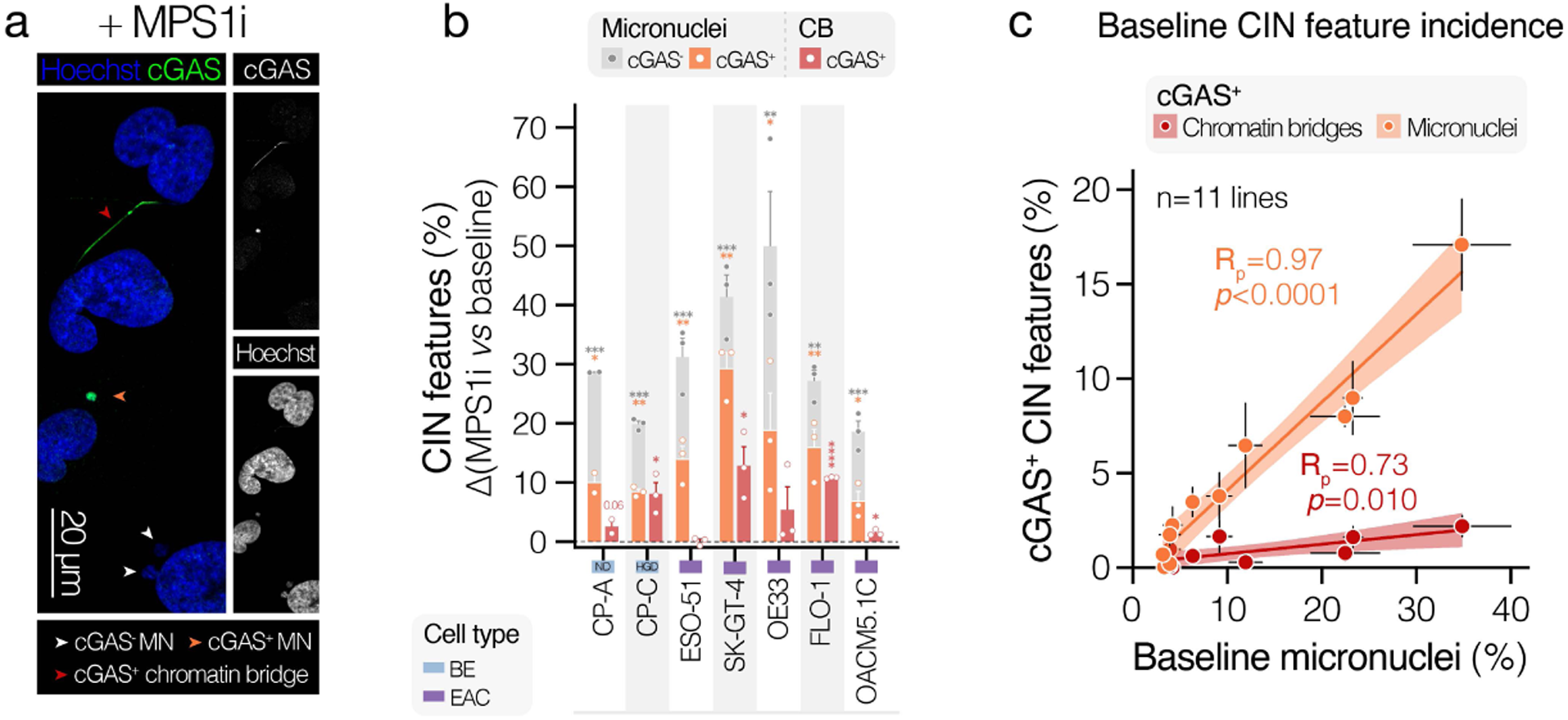
Chromosomal instability features in esophageal adenocarcinoma cell lines. (**a**) Representative confocal microscopy image of SK-GT-4 cells following treatment with the MPS1 inhibitor reversine (0.5 μM) for 48h, comprising examples of scored CIN features, including cGAS^-^ and cGAS^+^ micronuclei, as well as cGAS^+^ chromatin bridges. Cells were stained with anti-cGAS and Hoechst (DNA). The image represents a maximum-intensity projection of a Z-stack taken at 1 μm intervals. Scale bar corresponds to 20 μm. (**b**) Difference in frequency (number of features / 100 cells) of scored CIN features between MPS1 inhibitor-(0.5 μM, 48h) and DMSO-treated cells. Barrett’s esophagus (BE). Bars represent the mean ± SEM of n=2 (CP-A) or n=3 (all other lines) independent experiments, with ∼≥100 cells counted per experiment. Data analyzed by two-sided unpaired t-test. (**c**) Scatter plots of baseline cGAS^+^ chromosomal instability-associated feature (cGAS^+^ micronuclei and cGAS^+^ chromatin bridges) frequencies versus baseline micronuclei frequencies across BE and EAC cell lines. Dots represent the mean ± SEM of n=3 independent experiments, with ∼≥100 cells counted per experiment. Shown are estimated simple linear regression lines, 95% confidence intervals, Pearson correlation coefficients (R_p_) and p-values from Pearson correlation analyses. **** p ≤0.0001, *** p ≤0.001, ** p ≤0.01, * p ≤0.05.

**Extended Data Figure 3.**
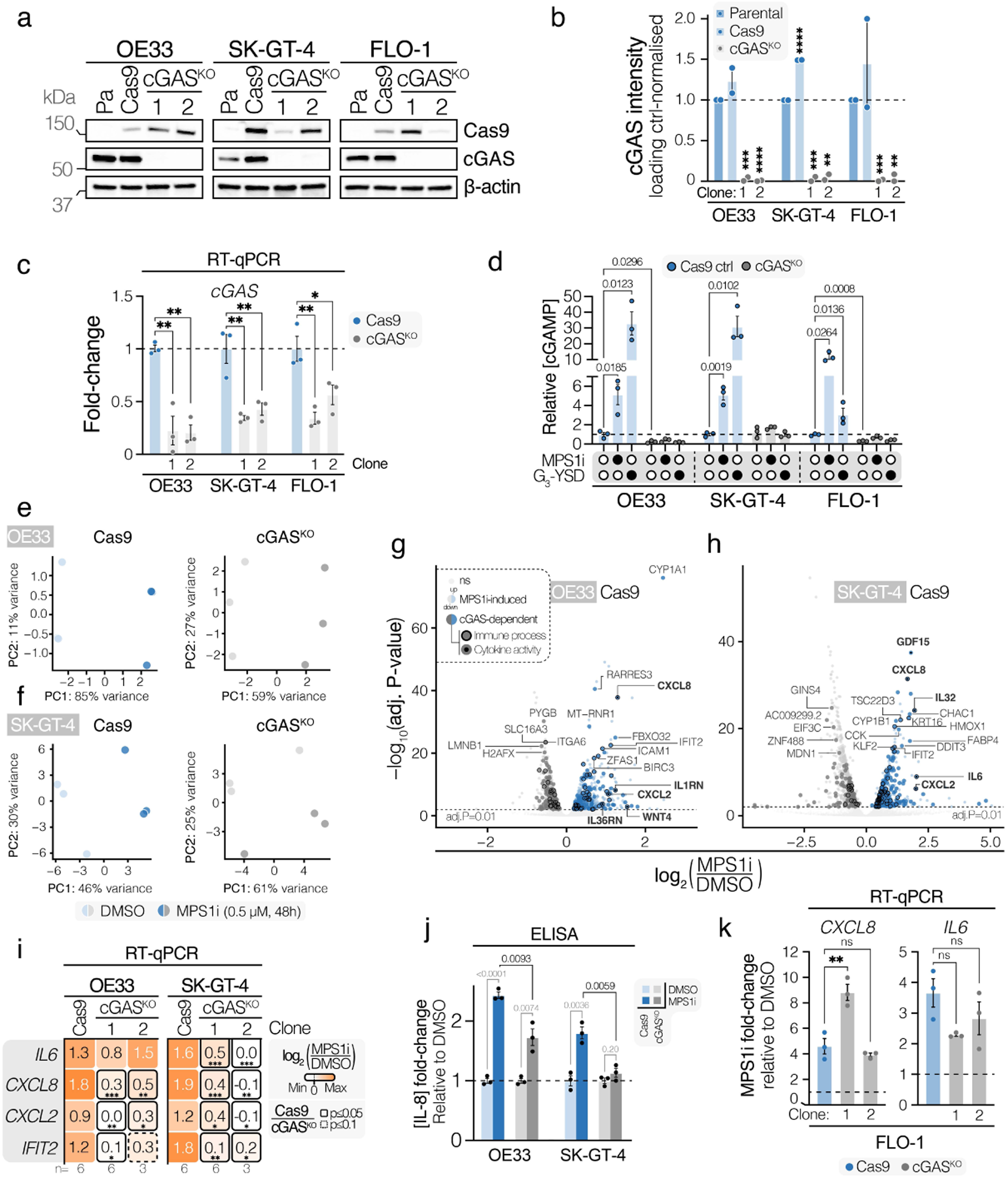
cGAS knockout and CIN-driven cGAS-dependent target validation in esophageal adenocarcinoma cells. (**a**) Representative immunoblot of OE33, SK-GT-4 and FLO-1 cells showing depletion of cGAS protein expression in cGAS^KO^ clones compared to parental and Cas9 (empty-vector) control cells and Cas9 expression in Cas9 control and cGAS^KO^ clones, but not parental cell lines. β-actin is used as a loading control. Data are representative of n=2 independent experiments. (**b**) Densitometry analysis of cGAS protein levels in OE33, SK-GT-4 and FLO-1 cells, showing suppression of cGAS protein in cGAS^KO^ clones, but not Cas9 control and parental cells. Band intensities have been normalized to loading controls and parental cell cGAS protein intensity for each experiment. Bars are shown as the mean ± SEM from n=2 independent experiments and were analyzed by ANOVA with FDR correction for each cell line. (**c**) Quantitative reverse transcription (RT-qPCR) analysis of baseline *cGAS* mRNA expression levels across OE33, SK-GT-4 and FLO-1 Cas9 and cGAS^KO^ clones. CGAS Ct values are normalized to the *18S* housekeeping gene using the ΔCT method and are expressed relative to normalized Cas9 control cGAS expression levels. Bars are shown as the mean ± SEM from n=3 independent experiments and were analyzed by ANOVA with FDR correction for each cell line. (**d**) Relative extracellular concentrations of 2’3’-cGAMP (determined through 2’3’-cGAMP ELISA) across OE33, SK-GT-4 and FLO-1 cGAS^KO^ (clone #1 for each) and Cas9 control clones, showing enhanced cGAMP production in cGAS-proficient (Cas9 control) clones upon CIN induction (MPS1i) and DNA transfection (G_3_-YSD), but not in cGAS^KO^ clones (clone #1 used for each line). Cells have either been pulse-treated with DMSO or 1 μM MPS1i (reversine) for 24h, 48h prior to media collection, or transfected with 2.5 μg/mL Y-form DNA (G_3_-YSD) 24h prior to collection. 2’3’-cGAMP concentrations have been normalized to absolute cell counts ([cGAMP] / 10^6^ cells) at treatment endpoint and are expressed relative to concentrations in DMSO-treated Cas9 control cells for each cell line. Bars are shown as the mean ± SEM from n=3 independent experiments and were analyzed by ANOVA with FDR correction for each cell line. (**e**, **f**) Principal component analysis of transcriptomes of DMSO and MPS1i (0.5 μM, 48h)-treated Cas9 and cGAS^KO^ clones of the (**e**) OE33 cell line and (**f**) SK-GT-4 line. (**g**, **h**) Volcano plots of differential gene expression analysis in (**g**) OE33 and (**h**) SK-GT-4 cGAS-proficient Cas9 control cells between MPS1i (reversine)-treated (0.5 μM, 48h) and control (DMSO)-treated cells. Genes highlighted in dark blue or dark grey are significantly more upregulated or downregulated (paired t-test p ≤0.05) in Cas9 versus cGAS^KO^ cells (i.e. ‘cGAS-dependent’) upon treatment. Genes related to the immune system (’Immune System Process’-annotated) are highlighted with a border; genes with known cytokine activity (mapping to ‘Cytokine activity’ GO term, GO:0005125) are highlighted with a central dot. (**i**) Heatmap of quantitative reverse transcription (RT-qPCR) analysis of the immune targets *IL6*, *CXCL8*, CXCL2 and *IFIT2* in OE33 and SK-GT-4 Cas9 and cGAS^KO^ clones treated for 48h with DMSO or 0.5 μM reversine (MPS1i). Fold-changes were derived using the 2^-ΔΔCt^ method, normalizing expression values to *18S* expression and corresponding DMSO control treatment gene expression. Color maps to log_2_-transformed fold-changes (FC) between MPS1i-treated cells and DMSO-treated cells. Boxes with solid borders correspond to clones with a significantly different log_2_(FC) for that gene compared to its corresponding Cas9 control FC value. Values inside heatmaps represent the mean log_2_(FC) for a gene across n=3 or n=6 independent experiments and were analyzed independently by ANOVA with FDR correction, comparing log_2_(FC) induction of each gene in cGAS^KO^ clones to that of its respective Cas9 control clone. (**j**) IL-8 concentrations (determined by ELISA) in conditioned media of cells pulse-treated with DMSO or 1 μM reversine (MPS1i) for 24h, followed by 48h media conditioning in the absence of treatment. IL-8 concentrations have been normalized to respective DMSO-treatment concentrations for each genotype. Grey and blue shades correspond to Cas9 and cGAS^KO^ genotypes, respectively. Darker shades correspond to MPS1i-treatment, whereas lighter shades correspond to DMSO treatment. Bars represent the mean ± SEM from n=3 independent experiments and were analyzed by ANOVA with FDR correction for each cell line. (**k**) Quantitative reverse transcription (RT-qPCR) analysis of *IL6* and *CXCL8* mRNA fold-inductions upon MPS1i versus DMSO control treatment in Cas9 and cGAS^KO^ FLO-1 cells. Fold-changes were derived using the 2^-ΔΔCt^ method, normalizing expression values to *18S* expression and DMSO control treatment gene expression levels. Bars are shown as the mean ± SEM from n=3 independent experiments and were analyzed by ANOVA with FDR correction for each gene. **** p ≤0.0001; *** p ≤0.001; ** p ≤0.01; * p ≤0.05; ns, not significant.

**Extended Data Figure 4.**
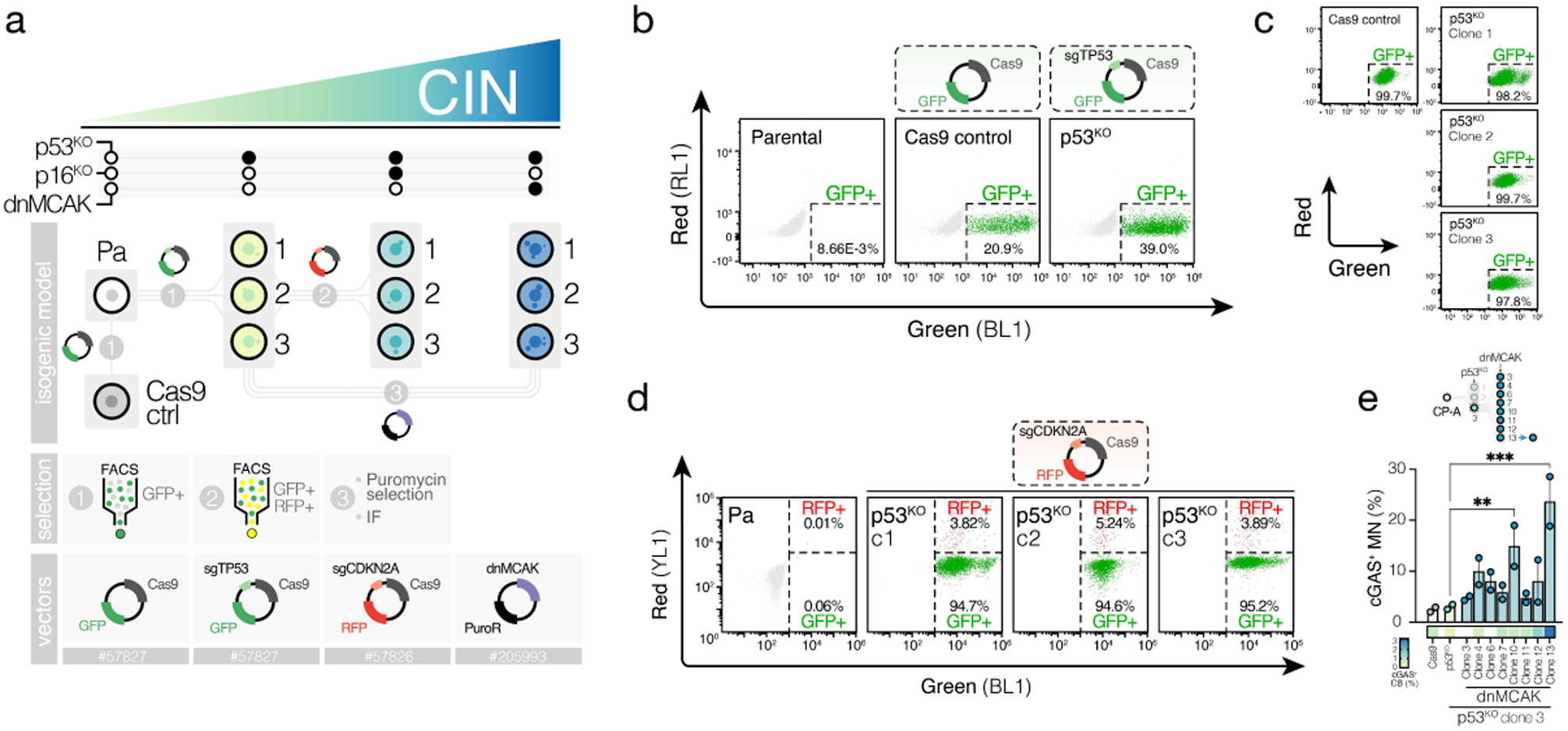
Generation of an isogenic non-dysplastic Barrett’s esophagus cell line model of variable chromosomal instability. (**a**) Schematic of the experimental strategy used to generate an isogenic cell line model with varying levels of CIN, starting with an hTERT-immortalized, non-dysplastic BE-derived CP-A founder. Parental (Pa) CP-A cells were sequentially altered to disrupt intrinsic barriers to chromosomal instability, through CRISPR-Cas9-mediated targeting of *TP53* (encoding p53) and *CDKN2A* (encoding p16) and disruption of mitotic checkpoints through overexpression of a dominant-negative mutant form of the mitotic regulator MCAK (dnMCAK). A Cas9 control (Cas9 ctrl) clone, expressing the pL-CRISPR.SFFV.eGFP, was generated to control for confounding effects associated with Cas9 overexpression. Clonal selections were performed through single-cell sorting, sorting GFP^+^ cells (p53^KO^) and GFP^+^RFP^+^ cells (p53p16^DKO^s), as well as through puromycin selection (p53^KO^dnMCAKs, expressing the puromycin resistance [PuroR] gene) and limiting dilution, followed by immunofluorescence-based screening of micronucleation rates. Cloning was performed in triplicate, with three independent clones for every altered genotype (aside from Cas9 control cells) to avoid clone-specific confounding effects. Vector codes correspond to Addgene plasmid numers. (**b**) Flow cytometric profiling of untransduced parental CP-A cells and CP-A cells transduced with CRISPR.SFFV.eGFP (Cas9 mixed population) and CRISPR.SFFV.eGFP.sgTP53 (p53^KO^ mixed population) lentiviral vectors. Cells were gated on the GFP^+^ population for single-cell sorting. (**c**) Flow cytometric profiling of expanded Cas9 and p53^KO^ single-cell clones, gated on GFP^+^ as in (**a**), showing a universal uptake of CRISPR.SFFV.eGFP vectors among selected clones. (**d**) Flow cytometric profiling of untransduced parental CP-A cells and p53^KO^ clones transduced with CRISPR.SFFV.tRFP (p53p16^DKO^ mixed population) lentiviral vectors. Cells were gated on the indicated GFP^+^RFP^+^ population for single-cell sorting. (**e**) Example of IF-based screening of candidate p53^KO^dnMCAK single-cell clones (derived from p53^KO^ clone 3) for CIN^high^ clones, showing quantifications of cGAS^+^ chromatin bridges (heatmap) and cGAS^+^ micronuclei. Bars represent the mean of cGAS^+^ MN frequencies ± SEM of n=2 technical replicates (two independent slides per clone, seeded in parallel), with ∼≥100 cells counted per experiment. Data were analyzed by ANOVA with FDR correction, comparing all candidate clones to the Cas9 control clone. *** p ≤0.001, ** p ≤0.01.

**Extended Data Figure 5.**
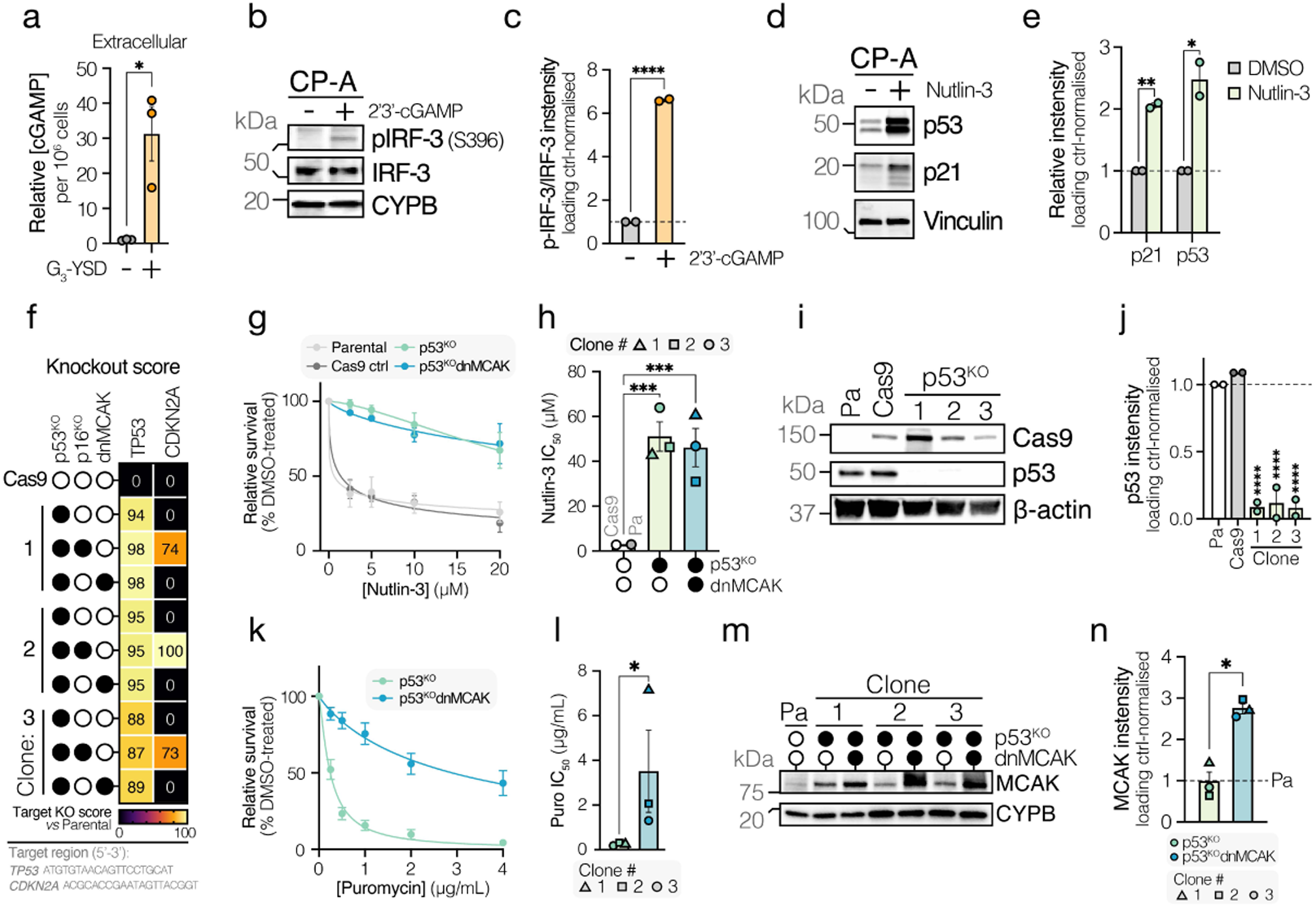
Validation of an isogenic non-dysplastic Barrett’s esophagus model founder line and derived genotypes. (**a**) Relative extracellular concentrations of 2’3’-cGAMP (determined by ELISA) in CP-A cells that were mock transfected or transfected with 2.5 μg/mL G_3_-YSD 24h prior to sample collection. 2’3’-cGAMP concentrations have been normalized to absolute cell counts ([cGAMP] / 10^6^ cells) at treatment endpoint and are expressed relative to concentrations in mock-transfected CP-A cells. Bars are shown as the mean ± SEM from n=3 independent experiments and were analyzed by two-sided unpaired t-test. (**b**) Immunoblot of CP-A cells mock transfected or transfected with 10 μg/mL exogenous 2’3’-cGAMP, showing increased activating phosphorylation of IRF3 at Ser396 upon cGAMP treatment. Data are representative of n=2 independent experiments. CYPB is used as a loading control (**c**) Densitometric quantification of (**b**), showing the ratio between phosphorylated IRF3 (Ser396) and total IRF3 protein abundance. Band intensities have been normalized to CYPB abundance and are expressed relative to normalized mock transfection band intensities. Bars are shown as the mean ± SEM of n=2 independent experiments and were analyzed by two-sided unpaired t-test. (**d**) Immunoblot of CP-A cells treated with diluent (DMSO) or 10 μm Nutlin-3 for 24h, showing p53 and p21 accumulation in CP-A cells, indicative of a proficient p53 pathway. Vinculin (VCL) is used as a loading control. Data are representative of n=2 independent experiments. (**e**) Densitometric quantification of (**d**), showing p53 and p21 accumulation in CP-A cells upon Nutlin-3 treatment. Band intensities have been normalized to loading control abundance, expressed relative to normalized DMSO band intensities. Bars are shown as the mean ± SEM of n=2 independent experiments and were analyzed by two-sided unpaired t-test. (**f**) Heatmap of *TP53* and *CDKN2A* sgRNA target site knockout scores (Synthego ICE tool) for Cas9, p53^KO^, p53p16^DKO^ and p53^KO^dnMCAK CP-A clones. Parental sequences have been used as a reference. Target sites are shown. (**g**) Titration of Nutlin-3 in a 5-day viability (WST-8) assay for parental, Cas9 control, p53^KO^ and p53^KO^dnMCAK CP-A cells. Data are from n=3 independent experiments, with n=3 technical replicates per experiment. Data are represented as mean ± SEM. Genotype-level averages have been pooled for visualization purposes. (**h**) Quantification of Nutlin-3 IC_50_ values from IC_50_ dose-response curves in (**g**). Bars represent the mean ± SEM of indicated genotypes. Datapoints represent the mean of n=3 independent experiments, with n=3 technical replicates per experiment. Datapoint shapes represent clone numbers. Significance was tested by one-way ANOVA with Tukey’s HSD. (**i**) Immunoblot of parental, Cas9 control and p53^KO^ CP-A cells, showing depletion of p53 in p53^KO^ clones and Cas9 expression in Cas9 control and p53^KO^ cells. Data are representative of n=2 independent experiments. β-actin is used as a loading control. (**j**) Densitometry analysis of (**i**). Band intensities are normalized to the loading control and parental cell band intensities. Bars represent the mean ± SEM. Significance was tested by one-way ANOVA with Tukey’s HSD. (**k**) Titration of puromycin in a 3-day viability (WST-8) assay for p53^KO^ and p53^KO^dnMCAK CP-A cells. Data are from n=3 independent experiments, with n=3 technical replicates per experiment. Data are represented as mean ± SEM. Genotype-level averages have been pooled for visualization purposes. (**l**) Quantification of puromycin IC_50_ values from IC_50_ dose-response curves in (**k**). Bars represent the mean ± SEM of indicated genotypes. Datapoints represent the mean of n=3 independent experiments, with n=3 technical replicates per experiment. Datapoint shapes represent clone numbers. Significance was tested by two-tailed ratio paired t-test. (**m**) Immunoblot of parental, p53^KO^ and p53^KO^dnMCAK CP-A cells, showing elevated MCAK expression in p53^KO^dnMCAK clones. Data are representative of n=3 independent experiments. CYPB is used as a loading control. (**n**) Densitometry analysis of (**m**). Band intensities were normalized to the loading control and parental cell band intensities. Bars represent the mean ± SEM. Significance was tested by two-tailed paired t-test. **** p ≤0.0001; *** p ≤0.001; ** p ≤0.01; * p ≤0.05.

**Extended Data Figure 6.**
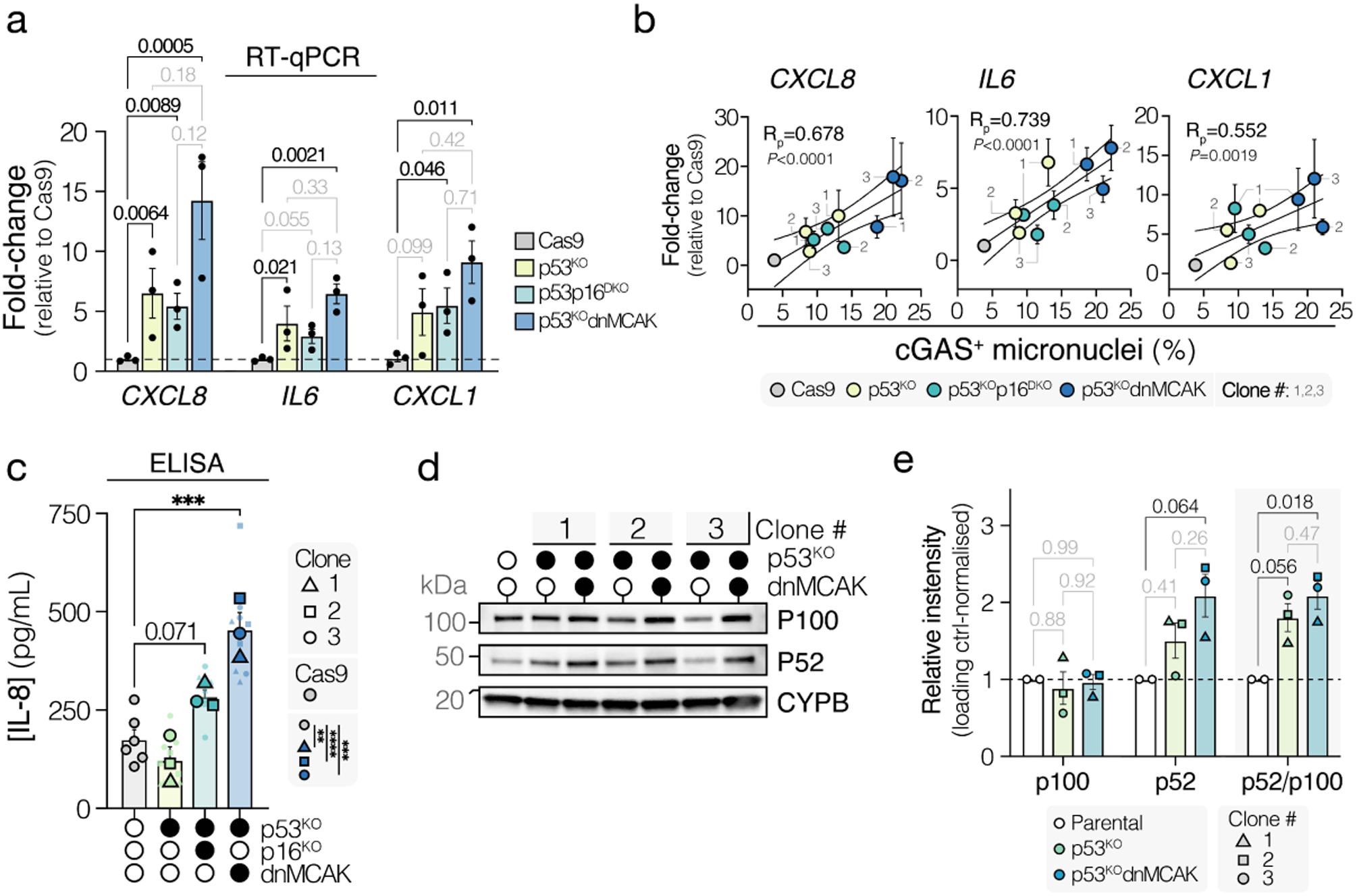
Validation of CIN-associated immune targets in an isogenic non-dysplastic Barrett’s esophagus cell line model. (**a**, **b**) Quantitative reverse transcription (RT-qPCR) analysis of baseline expression of the immune targets *IL6*, *CXCL8*, and *CXCL1* among CP-A cell model lines. Fold-changes were derived using the 2^-ΔΔCt^ method, normalizing expression values to *18S* expression and expression values in Cas9 control cells. (**a**) Bars for p53^KO^, p53p16^DKO^, p53^KO^dnMCAK genotypes represent the mean of n=3 independent clones ± SEM, where each point represents the mean of n=3 independent experiments for a given clone. Bars for Cas9 cells represent the mean ± SEM of n=3 independent experiments. Data were analyzed by ANOVA with FDR correction, comparing fold-changes in p53^KO^, p53p16^DKO^ and p53^KO^dnMCAK genotypes to Cas9. Statistical analysis was performed on log_2_-transfomed data. (**b**) Linear relationship between baseline cGAS^+^ MN burden for CP-A-derived cell lines and qPCR-derived relative expression levels of immune targets. Each point and its associated error bars represent the mean ± SEM of n=3 independent qPCR experiments (y-axis) and n=3 independent IF experiment (x-axis). Clone numbers are indicated. Pearson correlation analysis coefficients (R_p_) and p-values are shown. Statistical analysis was performed on log_2_-transfomed data. (**c**) IL-8 concentrations (determined by ELISA) in conditioned media (conditioned for 48h) of CP-A-derived cell lines. Bars for p53^KO^, p53p16^DKO^, p53^KO^dnMCAK genotypes represent the mean value across n=3 independent clones ± SEM, where each point represents the mean of n=3 independent experiments for a given clone. The bar for Cas9 cells represents the mean ± SEM of n=6 independent experiments. Datapoint shape indicates clone number for p53^KO^, p53p16^DKO^ and p53^KO^dnMCAK clones. Data were analyzed by ANOVA with FDR correction, comparing genotype-level IL-8 concentrations in p53^KO^, p53p16^DKO^ and p53^KO^dnMCAK genotypes to Cas9. Significant ANOVA p-values for individual clones (not pooled by genotype) are shown in the sidebar. *** p ≤0.001. (**d**) Immunoblot of CP-A cell lines showing increased abundance of the non-canonical NF-κB pathway components p100 and p52. CYPB was used as a loading control. Data are representative of n=2 independent experiments. (**e**) Densitometry analysis of (**d**). Band intensities are normalized to the loading control and parental cell band intensities. The ratio of p52 to p100 normalized protein abundance is shown. Bars for p53^KO^ and p53^KO^dnMCAK genotypes represent the mean of n=3 independent clones ± SEM, where each point represents the mean of n=2 independent experiments for a given clone. Bars for Cas9 cells represent the mean ± SEM of n=2 independent experiments. Data were analyzed by ANOVA with Tukey’s HSD.

**Extended Data Figure 7.**
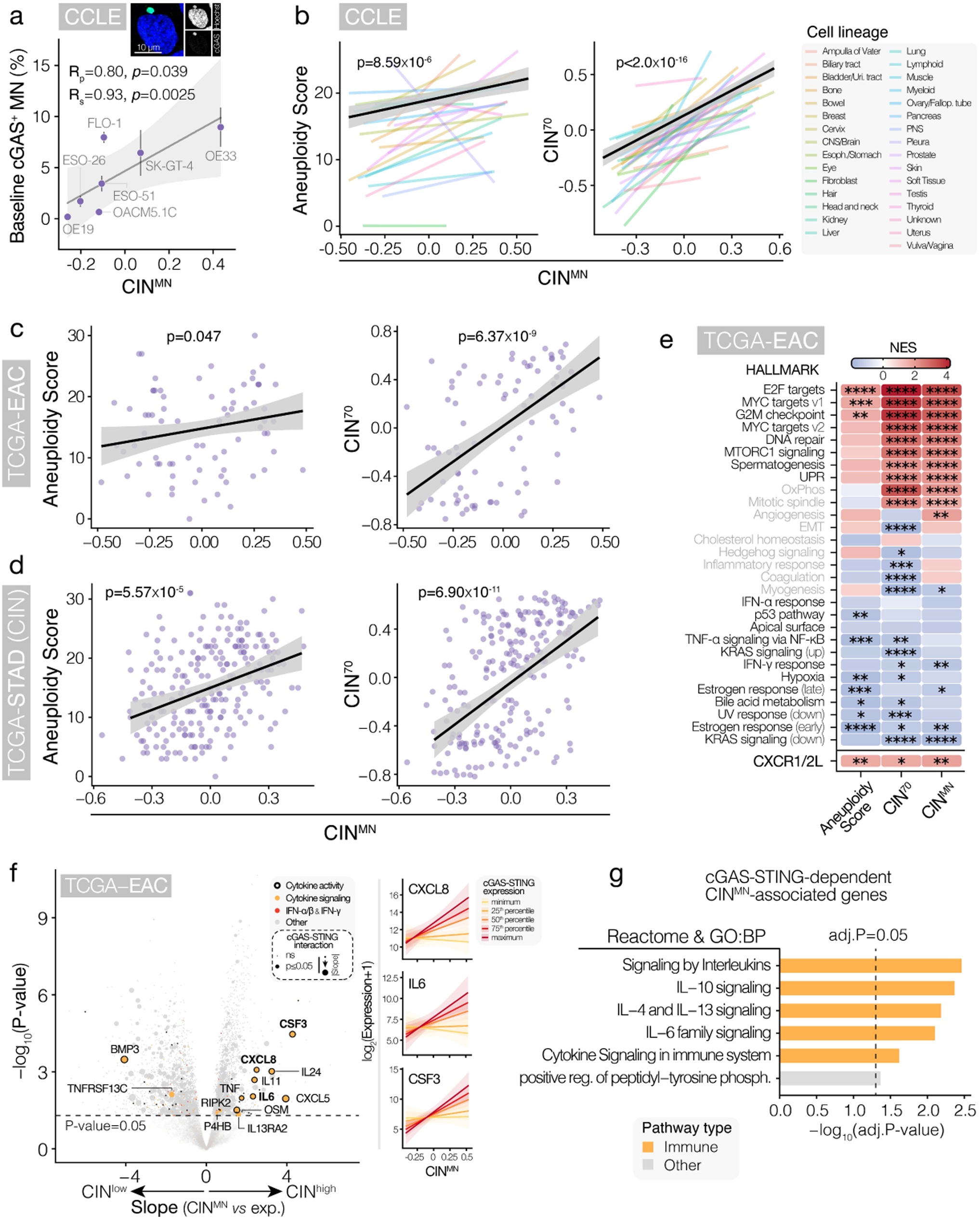
Validation of a novel transcriptional signature of chronic ongoing chromosomal instability in esophageal cells. (**a**) Scatter plot of microscopy-derived baseline cGAS^+^ MN frequencies (exemplar image shown) versus CIN^MN^ scores (derived from CCLE RNA-sequencing data) for indicated EAC cell lines. The simple linear regression line, 95% confidence intervals, Pearson and Spearman correlation coefficients (R_p_ and R_s_, respectively), and p-values are shown. (**b**) Linear association between CIN^MN^ score and orthogonal metrics of CIN (CIN^70^ signature score and Aneuploidy score) across cell lines of the Cancer Cell Line Encyclopedia (CCLE). Colored lines represent simple linear regression lines in specific cell lineages. The significance of the association between CIN^MN^ and other CIN scores is derived using a linear model accounting for cell lineage as a covariate. The black regression line and grey 95% confidence intervals correspond to the marginal effect estimates of CIN^MN^ on CIN score over different cell lineages. (**c**, **d**) Linear association between CIN^MN^ score and orthogonal metrics of CIN (CIN^70^ signature score and aneuploidy score) across (**c**) EAC tumors and (**d**) chromosomally unstable stomach adenocarcinoma (STAD-CIN) tumors comprised in The Cancer Genome Atlas (TCGA). The significance of the association between CIN^MN^ and other CIN scores is derived using a linear model accounting for tumor purity and leukocyte fraction as covariates. The black regression line and grey 95% confidence intervals correspond to the marginal effect estimates of CIN^MN^ on CIN score over covariates. (**e**) Heatmap of gene set enrichment analysis (GSEA) outputs, showing MSigDB Hallmark gene sets that are most strongly commonly enriched with orthogonal CIN metrics (Aneuploidy score, CIN^70^, CIN^MN^) in EAC tumors from the TCGA. GSEA of a CXCR1/2 ligand (CXCR1/2L; *CXCL1-3*, *CXCL5-8*) signature shows a positive enrichment across all three CIN metrics. **** p ≤0.0001; *** p ≤0.001; ** p ≤0.01; * p ≤0.05. (**f**) Volcano plot showing outputs of a linear model looking at the association between CIN^MN^ score and gene expression that are conditional on cGAS–STING expression (whilst accounting for leukocyte fraction and tumor purity as confounders) in EAC tumor from the TCGA. Slopes and p-values of genes indicate the strength of association with CIN^MN^ across EAC tumors. Genes that have an additional dependence (i.e. a statistically significant interaction; interaction term p ≤0.05) on cGAS–STING expression are highlighted through size. Genes annotated with the ‘Cytokine signaling’ gene ontology term (GO:0019221) are highlighted in orange. Genes with known IFN-α/β (R-HSA-909733) or IFN-γ (R-HSA-877300) pathway involvement are highlighted in red. Genes with reported cytokine activity (GO:0005125) are marked by a border. The right panel includes examples of linear model predictions of the relationship between select genes (*CXCL8*, *IL6*, *CSF3*) across multiple levels of tumoral cGAS–STING expression. (**g**) Overrepresentation analysis (using Reactome and Gene Ontology: Biological Process pathway sets) of cGAS-dependent CIN-associated genes (i.e. genes that scale significantly more strongly in high cGAS– STING expression settings) showing a statistical overrepresentation of inflammatory pathways.

**Extended Data Figure 8.**
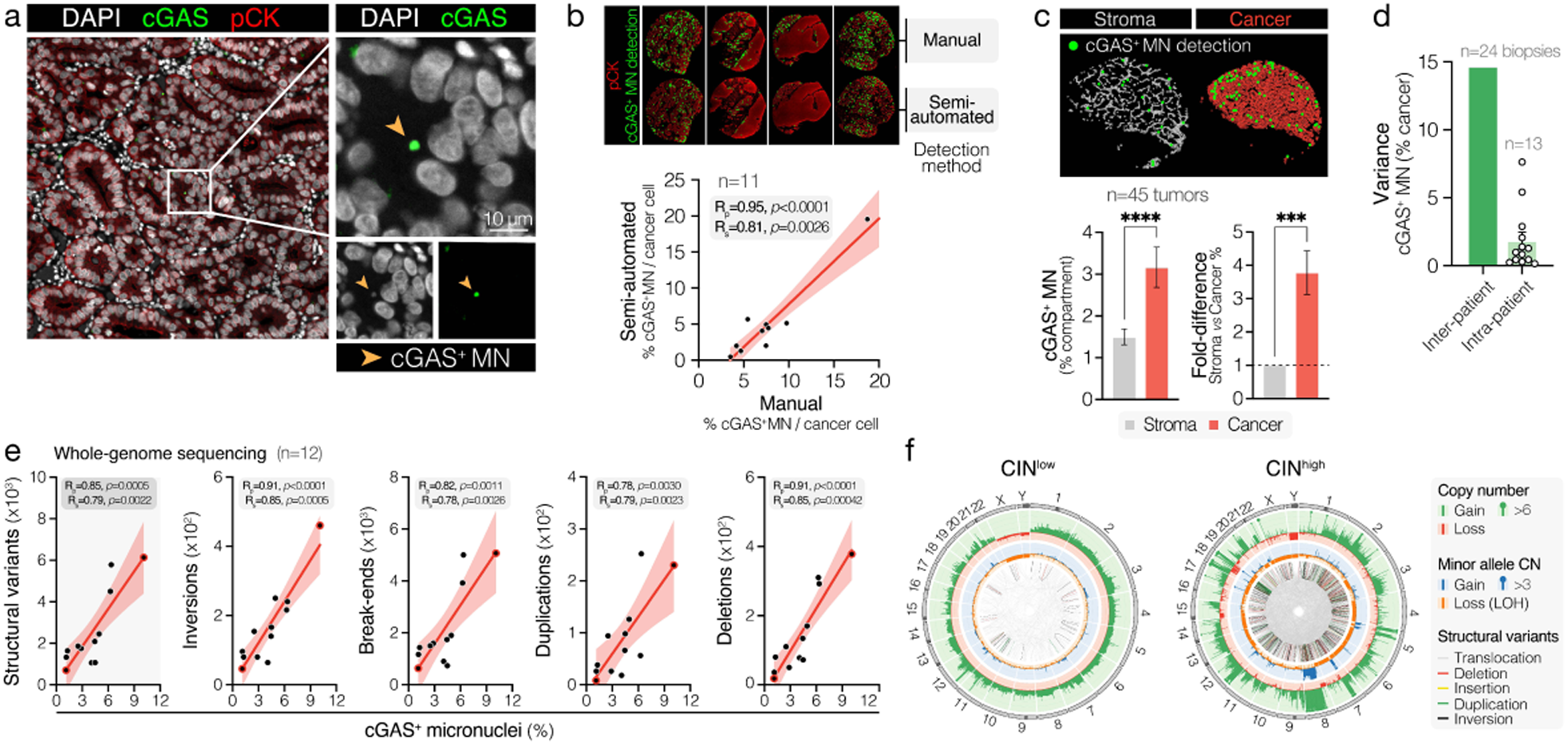
Detection of cGAS^+^ micronuclei as a measure of ongoing cGAS-activating chromosomal instability in human EAC tumors. (**a**) Representative high-resolution image of a human EAC tumor biopsy specimen stained with DAPI (DNA), anti-cGAS Ab and anti-pan-cytokeratin (pCK) Ab showing selective localization of cGAS at micronuclei. Scale bar corresponds to 10 μM. (**b**) *Upper panel:* Examples of manual and semi-automated tumoral cGAS^+^ MN detections. *Lower panel:* Scatter plot of manual cGAS^+^ MN quantifications (’ground truth’) versus quantifications obtained using a semi-automated detection approach across n=11 EAC tumor specimens, showing high concordance between methods. Detections were limited to the cancer cell compartment and were normalized to the total number of detected cancer cells to obtain cancer compartment-specific cGAS^+^ MN frequencies. The simple linear regression line, 95% confidence intervals and Pearson and Spearman correlation coefficients (R_p_ and R_s_, respectively) and p-values are shown. (**c**) *Upper panel:* Example of semi-automated detections in the stromal versus cancer cell compartment of a primary EAC tumor. *Left lower panel:* Bar plots of semi-automated cGAS^+^ MN detections in cancer versus stromal compartments across n=45 patient tumors, showing a higher preponderance of micronuclei in malignant compartments. *Right lower panel:* Bar plots showing the fold-difference in cGAS^+^ MN detection frequencies in cancer compartments relative to corresponding stromal compartments. Bars represent the mean ± SEM for n=45 human EAC tumors. Data were analyzed by Wilcoxon matched-pairs signed-rank test, pairing malignant and stromal values from the same tumors. **** p ≤0.0001; *** p ≤0.001. (**d**) Barplot comparing inter- and intra-patient variances in tumoral cGAS^+^ MN frequencies, showing lower variance among samples derived from the same patient (e.g. patients with matched pre-treatment biopsies and post-treatment resections) compared to pre-treatment samples from different patients. Bars represent the mean ± SEM. (**e**) Scatter plots of the observed tumoral cGAS^+^ MN frequency versus whole-genome sequenced (WGS) - derived structural variant (SV) loads across n=12 primary EAC tumor specimens. The simple linear regression line, 95% confidence intervals and Pearson and Spearman correlation coefficients (R_p_ and R_s_, respectively) and p-values are shown. Datapoints used as exemplars in (**f**) are highlighted with a red border. (**f**) Circos plots showing copy number alterations (CNAs) and SVs for representative EAC tumors exhibiting the highest and lowest measured cGAS^+^ MN frequencies across all sequenced tumors (CIN^high^ and CIN^low^, respectively). The outermost circle shows chromosomes, with darker shading representing gaps in the reference human genome (e.g. centromeres, heterochromatin and missing short arms). The second circle shows tumor purity-adjusted copy number (CN) changes, with gains shown in green and losses shown in red. Absolute copy numbers > 6 are shown with a green dot. The third shell represents minor allele CNs, with gains and losses shown in blue and orange, respectively. Minor allele CN > 3 are highlighted with a blue dot. The innermost circle displays intra- and inter-chromosomal SVs. SV categories are highlighted by color, as indicated.

**Extended Data Figure 9.**
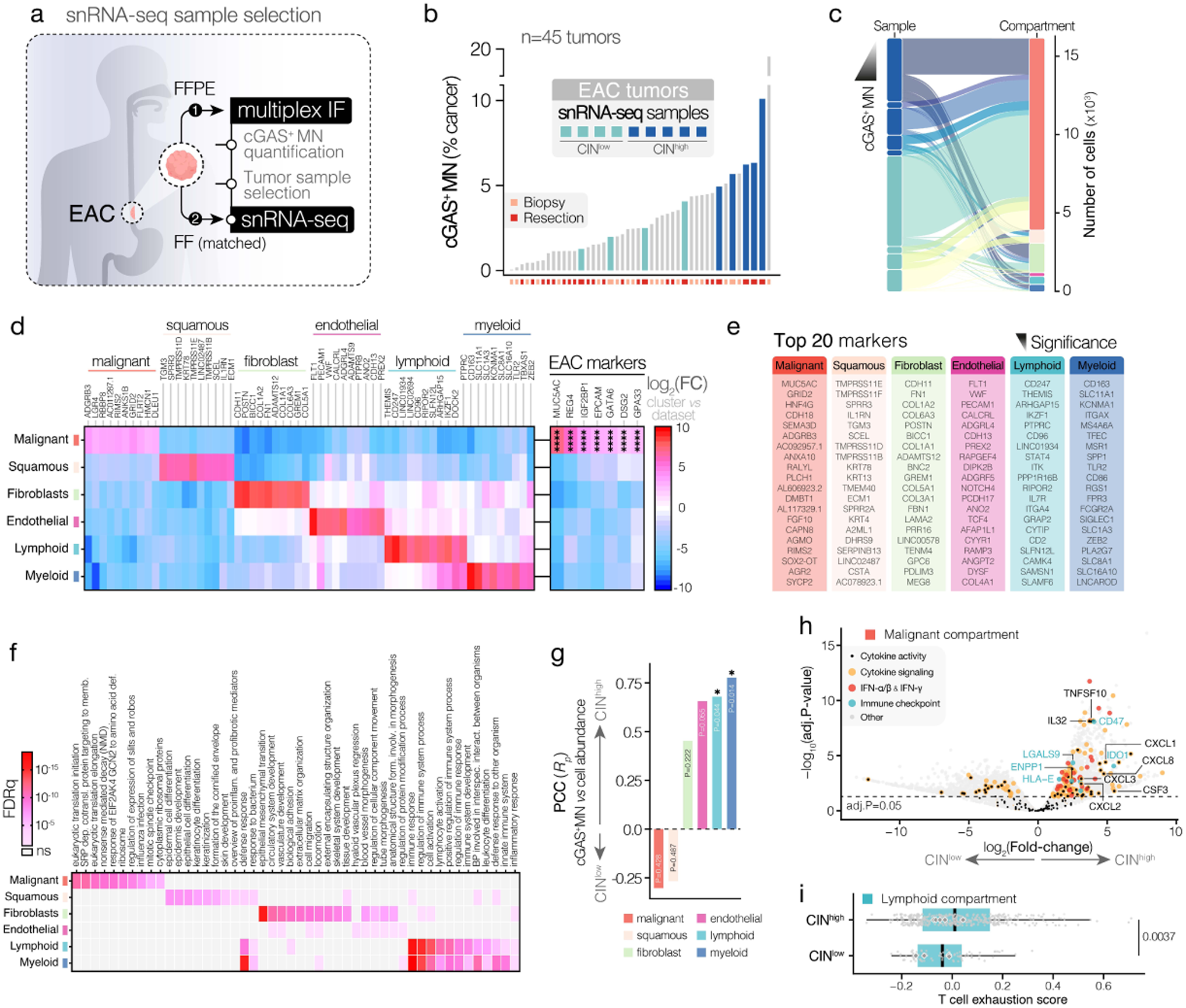
Single-nucleus RNA sequencing of human EAC tumors. (**a**) Experimental strategy used to shortlist samples for single-nucleus RNA-sequencing (snRNA-seq). Matched fresh frozen (FF) and formalin-fixed paraffin-embedded (FFPE) tissue specimens were obtained for tumors. FFPE tissues were stained with a multiplex immunofluorescence (mIF) panel using antibodies targeting cGAS and pan-cytokeratin (pCK), as well as DAPI (DNA) to evaluate the range of CIN across n=45 patient tumors. Tumors for snRNA-seq were shortlisted to span the observed spectrum of CIN across all queried tumors. SnRNA-seq was performed on matched FF tissue. (**b**) Multiplex immunofluorescence (mIF)-derived cGAS^+^ MN frequencies across n=45 patient tumors, including treatment-naïve staging laparoscopy biopsies and surgical resection specimens. (**c**) Sankey plot showing the contribution (number of cells) of each sample to identified cell compartments. Samples are ranked in descending order according to their mIF-inferred cGAS^+^ MN burden. (**d**) Heatmap showing the top 10 most significant highly expressed markers associated with each of the identified cell clusters. Color maps to the log_2_-transformed fold-change of a given marker in a cell cluster versus all other cell clusters. Select well-established EAC markers are indicated, showing high specific expression in the malignant compartment. (**e**) The top 20 most significantly enriched markers in each cluster, ranked from top to bottom by significance of enrichment. Unlike (**c**), shown markers are not limited to genes with abundant expression (i.e. minimum 1 count / cell). (**f**) Heatmap of the top 10 most significantly positively enriched pathways in each cluster, showing cluster specific pathway activities aligned with cluster identities. Pathway enrichment was performed using gene set enrichment analysis (GSEA) on ‘FindAllMarkers’ differential expression analysis outputs for each cluster. Color maps to the –log_10_-transformed significance of GSEA enrichment (FDRq). Cluster-enriched pathways are ranked from left to right within each cluster by significance of enrichment. (**g**) Bar plots of the Pearson correlation coefficients (R_p_) of the relative abundance of each cell compartment (the percentage of cells of a given identity relative to all cell detection within a sample) versus the tumoral cGAS^+^ MN burden across samples. P-values of Pearson correlation analyses are shown. (**h**) Volcano plot showing differential gene expression between malignant cells of CIN^high^ and CIN^low^ snRNA-seq samples. Genes annotated with the ‘Cytokine signaling’ gene ontology term (GO:0019221) are highlighted in orange. Genes with known IFN-α/β or IFN-γ (R-HSA-909733 or R-HSA-877300, respectively) pathway involvement are highlighted in red. Genes with reported cytokine activity (GO:0005125) are marked by a black central dot. Common tumor-expressed checkpoints (see **Supplementary Table 1**) are highlighted in teal. The dashed line indicates a Benjamini-Hochberg-adjusted p-value of 0.05. (**i**) Box plots of T cell exhaustion scores in lymphoid cells from CIN^high^ and CIN^low^ snRNA-seq samples. T cell exhaustion scores were computed using the ‘AddModuleScore’ function of the Seurat package using a literature-curated T cell exhaustion signature (**Supplementary Table 1**). Boxes represent the median ± interquartile range and whiskers were plotted using Tukey’s method. Mean scores for each sample are plotted. Significance was determined by Mann-Whitney U test. **** p ≤0.0001; * p ≤0.05.

**Extended Data Figure 10.**
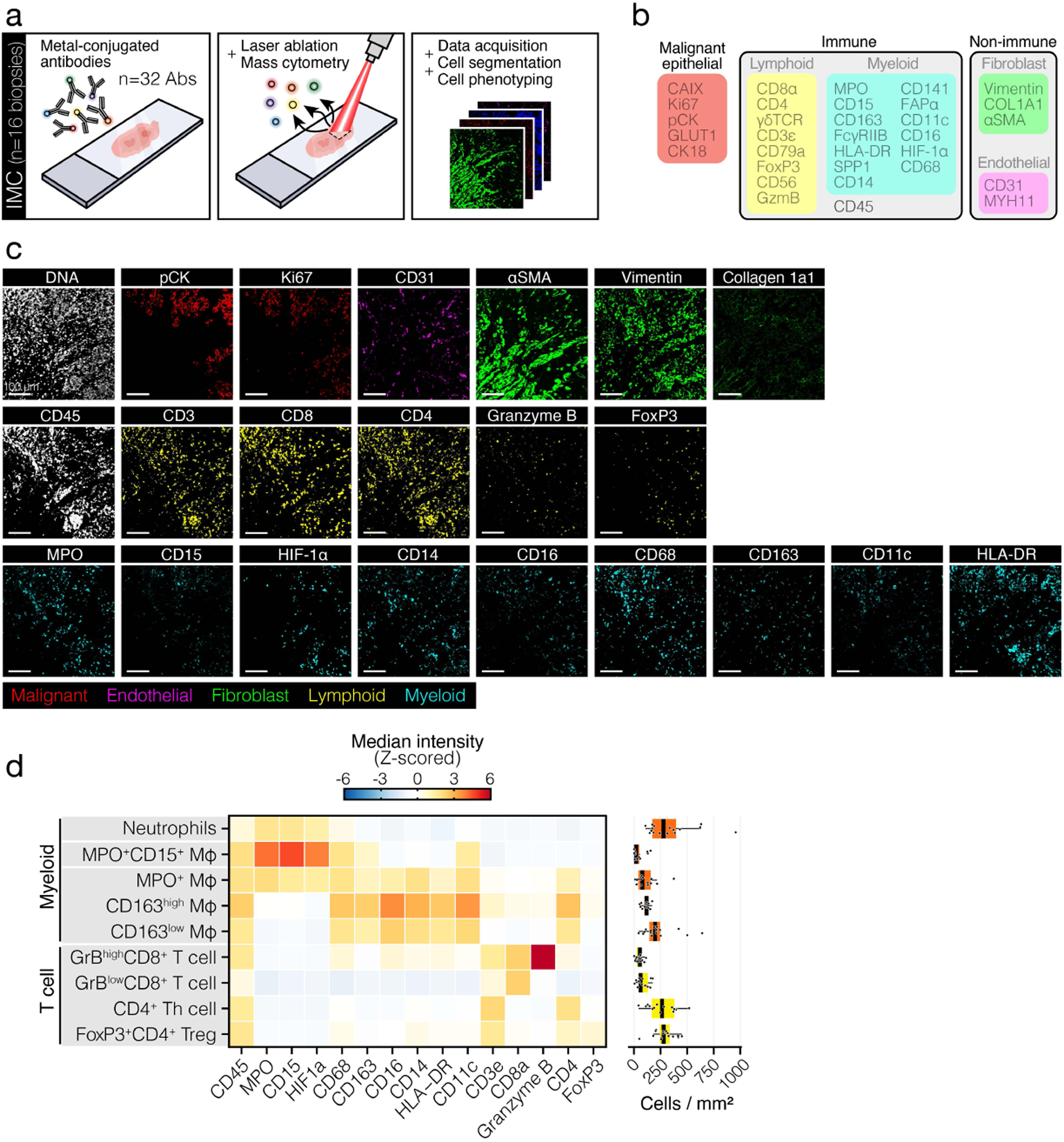
Imaging mass cytometry-based profiling of the chromosomally unstable EAC tumor immune landscape. (**a**) Multiplexed imaging mass cytometry (IMC) workflow applied across n=16 pre-treatment human EAC tumors. (**b**) Schematic of the IMC antibody panels used for human EAC tumor microenvironment phenotyping, showing the targets used to differentiate malignant epithelial, immune (lymphoid and myeloid) and non-immune stromal (fibroblasts and endothelial cells) cells. (**c**) Representative images of various malignant, immune and non-immune stromal cell IMC markers in a pre-treatment human EAC tumor region. Scale bars correspond to 100 μM. (**d**) *Left panel:* Heatmap of median Z-scored (on a per sample basis) intensities of markers from the immune panel across identified immune cell subtypes. *Right panel:* Box plots of cell densities of identified immune cell subtypes across n=16 primary EAC biopsies. Boxes represent the median ± interquartile range, whiskers were plotted using Tukey’s method.

**Extended Data Figure 11.**
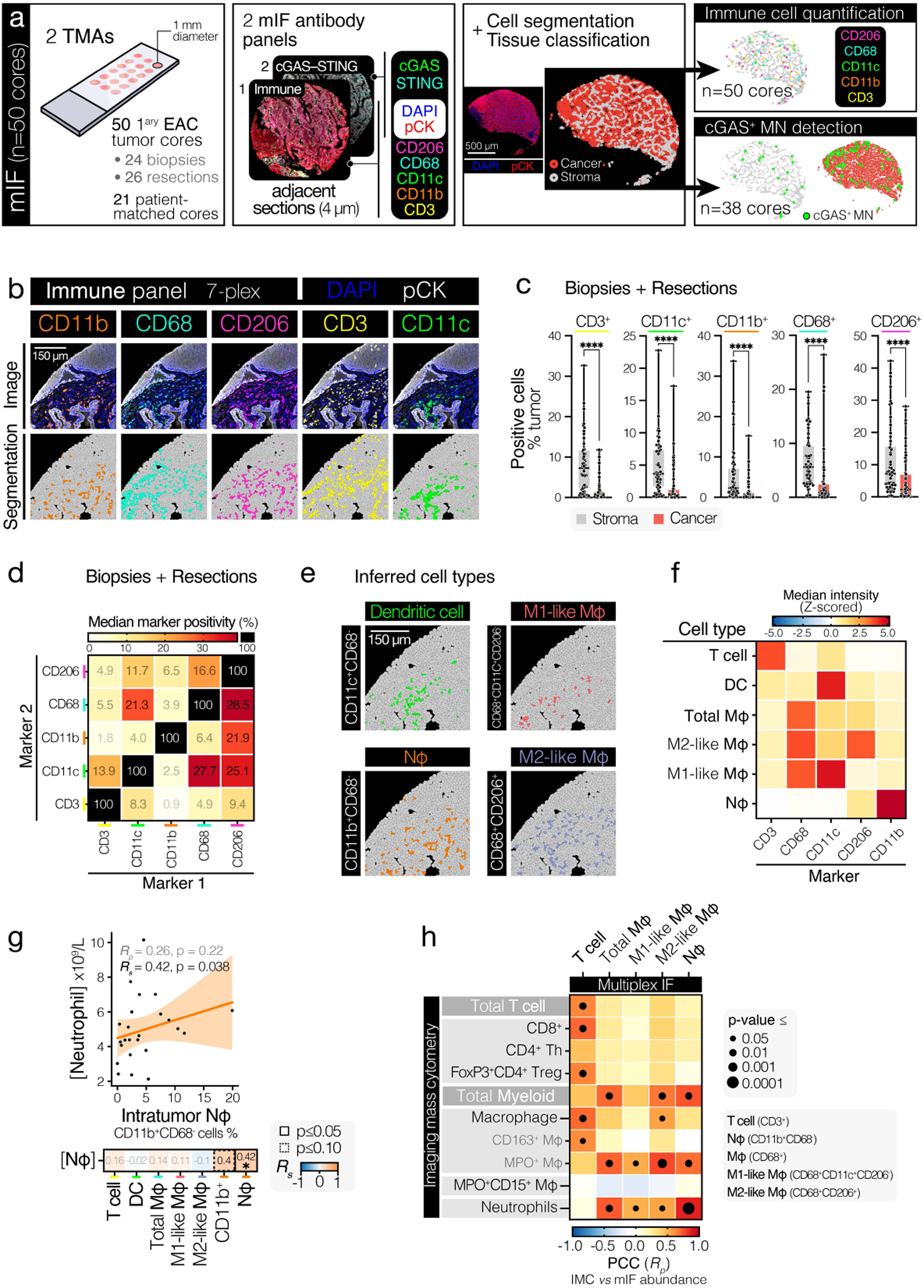
Multiplex immunofluorescence-based profiling of the chromosomally unstable EAC tumor immune landscape. (**a**) Multiplex immunofluorescence workflow. (**b**) *Image*: Representative high-resolution image of a human EAC tumor biopsy specimen stained with DAPI (DNA), as well as anti-pan-cytokeratin (pCK), anti-CD11b, anti-CD68, anti-CD206, anti-CD3 and anti-CD11c antibodies (Abs; 7-plex staining), showing broadly stromal staining patterns for immune cell markers. *Segmentation*: Representative cell segmentations showing immune marker-positive cell detections after setting appropriate intensity thresholds. Scale bar corresponds to 150 μM. (**c**) Box plots showing the number of stromal compartment versus cancer compartment immune marker-positive cell detections (as a percentage of all cell detections in a tumor) across all analyzed biopsy and resection specimens, showing a predominance of immune marker-positive cells in stromal compartments. Boxes represent the median ± interquartile range. Box plot whiskers range from minimum to maximum. Data were analyzed by Wilcoxon matched-pairs signed-rank test, pairing malignant and stromal detections from the same tumors. (**d**) Coincidence heatmap of the degree of marker positivity co-occurrence across all detected cells in human EAC biopsy and resection specimens, showing a pronounced co-occurrence (e.g. CD68 and CD206) or mutual exclusivity (e.g. CD11b and CD3) between some immune markers. Color maps to the median marker positivity across all tumors. (**e**) Representative cell segmentations in an EAC biopsy tumor showing multi-marker cell detections indicative of inferred cell types. Scale bar corresponds to 150 μM. (**f**) Heatmap of median Z-scored (on a per sample basis) intensities of markers from the mIF immune panel across identified immune cell subtypes. (**g**) Scatter plot of the abundance of infiltrating neutrophils (Nϕ) in pre-treatment tumors versus concurrently collected blood-circulating neutrophil counts [Nϕ], showing a positive association. The heatmap shows spearman correlations coefficients (R_s_) of associations between circulating neutrophil counts and all detected intratumoral cell types, showing an association specific to the Nϕ compartment. Heatmap color maps to R_s_. R_s_ values of correlations are shown inside the boxes. Pairwise correlations with a p ≤0.05 are highlighted with a solid black border, whereas correlations with a p ≤0.10 are highlighted with a dashed black border. The linear regression line, 95% confidence interval, R_s_ and Spearman correlation p-value of the association between Nϕ abundance and circulating [Nϕ] are shown on the scatter plot. (**h**) Correlation heatmap showing high inter-correlations between immune cell type abundances inferred through imaging mass cytometry (IMC)-based immunophenotyping and through mIF across n=16 human EAC tumors analyzed using both methods. Color maps to the Pearson correlation coefficient (R_p_). Significant associations in the abundance between cell types inferred through different methods are highlighted with a central black dot. Dot size maps to the magnitude of significance, as indicated. **** p ≤0.0001; * p ≤0.05.

**Extended Data Figure 12.**
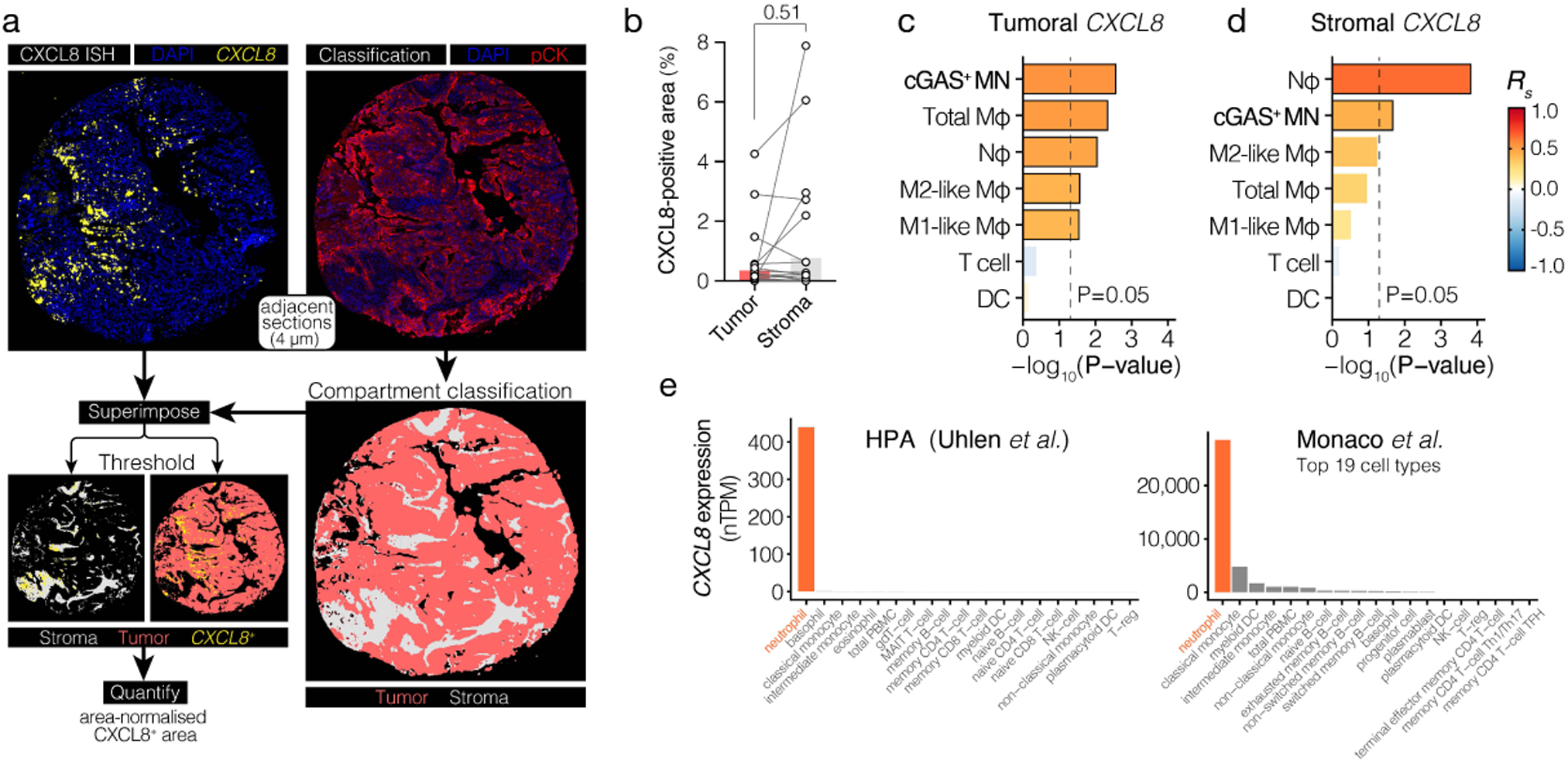
Tumor cell-intrinsic and stromal *CXCL8* mRNA expression in human EAC tumors. (**a**) Workflow to quantify malignant and stromal compartment-specific *in situ CXCL8* expression in human EAC tumors from *CXCL8* RNAscope images. (**b**) Tumoral (cancer cell-intrinsic) versus stromal *CXCL8* expression across n=29 EAC tumors. Bars represent the mean and lines between datapoints link tumor and stromal values for a given sample. Data were analyzed by Wilcoxon matched-pairs signed-rank test, pairing malignant and stromal *CXCL8* expression from the same tumors. (**c**, **d**) Bar plots of Spearman correlations of the association between (**c**) Tumoral or (**d**) stromal *CXCL8* expression and CIN (tumoral cGAS^+^ MN frequency) or intratumoral immune cell abundance. Color maps to Spearman correlation coefficient (Rs). The x-axis corresponds to the –log_10_-transformed Spearman correlation p-value. The dashed line corresponds to a p-value of 0.05. (**e**) *CXCL8* mRNA expression level across peripheral blood mononuclear cell (PBMC)-derived immune cell subtypes in the Human Protein Atlas^33^ and the Monaco *et al.*^34^ datasets.

**Extended Data Figure 13.**
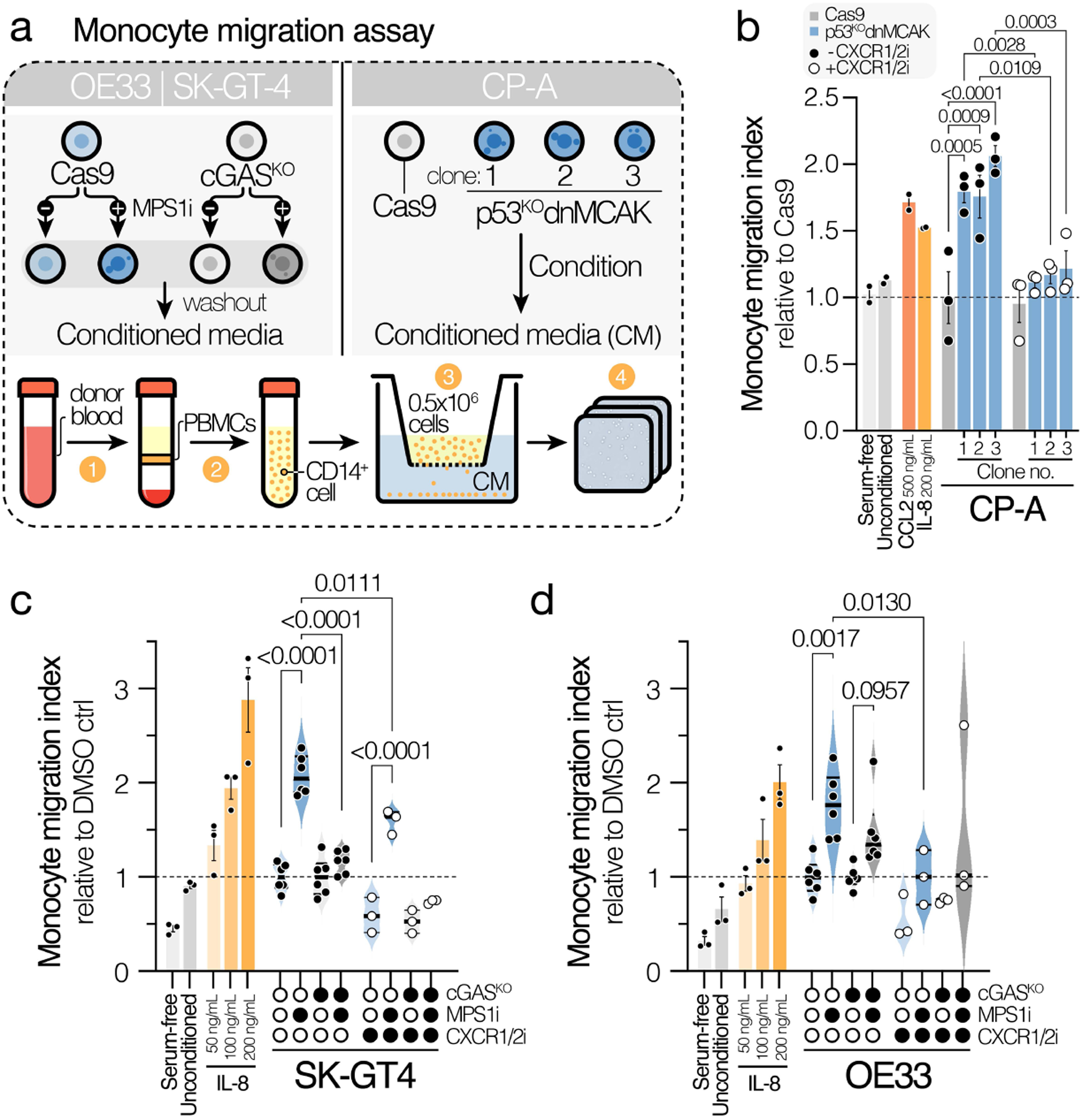
Chromosomal instability-driven cGAS–STING as a monocyte-attracting cue. (**a**) Schematic illustrating the experimental strategy for monocyte migration assays. For EAC cells, Cas9 and cGAS^KO^ cells were exposed to 1 μM MPS1i (reversine) for 24h, followed by extensive wash-out to avoid drug carry-over. Cells were then allowed to condition media for 48h prior to conditioned media (CM) collection. For CP-A cells, cells were covered in fresh untreated media and left to condition media for 48h. CD14^+^ monocytes were positively selected from healthy donor blood-derived peripheral blood mononuclear cells (PBMCs). Migration assays were performed using transwell migration chambers, allowing monocytes to migrate towards CM for 24h before whole-well imaging based quantification. (**b**) Transwell assays were performed as described in (**a**). Data were normalized to the untreated Cas9 control. Data of n=3 independent experiments are shown. Significance was tested by one-way ANOVA with FDR-correction. (**c**, **d**) Transwell assays were performed as described in (**a**), using (**c**) SK-GT-4 cell conditioned media (CM) or (**d**) OE33 CM. Migration indexes were normalized to respective DMSO controls. Data of n=6 independent experiments are shown for experimental samples and n=3 for migration control samples. Monocytes were derived from n=2 independent healthy donors. Significance was tested by one-way ANOVA with FDR-correction.

**Extended Data Figure 14.**
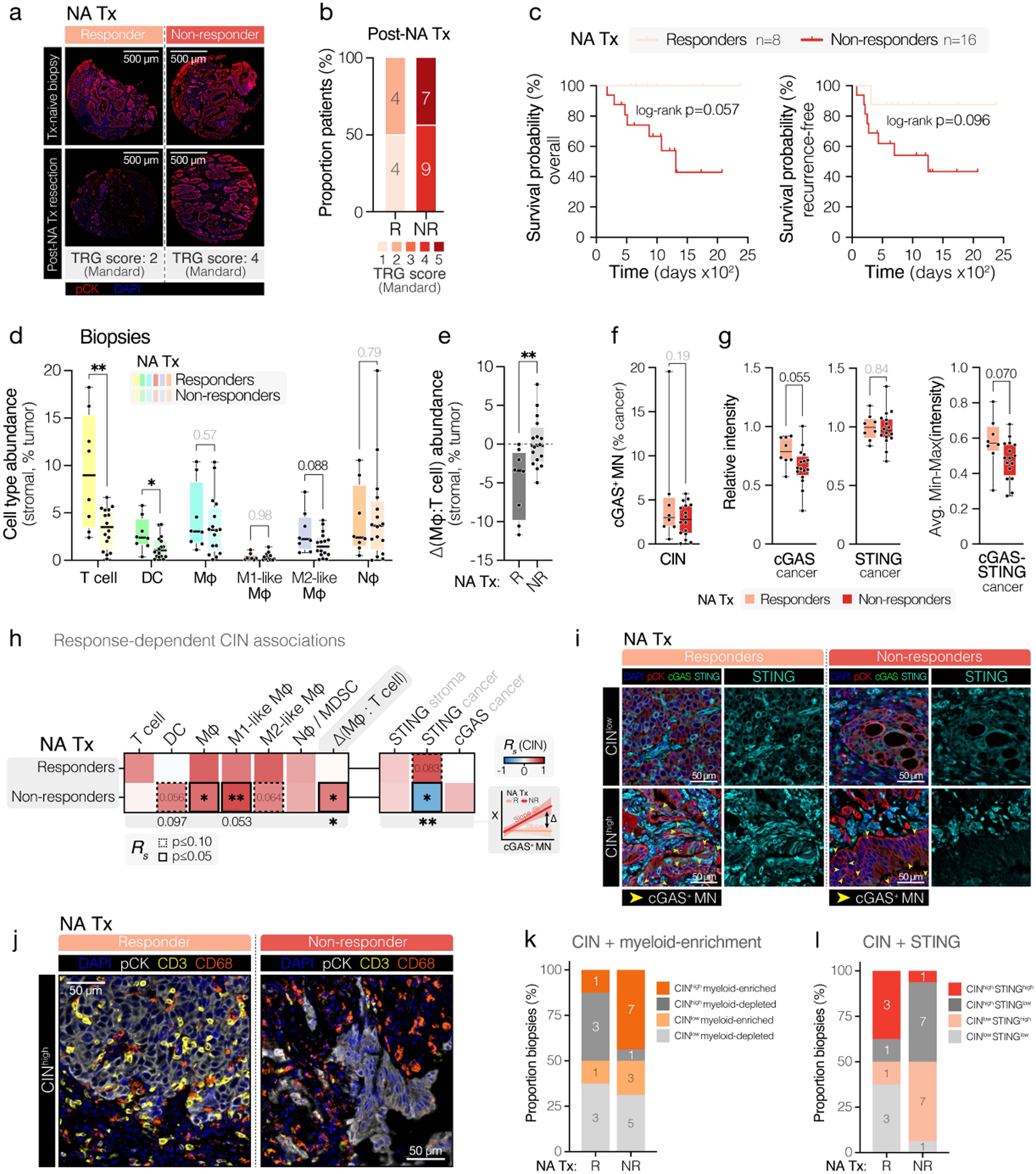
EAC patient response-associated tumor immune features. (**a**) Representative high-resolution images of matched human EAC tumor biopsy and resection specimens from neo-adjuvant treatment (NA Tx) responders and non-responders, stained with DAPI (DNA) and anti-pan-cytokeratin (pCK) Ab, showing poor histopathological response in the non-responder group. Tumor regression grade (TRG; also known as Mandard score) is indicated for each patient. Scale bar corresponds to 500 μM. (**b**) Bar plots showing the number of patients classed as NA Tx histopathological responders (R) or non-responders (NR) and proportion of tumors classed as TRG 1-5. (**c**) Kaplan–Meier plots of overall survival (left panel) and recurrence-free survival (right panel) of NA Tx responders and non-responders. The significance of the difference between patient survival was determined using the log-rank test. (**d**) Box plots of immune cell abundances (immune cell detections in the stromal compartment as a % of all cell detections in the tumor) for all queried immune cell types in biopsies of neoadjuvant treatment (NA Tx) responders versus non-responders. (**e**) Box plot showing the relative proportion between intratumoral macrophages (Mφ; CD68^+^) and T cells (CD3^+^; Δ[Mφ: T cell]), in biopsy tumors of NA Tx responders versus non-responders. (**f**, **g**) Box plots of (**f**) tumoral cGAS^+^ MN frequencies and (**g**) cGAS–STING pathway component expression in biopsy tumors of NA Tx responders versus non-responders. (**d**–**g**) Boxes represent the median ± interquartile range. Box plot whiskers range from the 10^th^-90^th^ percentile. Data were analyzed via unpaired two-tailed t-test. (**h**) Heatmap of Spearman correlations between cGAS^+^ MN frequency and immune cell abundances or cGAS–STING pathway component expression. The significance of the difference between cGAS^+^ MN-associated slopes in neo-adjuvant treatment (NA Tx) pathological responder versus non-responder patients was tested using a linear interaction model and is reported at the bottom of the heatmap. Color maps to the Spearman correlation coefficient of associations (R_s_). P-values of correlations ≤0.10 are highlighted inside the boxes. Pairwise correlations with a p ≤0.05 are highlighted with a solid black border, whereas correlations with a p ≤0.10 are highlighted with a dashed black border. (**i**) Representative high-resolution images of CIN^high^ and CIN^low^ human EAC tumor biopsies specimens for both pathological NA Tx responders and non-responders, stained with DAPI (DNA), as well as anti-pan-cytokeratin (pCK), anti-cGAS and anti-STING antibodies. Images show a loss of malignant cell STING signal in CIN^high^ non-responders, but not CIN^high^ responders. Scale bars correspond to 50 μM. (**j**) Example of a CIN^high^ myeloid-dominated NA Tx non-responder tumor and a CIN^high^ myeloid- and T cell-enriched responder tumor. (**k**) Distribution of EAC tumor biopsies quartilized by CIN and degree of myeloid-enrichment (i.e. the extent of macrophage:T cell skew) across NA Tx responders and non-responders. (**l**) Distribution of EAC tumor biopsies quartilized by tumoral STING level and degree of myeloid-enrichment across NA Tx responders and non-responders. **** p ≤0.0001, *** p ≤0.001, ** p ≤0.01, * p ≤0.05.

